# Universality of distribution of tumor mutation burden - a biomarker for the tumor evolution and disease risk

**DOI:** 10.1101/2022.03.03.482775

**Authors:** Xin Li, Sumit Sinha, D. Thirumalai

## Abstract

Cancers, resulting in uncontrolled cell proliferation, are driven by accumulation of somatic mutations. Genome-wide sequencing has produced a catalogue of millions of somatic mutations, which contain the evolutionary history of the cancers. However, the connection between the mutation accumulation and disease development and risks is poorly understood. Here, we analyzed more than 1,200,000 mutations from 5,000 cancer patients with whole-exome sequencing, and discovered two novel signatures for 16 cancer types in The Cancer Genome Atlas (TCGA) database. A clock-like mutational process, a strong correlation between Tumor Mutation Burden (TMB) and the Patient Age at Diagnosis (PAD), is observed for cancers with low TMB (mean value less than 3 mutations per million base pairs) but is absent in cancers with high TMB. We also validate this finding using whole-genome sequencing data from more than 2,000 patients for 24 cancer types. Surprisingly, we discovered that the distribution of TMB are universal. At low TMB it exhibits a Gaussian distribution and transitions to a power law at hight TMB. The differences in cancer risk between the sexes are also mainly driven by the disparity in mutation burden. The TMB variations, imprinted at the chromosome level, also reflect accumulation of mutation clusters within small chromosome segments in high TMB cancers. By analyzing the characteristics of mutations based on multi-region sequencing, we found that a combination of TMB and intratumor heterogeneity could be a potential biomarker for predicting the patient survival and response to treatment.

## I. INTRODUCTION

Cancer is a complex disease caused by a gradual accumulation of somatic mutations, which occur through several distinct biological processes. Mutations could arise spontaneously during DNA replication^1^, through exogenous sources, such as ultraviolet radiation, smoking, virus inflammation and carcinogens^2^ or could be inherited^3^. During the past two decades, a vast amount of genomic data have been accumulated from patients with many cancers, thanks to developments in sequencing technology^4^. The distinct mutations contain the fingerprints of the cancer evolutionary history. The wealth of data has lead to the development of mathematical models^5–8^, which provide insights into biomarkers that are reasonable predictors of the efficacy of cancer therapies^9^. Our study is motivated by the following considerations:

*At what stage do most mutations accumulate?* The relation between mutation accumulation during cancer progression and disease risks in different tissue types and populations are not only poorly understood but also have resulted in contradictory conclusions. For instance, it has been suggested that half or more of the genetic mutations in cancer cells are acquired before tumor initiation^10, 11^. In contrast, analyses of colorectal cancer tumors^12^ suggest that most of the mutations accumulate during the late stages of clonal expansion. In the cancer phylogenetic tree, representing tumor evolution, most mutations would appear in the trunk in the former case^10^ while the branches would contain most of the mutations if the latter scenario holds^12^. Therefore, simple measures that discriminate between the two scenarios would be invaluable either for effective drug screening (given more targetable trunk mutations) or prediction of resistance (presence of more branched mutations)^13^.

*The determinants of cancer risk:* The number of mutations accumulated in cancer cells is likely related to great variations in cancer risks among different human tissue types^14^. According to the bad-luck theory^15^, cancer risk across tissues is strongly correlated with the total number of adult stem cell (ASC) divisions during a lifetime, with enhanced mutations accumulating in those that divide frequently. The implication is that some tissue types are at a greater disease risk than others. However, it is still unclear how mutation burden of cells is related to cancer risk among different tissues and genders.

*Biomarkers for patient survival and response to treatment:* With advances in understanding the immune response against cancer, immunotherapy is fast becoming a very promising treatment for metastatic cancer patients^16–18^. However, a large fraction of patients do not respond to immunotherapy positively, and suffer from the inflammatory side effects, and likely miss out on other alternative treatments^19, 20^. Therefore, finding biomarkers that could be used to separate patients who show response from those who do not^19, 21^ is crucial. The tumor mutation burden (TMB), although is an important biomarker^22, 23^ is not accurate, raising the need for other predictors^24^.

Motivated by the considerations described above, we developed computational methods to analyze the dynamics of accumulation of somatic mutations^15, 25^ during cancer progression through a systematic pan-cancer study. We used the publicly available whole-exome sequencing (WES) data produced by The Cancer Genome Atlas (TCGA) project^26^, together with the whole-genome sequencing (WGS) data recently provided by the Pan-Cancer Analysis of Whole Genomes (PCAWG) Consortium^27^, which contain a large and well-characterized patient cohorts from multiple cancer types. In recent years, mutations in many cancers^28, 29^ have been analyzed at the base pair level, to arrive at a number of signatures, which are combination of mutation types that drives different mutational process. A strong relation between the mutation load of two signatures (signatures 1 and 5) and the patient age is found in many cancers (even in high TMB cancers)^28, 30^. However, either only a small fraction (23%)^28, 30^ of the genetic mutations are considered or it is found as data from all cancer types are mixed^27^. Here, we developed a simple coarse-grained measure of the total mutation load, instead of a small fraction, and a theoretical model that explains the distinct role of mutations in different cancers. We arrive at four major conclusions. (1) In the 16 cancers (with WES data), we discovered two distinct signatures for the accumulation of somatic mutations. There is a strong correlation between the tumor mutation burden (TMB) and the patient age as long as the overall mutation load is below a critical value of ∼ 3 mutations/Mb. In sharp contrast, the correlation is weak for cancers with TMB that exceeds the critical value. Similar finding is observed for 24 cancers with WGS data. (2) Using a simple model, we show that if TMB is less than ∼ 3 mutations/Mb, the majority of the mutations appear before the tumor initiation^10, 11^. In the high TMB limit, most mutations accumulate during the tumor expansion stage, especially for some of the most lethal cancer types. (3) Although the mutation burden is one of the predominant factors that determines cancer risk, it only accounts for less than 50% of the differences in cancer risks among different tissues. Factors, such as the number of driver mutations required for each cancer, immunological and sex-hormone differences, also play a vital role in determining cancer risk. After excluding these factors, we show that the cancer risk disparity between the sexes is explained by the mutation load differences. (4) Our findings also show that a combination of TMB and intratumor heterogeneity could serve as a two-dimensional biomarker for predicting patient survival and response to treatment.

## II. RESULTS

### Low Tumor Mutation Burden (TMB) strongly correlates with Patient Age at Diagnosis (PAD)

We define TMB as the total number of non-synonymous and synonymous single nucleotide variants (SNVs) per megabase (Mb) of the genome. First, we consider 16 cancer types with WES data taken from the TCGA database^26, 31^ (see Table I in the Supplementary Information (SI)). Two mutation signatures emerge across these cancers. A strong positive correlation between TMB and PAD is found for cancers with low overall mutation load (TMB *<* 3 mutations/Mb). However, such a correlation is absent for cancers with high TMB. The distinct mutation signatures are a reflection of different underlying evolutionary dynamics among these distinct cancer types. We note parenthetically that the boundary between low and high TMB is somewhat arbitrary^32^. Interestingly, TMB ≈ 3 mutations/Mb, also provides a separation between cancers even at the chromosome level.

**TABLE I:**
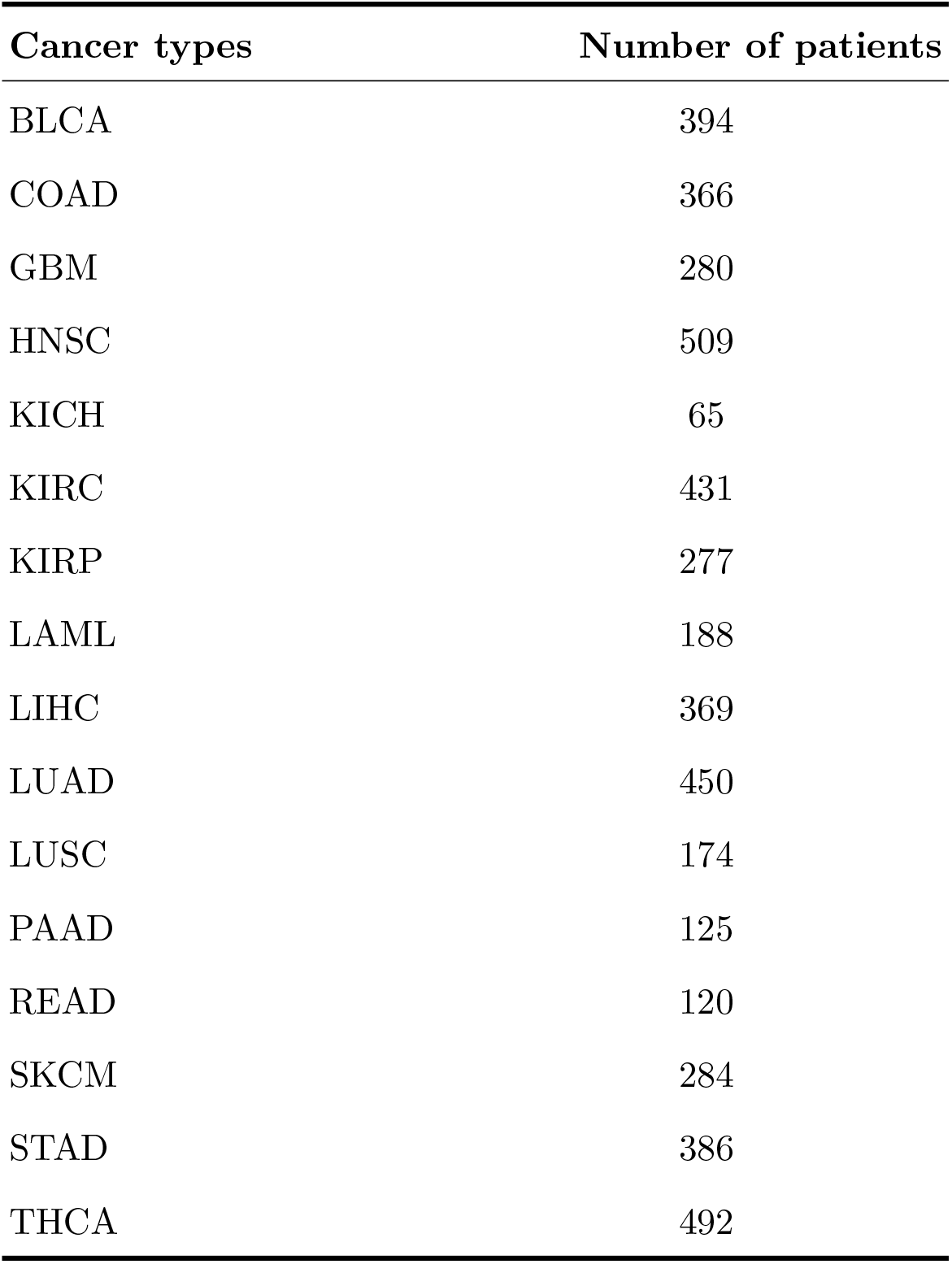
The number of patients for the 16 types of cancer taken from the TCGA database^31^. Cancer name abbreviations: BLCA – Bladder Urothelial Carcinoma; COAD – Colon Adenocarcinoma; GBM – Glioblastoma multiforme; HNSC – Head and Neck Squamous Cell Carcinoma; KICH – Kidney Chromophobe; KIRC – Kidney Renal Clear Cell Carcinoma; KIRP – Kidney Renal Papillary Cell Carcinoma; LAML – Acute Myeloid Leukemia; LIHC – Liver Hepatocellular Carcinoma; LUAD – Lung Adenocarcinoma; LUSC – Lung Squamous Cell Carcinoma; PAAD – Pancreatic Adenocarcinoma; READ – Rectum Adenocarcinoma; SKCM – Cutaneous Skin Melanoma; STAD – Stomach Adenocarcinoma; THCA – Thyroid Carcinoma.

Two key inferences can be drawn from Figs. 1A-F: (1) The linear behavior, for the six cancers in Fig. 1, between TMB and PAD reveals strong correlation with a Pearson correlation coefficient (*ρ*) varying between 0.66 to 0.93. The strong correlation (with a P value ≪ 0.001) is also evident directly from the violin plots shown in Fig. 1G (see also Fig. S1), which present the full mutation distribution for the cancers as a function of PAD. Note that if the overall mean TMB is less than 3 mutations/Mb, as is the case for LAML, THCA, KIRC, and KIRP cancers, the Pearson correlation coefficient is ≥ 0.85. However, the correlation is weaker as the TMB approaches and exceeds 3 mutations/Mb, as is the case for GBM and KICH (see Figs. 1E and 1F). The TMB in these cancers becomes higher as the PAD increases, which is similar to the behavior in normal tissues (see especially Fig. 1 here and Fig. 2b in^25^). (2) The mutation rates calculated from the slope of the solid lines in Figs. 1A-F vary from 13 (LAML) to 81 (KIRC) mutations per year for the whole genome. The range is in line with the mutation rate observed in normal tissues (40 to 65 mutations per year for normal colon, small intestine, liver and oesophageal epithelia cells)^12, 25, 33^. The calculated mutation rate, 13 mutations per year from Fig. 1A, for acute myeloid leukemia (LAML) coincides with the measured value (13±2) in previous studies^34^. Therefore, the cell mutation rate is approximately stable, and does not depend strongly on the tumor evolutionary process among the group of low TMB cancers (Fig. 1).

**FIG. 1:**
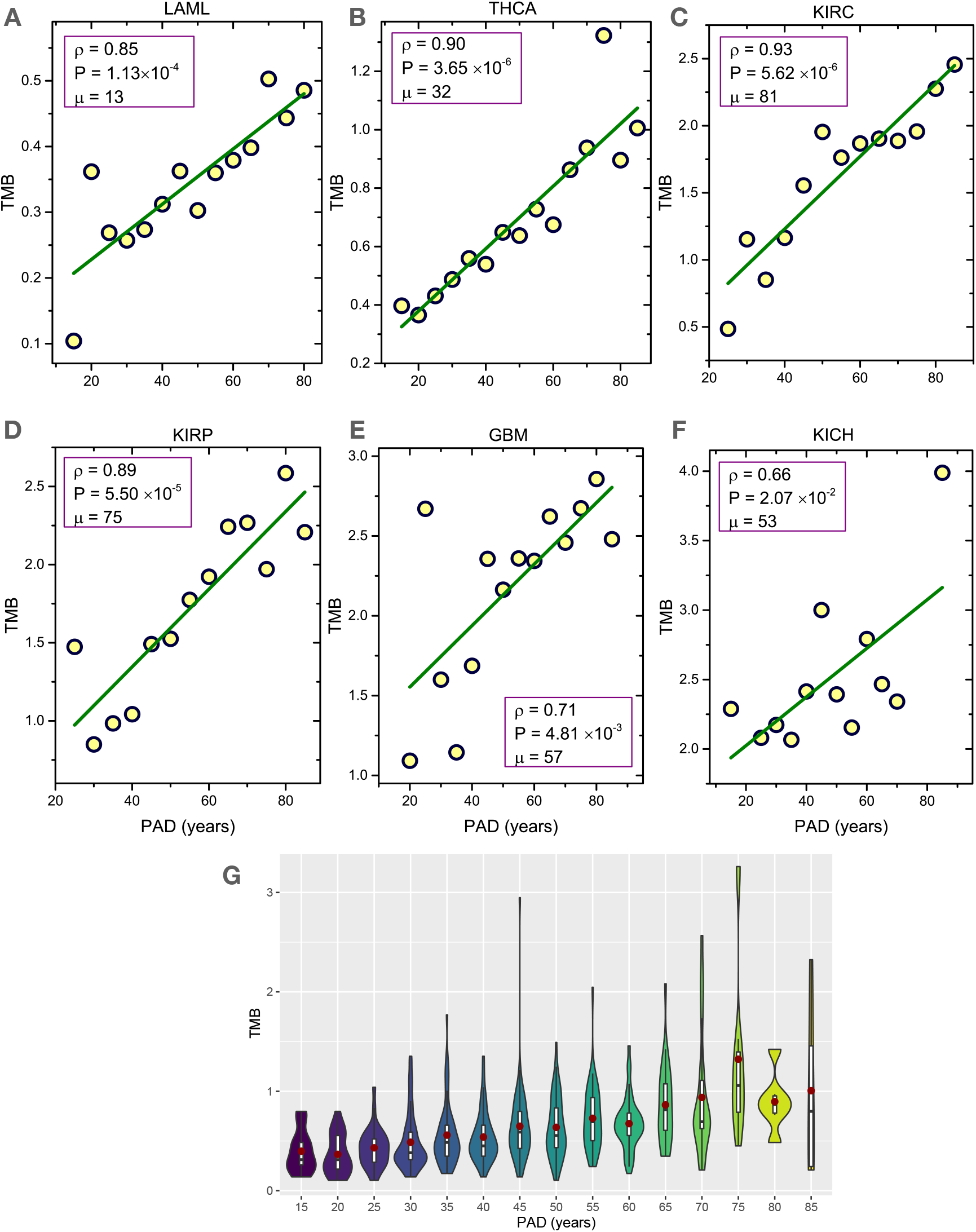
(A)-(F) The tumor mutation burden, TMB, (total number of non-synonymous and synonymous mutations per megabase) as a function of patient age at diagnosis (PAD). The mean value of TMB is used over a 5-year period. The green line shows the regression line. The Pearson correlation coefficient *ρ*, the significant P value from an F test and also the mutation rate *µ* are listed in the figures. The value of *µ* is derived from the slope of the green curve (0.0042 mutations/(Mb×year) for LAML in (a) for example, which leads to *µ* = 0.0042×3,000 ≈ 13 mutations/year for the whole genome). (G) Violin plots for the distribution of TMB as a function of PAD for THCA (see Fig. S1 for other cancers). Good correlation holds only if TMB is low. Data are taken from TCGA^26^.

**FIG. 2:**
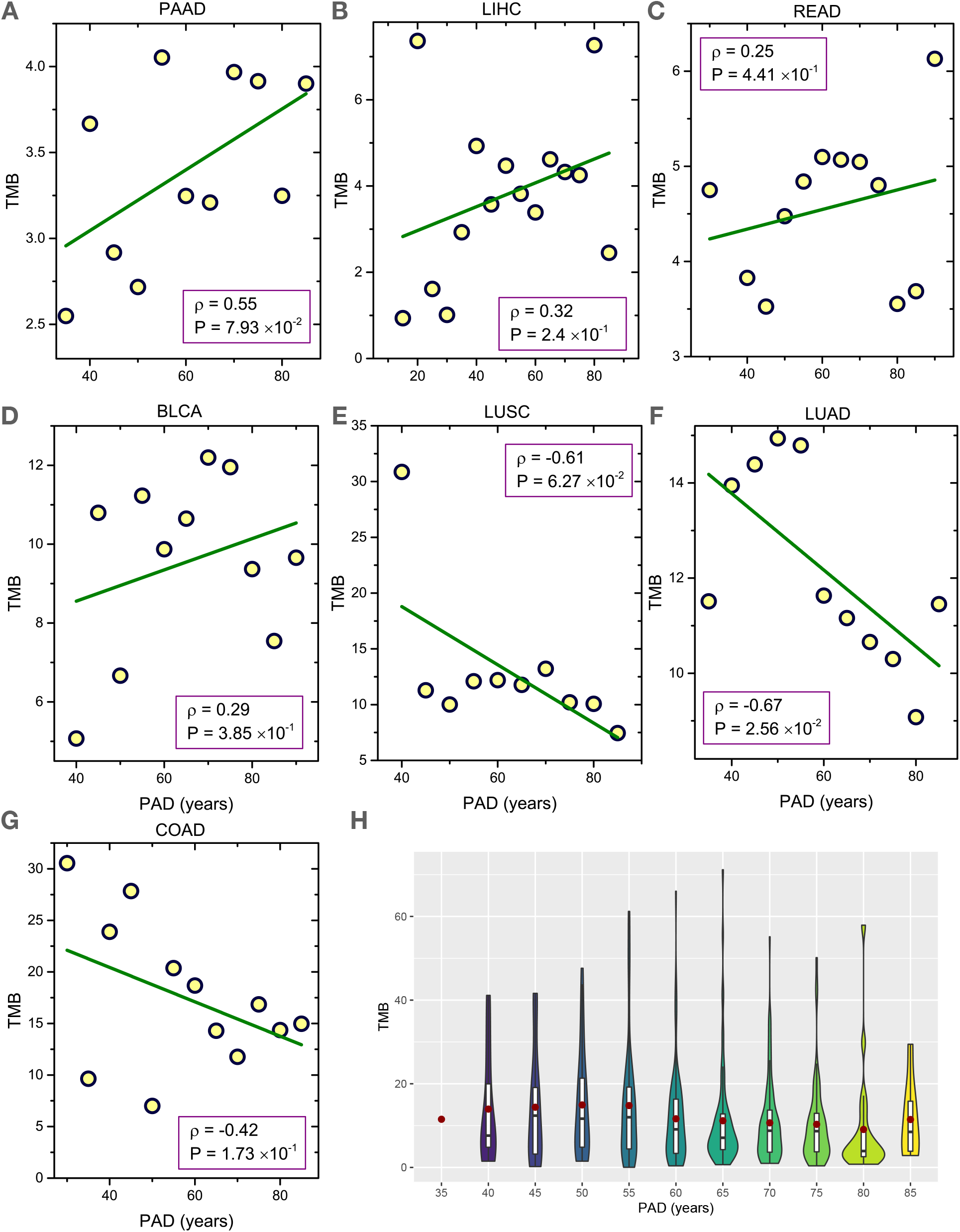
(A)-(G) Same as Fig.1 except the data are for cancers with high TMB. The green regression lines show a lack correlation between TMB and patient age. (H) Violin plots for the distribution of TMB as a function of PAD for LUAD (Fig. S2 provides data for other cancers).

### Large fraction of mutations accumulate before tumor initiation in low TMB cancers

The accumulation of mutations in cells of a tissue may be pictorially represented as a fish with the head, body and tail corresponding to the tissue development, self-renewal and tumor formation, respectively (see Fig. 1 in^10^). A prediction of the theory based on the “fish-like” picture is that the self-renewal stage (with a stable cell number) of a tissue is proportional to the patient age at tumor initiation. Therefore, a linear relation between the number of somatic mutations and the PAD must exist, assuming that a fixed time elapses from tumor initiation to diagnosis^10^. This important prediction is supported by our analyses for low TMB cancers, as shown in Fig. 1. Indeed, we can calculate the fraction of mutations accumulated in cells before tumor initiation once we know the average time it takes for tumor detection from the initiation time. Consider the thyroid carcinoma (THCA) as an example (Fig. 1B). The estimated latency period (*τ_L_*, from tumor initiation to diagnosis) is 5 to 20 years^35^. The number of mutations that accumulate in this time frame would be 150 to 600 (assuming an average accumulation rate of 30 mutations per year, see Fig. 1B). By noting that the TMB is around 0.7/Mb (Fig. 1B) at 46 years, the median age of THCA patient at diagnosis, and the human genome length 3,000Mb, the number of accumulated mutations at that age would be around 2100. Thus, for *τ_L_*=5 (20) years only 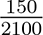 ≈ 7% 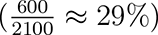 of the mutations occur during the period of tumor initiation till it is detected. This, implies that 93% (71%) of the mutations must have appeared before the initiation of the thyroid carcinoma for *τ_L_*=5 (20) years. In this case, the time from tumor initiation to detection is usually much shorter than the age of patients at diagnosis. From a similar analysis (see Table II), we conclude that the majority of mutations appear before the tumor initiation for cancers shown in Fig. 1. This is consistent with the results of other studies^10, 11, 34^. Take LAML as an example. By sequencing the M3-LAML genome with a known initiating event versus the normal karyotype M1-LAML genome and the exomes of hematopoietic stem/progenitor cells (HSPCs) from healthy people^34^, it was found that most of the mutations in LAML genomes are indeed random events that occurred in HSPCs before they acquired the initiating mutation. We also observe in Fig. 1 that the correlation between the TMB and patient age becomes weaker as the overall TMB increases. As noted above, the *ρ* value decreases from 0.9 to 0.7 as the overall mean value (averaged over all patients for different ages) of TMB increases from 0.4 mutations/Mb to 3.0 mutations/Mb.

### High TMB and PAD are weakly correlated

As the overall TMB exceeds 3 mutations/Mb, there is no meaningful correlation between TMB and PAD (Fig. 2 and Fig. S2). Interestingly, the results in Fig. 2 show that the TMB even decreases in certain cancer types as the PAD increases (negative correlation if one exists at all), see Figs. 2F-H as examples. This finding contradicts the conclusions reached in previous studies^10, 36^. Clearly, the findings for cancers with high TMB cannot be explained by the fish-like model because of the absence of a linear relation between the TMB and PAD, which holds only for cancers with low TMB (Fig. 1).

**TABLE II:**
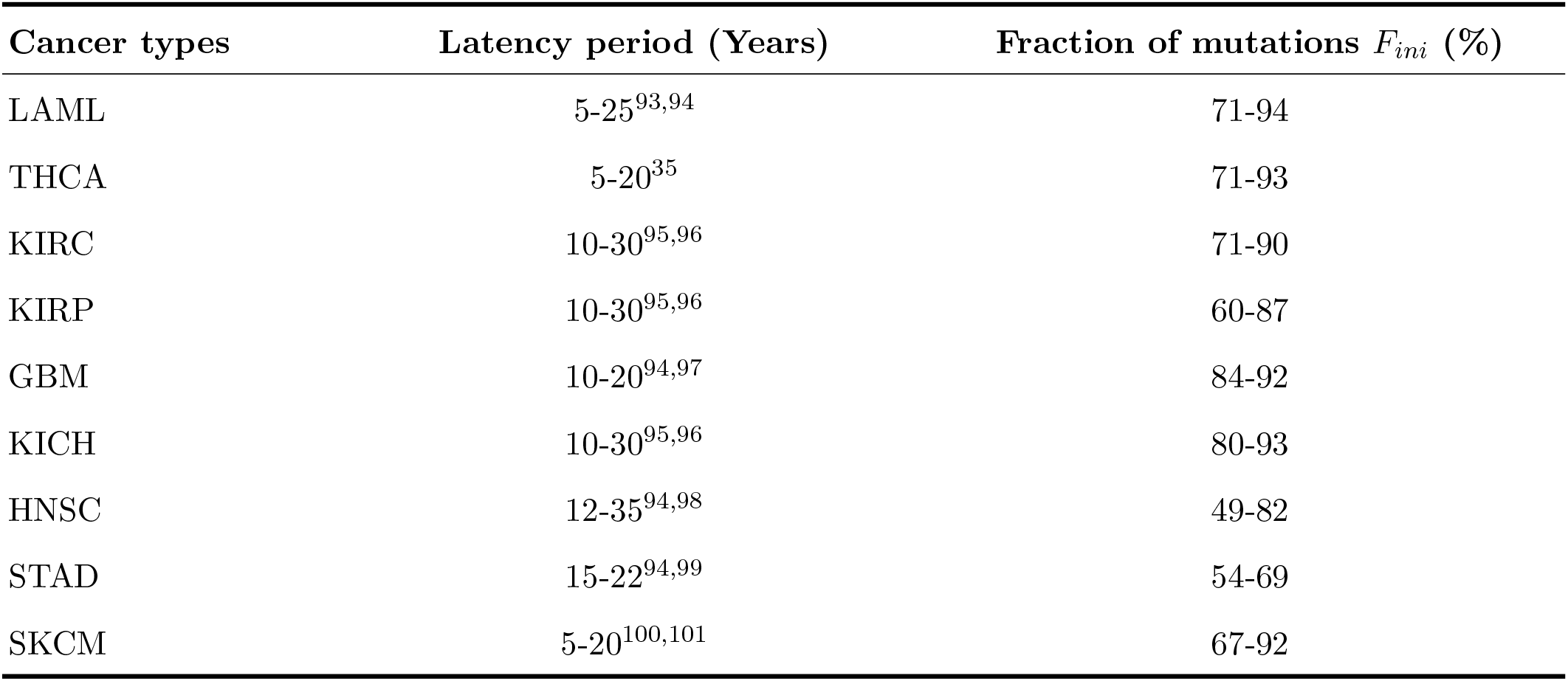
The latency period and fraction (*F_ini_*) of accumulated mutations before the initiation of tumors in Figs. 1 and 3.

### A simple model explains the TMB data

To understand the drastically different behavior between cancers with low and high TMBs in Figs. 1 and 2, we propose a simple theoretical model. It has been noted that the mutation rate can increase by a few orders of magnitude in tumor cells compared with normal cells caused by factors such as DNA mismatch repair deficiency or microsatellite instability^37, 38^. As a consequence, a very high TMB would be detected in the tumor cells at the time of diagnosis. Therefore, the number of accumulated mutations (AM) in cells during the tumor formation stage could be comparable or even larger than the number during the tissue self-renewal stage. Let *µ*_1_ be the mutation rate during normal DNA replication process, and Δ*µ*_1_, the enhanced mutation rate (relative to *µ*_1_) in tumor cells. The value of Δ*µ*_1_ would be zero if the mutation rate does not change during tumor progression. In this case, the number *N_t_* of AM in tumor cells is given by *N_t_* = *µ*_1_*T* where *T* is the age of the patient in years. The linear relation holds if the TMB is low (Fig. 1). However, the mutation rate increases in high TMB cancers (Fig. 2). Then, the number of mutations *N_t_* is given by,

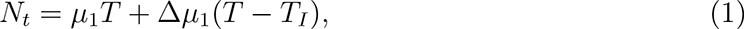

where the first term is the AM in the normal replication process, and the second term arises due to the accelerated mutations generated during the tumor formation stage. Note that *T_I_* corresponds to the time at tumor initiation. Because the latency period *T* − *T_I_* for tumors from the initiation till diagnosis is likely to be similar for patients with the same type of cancer^10^, the second term in Eq. (1) is likely to be a constant subject to only minor variations. If Δ*µ*_1 ≫_ *µ*_1_, then the first term in Eq. (1) is negligible, which leads to the weak dependence of TMB on the patient age as found in Fig. 2. Another potential mechanism for the finding in Fig. 2 could be that catastrophic events (such as chromoplexy, and chromothripsis) can lead to a burst of mutations that accumulate in tumors during a short period of time as observed in some cancers^39, 40^.

It is instructive to calculate the fraction of accumulated mutations before tumor initiation for a cancer with high TMB. For the hepatocellular carcinoma (LIHC) the median age of patients at diagnosis is 61, so the number *N_I_* = *µ*_1_*T_I_* of AM before tumor initiation is less than *µ*_1_*T* ≈ 3000 (with rate *µ*_1_ ≈ 50 mutations/year^12, 25^). From Fig. 2B, which shows that the TMB is ≈4 mutations/Mb at the same age and taking the genome length to be 3,000Mb, the total number of mutations accumulated at age 61 is about 12,000 (4 × 3000). Thus, the fraction of AM before tumor initiation should be less than 25%. We surmise that in cancers with high TMB (Fig. 2) most of the somatic mutations occur during the late stages of clonal expansion (see Table III). This conclusion agrees with the observations in colorectal cancers^12^, which allows us to conclude must be a cancer type with high TMB (*>* 3 mutations/Mb).

**TABLE III:**
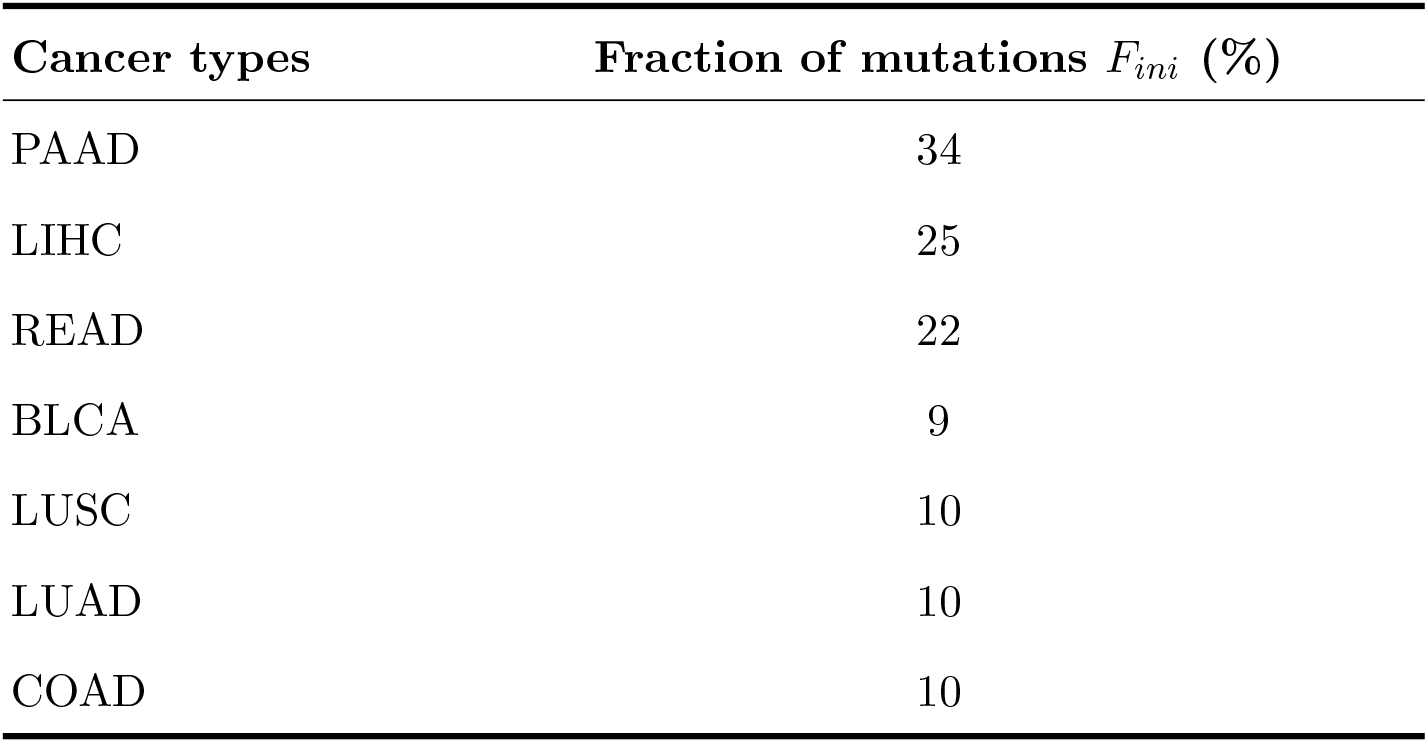
The upper bound for the fraction (*F_ini_*) of mutations that accumulate before the initiation of tumors in Fig. 2.

### Extrinsic Factors dominate mutation load in certain cancers

So far we have focused on somatic mutations arising from the intrinsic process due to DNA replication. The tissues of several cancers, such as head and neck, stomach, and skin, are constantly exposed to ultraviolet (UV) radiation and chemical carcinogens^41^. These factors directly increase the mutation rate (the “extrinsic origin”) of normal cells even in the absence of DNA replication during the cell lifetime^42^. To account for extrinsic factors, we modify Eq. (1) by adding an additional term *N_ext_*= *µ_ext_T*. As long as *µ_ext_ ≫*Δ*µ*_1_, a linear relation between TMB and patient age holds (see Fig. 3A-D and Fig. S3). In these cases, just as in Fig. 1, more mutations appear at the trunk of tumor phylogenetic trees types (see Table II and Fig. 3E). The strong linear correlation, with high Pearson correlation coefficient in Fig. 3A-C, shows that in these cancers *µ_ext_* far exceeds *µ*_1_ and Δ*µ*_1_. It might be relatively easy for the immune system or drugs to clear cancer cells from patients with such cancers because a large number of mutations are shared among all the cancer cells^13^.

**FIG. 3:**
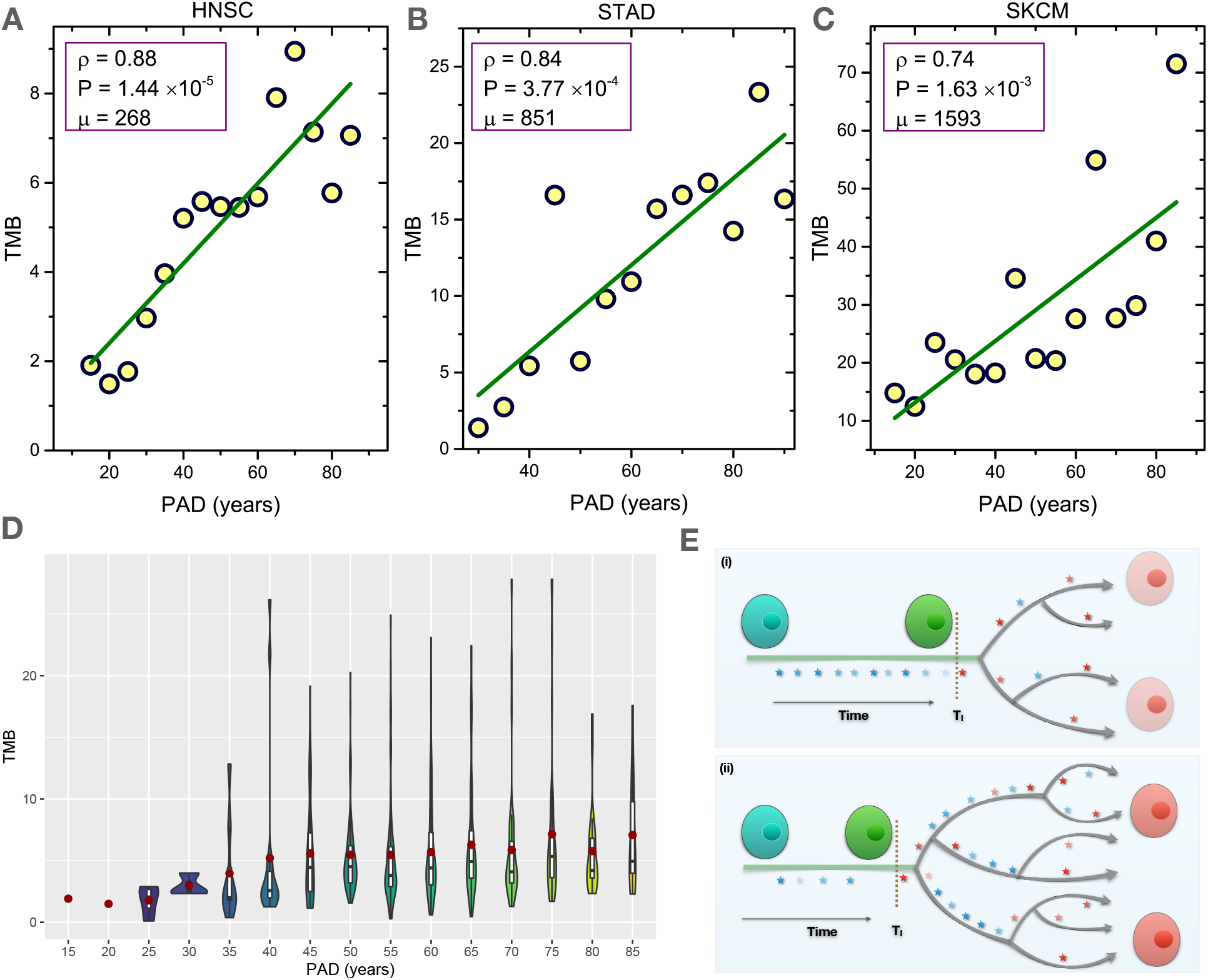
(A)-(C) TMB as a function of age for cancers strongly influenced by environments. The regression lines are in green. The Pearson correlation coefficient *ρ*, the significant P value, and the mutation rate *µ* (number of mutations per year for the whole genome) are shown in the figures. (D) Violin plots for the distribution of TMB as a function of PAD for HNSC (see Fig. S3 for other cancers). (E) Schematic of the mutation accumulation in cells during cancer progression. (i) corresponds to the case in Fig. 1 and Fig. 3(a)-(c), and (ii) shows the case in Fig. 2. Mutations are indicated by stars, and *T_I_* is the tumor initiation time. Normal cells are in blue/green colors without/with neutral mutations (blue stars) while cancer cells are in red color with driver mutations (red stars).

In contrast, the majority of the mutations are expected to be present in low frequencies and appear at the branches and leaves of the tumor phylogenetic tree (see Fig. 3E) for the cancers shown in Fig. 2. This results in immune surveillance escape and drug resistance^12^. In support of this finding, we found that the most deadly cancers (ones with the lowest 5-year survival rate), such as pancreatic cancer (8.5%), liver hepatocellular carcinoma (17.7%), and lung cancer (18.6%)^43^ (see Fig. S4), appear in Fig. 2.

### Relation of TMB and PAD from whole-genome sequencing

Because a strong linear relation is observed (see Fig. S5) between the number of SNVs for the whole genome and that for the whole exome of cancer patients, we expect similar correlation to be borne out by analyzing the WGS data. Using the recently available whole-genome sequencing data provided by the ICGC/TCGA Pan-Cancer Analysis of Whole Genomes (PCAWG) Consortium^27^, we performed a similar analysis for 24 cancer types (see Fig. 4). Due to the limited number of samples for each cancer type (see Table VII), we plotted all the patient data without binning into age groups. We found a strong correlation between TMB and PAD for cancer types with low mutation burden (see Fig. 4A-K and Table VII). However, such a correlation is absent for cancer types with high mutation burden (see Fig. 4L-X), which agrees well with our findings from the analysis of the WES data. For those cancers (Fig. 4R, S, X), under a high mutation burden but strongly influenced by extrinsic factors, a weak correlation is observed which might be due to the limited number of samples. A relatively strong correlation emerges for HSCC (head and neck cancer) compared to STAD (Stomach Adenocarcinoma) given a larger sample size for the former (see Fig. 4R and S). For MELA (Skin Melanoma), a very larger sample might be needed to observe the correlation due to substantial variations in the mutation burden. Therefore, the findings extracted from the WES data is robust, and is independent of the type of database we used.

**FIG. 4:**
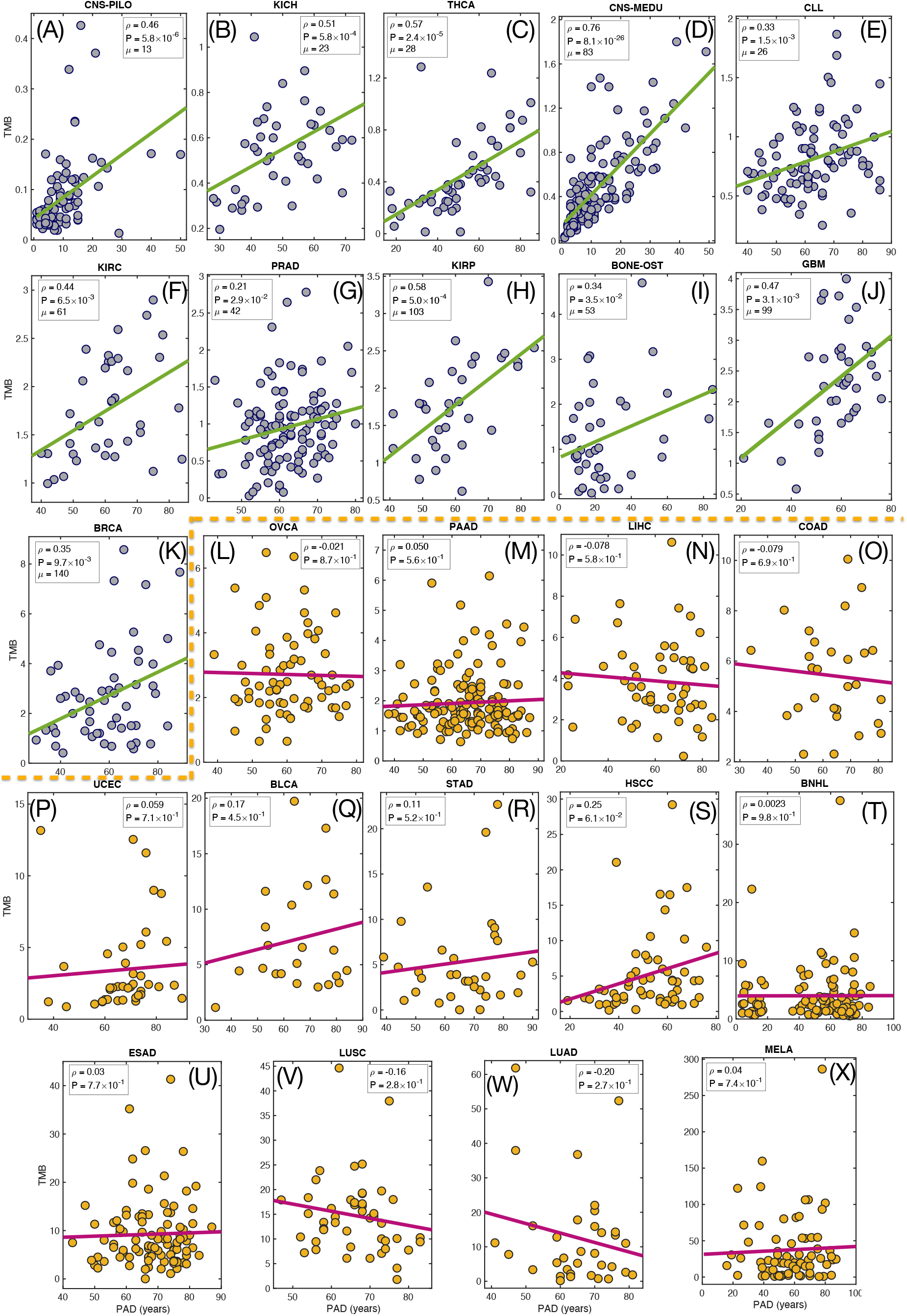
Tumor mutation burden, TMB, (total number of non-synonymous and synonymous mutations per megabase) as a function of patient age at diagnosis (PAD) for 24 types of cancers from the whole-genome sequencing^27^. Each filled circle represents one patient data. The green/magenta line shows the regression line. The Pearson correlation coefficient *ρ*, the significant P value from an F test and also the mutation rate *µ* (number of mutations per year for the whole genome in Figs. 4A-K) are listed in the figures. Good correlation holds only if TMB is low which confirms the findings in Figs. 1 and 2 obtained by analyzing the whole-exome sequencing data.

### Two universal patterns in cancers with low and high TMB

Our analyses above show two distinct signatures for TMB across many cancers. In order to elucidate if there are underlying universal behaviors in the cancers demarcated by the TMB, we first rescaled the TMB based on linear regression over the PAD for cancers listed in Figs. 1 and 3. Most interestingly, the data from the nine cancers fall on a linear curve, TM̂B ≡ (TMB− *b*_0_)*/a*_0_ = PAD (see Fig. 5A), which clearly demonstrates a clock-like mutational process (with a constant mutation rate) in cancer evolution^30^. For the remaining cancers in Fig. 2, we rescaled the TMB only by the mean value *µ*_0_ because of the absence of a linear relationship. Surprisingly, the data from these cancers also fall into a straight line but with a nearly *zero* slope (see the Pearson correlation coefficient *ρ*, P-value and the green line in Fig. 5B), which supports the absence of *any* correlation between TMB and PAD for these cancer types.

**FIG. 5:**
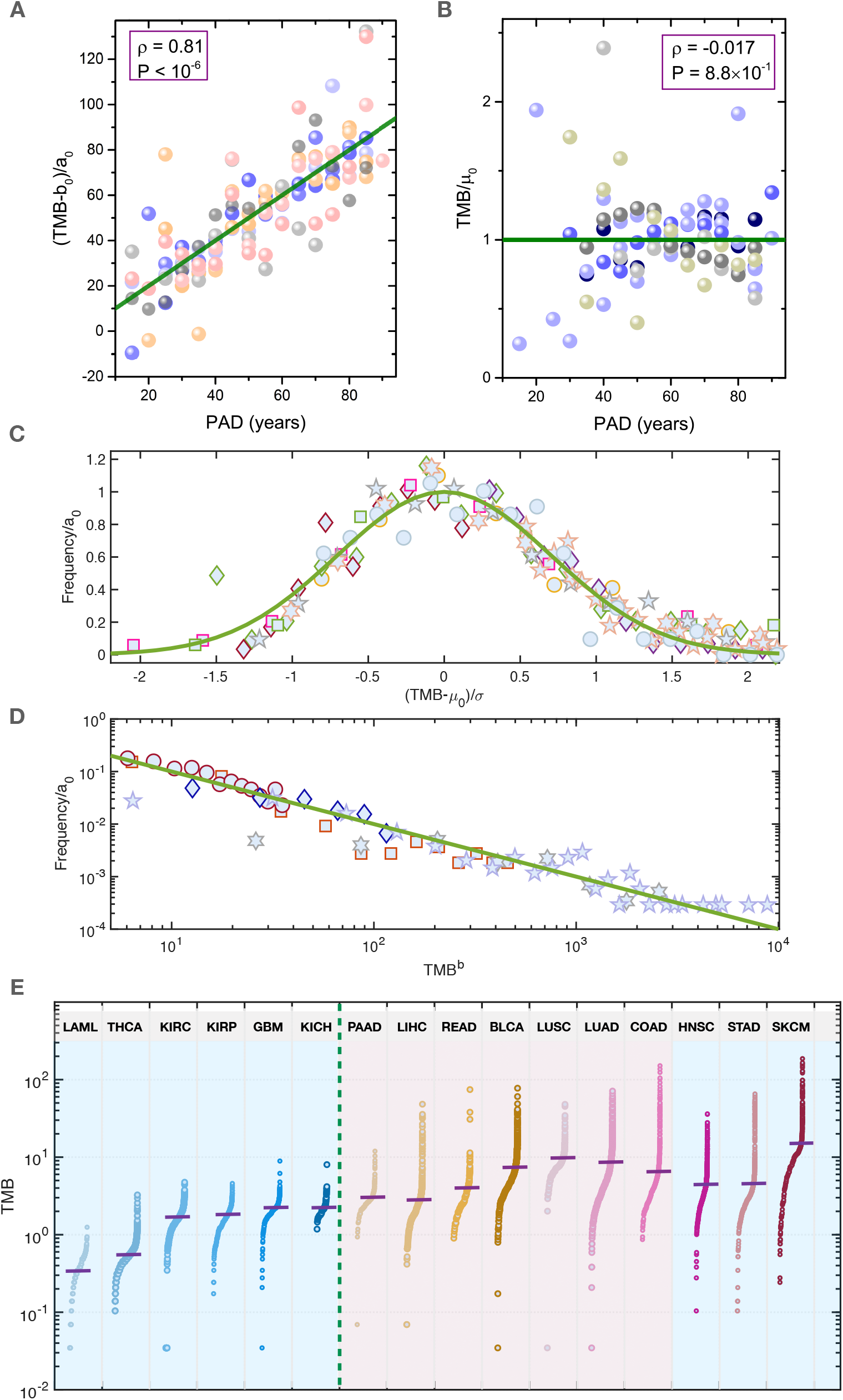
The rescaled TMB versus PAD for the 16 cancer types. (A) All nine types of cancer in Figs. 1 and 3 are displayed together in (A) after the TMB is rescaled by the slope (*a*_0_) and intercept (*b*_0_) of the corresponding linear regression lines. (B) The other seven types of cancers (Fig. 2) with TMB rescaled by the mean TMB value (*µ*_0_) of each cancer. The green line is described by *y* = *x* in (A) and *y* = 1 in (B). The Pearson correlation coefficient *ρ*, and the significant P value are given in the figure. (C) Rescaling the TMB distribution for each cancer type gives a Gaussian distribution (with the mean value *µ*_0_, variance *σ*, and coefficient *a*_0_) fits the data for cancers in Figs. 1 and 3. The rescaled TMB distributions for these cancers collapse into the standard normal distribution (see the green line). (D) Same procedure as in (C) leads to a fat tail power-law distribution (*P* (TMB) = *a*_0_ × TMB*^b^*) for data analyzed in Fig. 2. The green linear line has a slope value of −1. Each symbol represents one type of cancer. (E) The TMB across the 16 types of cancer. Each dot represents the TMB of a single cancer patient. The purple solid lines mark the median TMB value for each cancer. The green dashed line shows the boundary between cancers with low and high TMB defined in Figs. 1 and 2. The three cancer types discussed in Fig. 3 are listed at the end.

Next, we investigated the TMB distribution for each cancer type. Surprisingly, we found two universal TMB distribution patterns. A Gaussian distribution is found for all cancers analyzed in Figs. 1 and 3 (see Fig. S6). Remarkably, after rescaling, the TMB distributions from the nine cancer types collapse onto a single master curve described by the normal distribution (see Fig. 5C). The universal TMB distribution found in Fig. 5C for the nine cancers is related to the Gaussian distribution of PAD (see Figs. S17 and S18), and the linear relation between TMB and PAD (see Figs. 1 and 3). By simulating a population of patients with such properties, we obtain a Gaussian distribution for TMB (see Fig. S8 and Materials and Methods). We find it remarkable that for tumor evolution, which is a complex process, essentially a *single easily calculable parameter (TMB) from database* explains a vast amount of data for low TMB cancers.

In contrast, the distribution of cancers in Fig. 2 with large TMB is governed by a power-law (see Fig. S7 and Fig. 5D), except for PAAD (with TMB close to the critical value), and LUSC which have a Gaussian distribution. In these cancers, majority of the mutations accumulate after tumor initiation and the dynamics is likely to be highly complicated with many interrelated mutational processes contributing to clonal expansion. Similar signatures are found for cancers with low and high TMB, respectively, after we separate the patient data into two groups according to sex (see Figs. S9-S11). The entirely different distributions of TMB found for the two groups of cancers show that cancers with high and low TMB follow distinct evolutionary processes.

Taken together, we discovered two distinct universal signatures for TMB across many cancers, which could be understood in terms of the mean/median TMB (∼ 3 mutations/Mb, see the green dashed line in Fig. 5E as the boundary and Figs. S12 and S13 for further support). The linear relation between TMB and PAD, and the universal Gaussian distribution of TMB in the first group of cancers imply a clock-like mutation dynamics holds^30^. In contrast, the power-law distribution of TMB and lack of linear relation between TMB and PAD suggest a more complex mutation dynamics in the other group of cancers.

### Imprints of TMB in chromosomes

To illustrate the different mutation signatures between cancers with low and high TMB further, we investigate the mutation profile of each chromosome^44^ using LAML from Fig. 1 and LIHC from Fig. 2 as two examples. The mutation frequency is much higher along each chromosome (at 10 Mb resolution) in the high TMB LIHC compared to the low TMB LAML (see Figs. 6A and 6B) except in the region around the centromere where a very low mutation frequency is observed in many chromosomes. From this coarse-grained description, a non-uniform distribution is observed clearly for the mutation profile in LIHC while it is rather uniform with only small deviations in LAML.

**FIG. 6:**
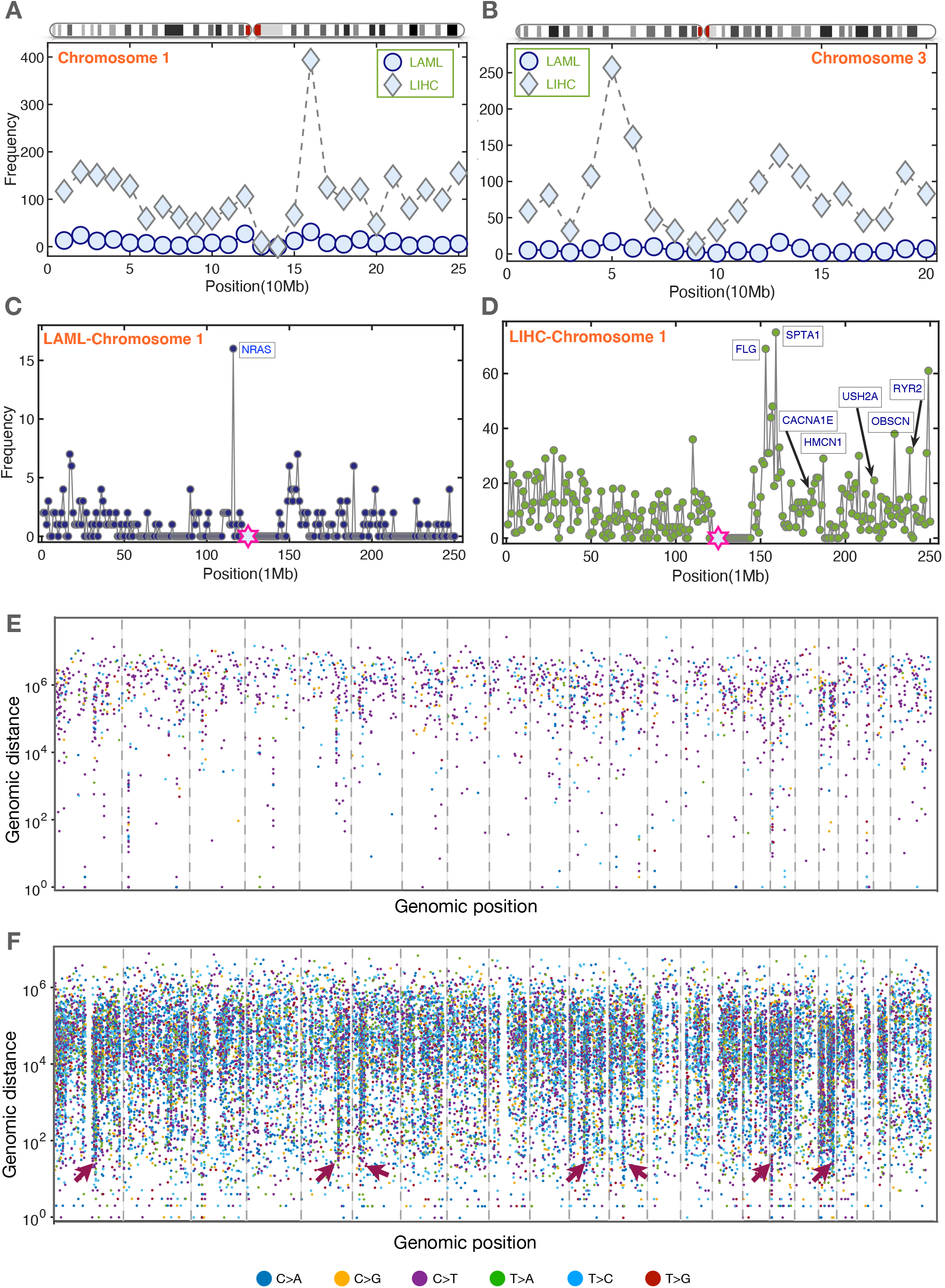
Tumor mutation profiles along chromosomes in LAML (low TMB) and LIHC (high TMB). (A) and (B): Number of mutations per 10 Mb across chromosomes 1 and 3 for LAML (197 patients) and LIHC (198 patients). The chromosome ideograms are also listed on top of each figure. (C) and (D): Number of mutations per 1 Mb along chromosomes 1 for LAML and LIHC. The genes (labelled in blue) mutated at a high frequency are also listed in the figures. The hexagram in magenta indicates the centromere location on each chromosome. (E) and (F): Rainfall plots for LAML and LIHC. The somatic mutations represented by small dots are ordered on the x axis in line with their positions in the human genome. The vertical value for each mutation is given by the genomic distance from the previous mutation. The colors of the dots represent different classes of base substitution, as demonstrated at the bottom of the figure. The red arrowheads indicate hypermutation regions. The 22 autosome and X chromosome are separated by the dashed lines.

Next, we consider a higher resolution (1Mb) mutation profiles (see Figs. 6C-D and Figs. S14 and S15) to capture more nuanced differences between these two cancers. Although the mutation frequency profiles are uniform in chromosomes 1 and 3 in LAML, occasionally, deviation from the mean value is detected (see the peak labeled with the name of a mutated gene in Fig. 6C and Fig. S14) as a consequence of strong selection of driver mutations^45^. For the high TMB cancer, LIHC, a much more complex landscape is observed in the mutation profile. In addition to great variations of mutation frequency from one region to another, more highly mutated genes are detected and are often found to appear in a very close region along the genome (see Fig. 6D and Fig. S15). This has been found in other high TMB cancers such as PAAD and LUAD^28^.

We then created the rainfall plot, which is frequently used to capture mutation clusters in chromosome^28^, to describe the different mutation characteristics for low and high TMB cancers (see Figs. 6E-F). The genomic distance between two nearby mutations is around 10^5^ - 10^6^*bp* for LAML. There is no clear hypermutation region for the 23 chromosomes (see Fig. 6E and Fig. S16) in this cancer type, which is consistent with the mutation profile illustrated in Figs. 6A and 6C and the simple Gaussian distribution found in Fig. 5C for low TMB cancers. In contrast, many hypermutation regions (with intermutation distance *<* 10^4^*bp*) are present in LIHC (see the red arrowheads in Fig. 6F and Fig. S17 for the detailed signatures in chromosome 1). Such a non-trivial mutation profile for high TMB cancers provides a hint for the power-law TMB distribution found in Fig. 5D. Similar profiles are also found for the kataegis patterns based on the WGS data (see Fig. S18). These findings suggest that, on all scales, the evolutionary dynamics is much more complicated in high TMB cancers compared to low TMB cancers.

### Cancer risk and TMB

It is thought that the continuous accumulation of mutations drives the transformation of normal cells to cancerous cells. However, there is a controversy on the origin of the great variations in cancer risk among different tissues. A strong correlation is found between cancer risk in different tissues and the total number of adult stem cell divisions^15^. Therefore, the variations of cancer risk might be mainly determined by the mutation burden in the tissues, which seems to be consistent with the extreme variations of TMB observed in different cancer types (see Fig. 5E). We now assess the correlation between the cancer risk and the mutation burden in cancer patients directly by taking data for the 16 cancers considered above from the TCGA and SEER database (see Tables IV, V and the Materials and Methods). The cancer risk for males and females vary greatly (see Table V), which we discuss further in the following sections. Here, we include the data for both the sexes. The median age for all cancer patients regardless of sex differences at diagnosis is around 66 years^14^. In order to rationalize the age disparity in different cancers for both the sexes (see Figs. S19-S20, and Table VI), we adjusted the value of TMB for both the sexes. We assumed that both the sexes accumulate mutations in a similar way (see Figs. S21-S23 and Materials and Methods for validation). We investigated the relation between cancer risk and TMB (see Fig. S24 (a) and (b)). The Pearson correlation coefficient *ρ* is 0.6 with a P value of 2.67 × 10^−4^ (*ρ* = 0.7 with *P* = 2.70 × 10^−5^) between cancer risk and TMB for 16 (15) cancers. Additional results are shown in Fig. S24 (c) and (d) without age-adjustment. A similar Pearson correlation coefficient (*ρ* = 0.66) is also found between cancer risk and mutation frequency across 41 cancers (using double-logarithmic coordinates with the data from a uniform sequencing pipeline) in a recent study^46^. Therefore, the cancer risk and TMB are correlated.

**TABLE IV:**
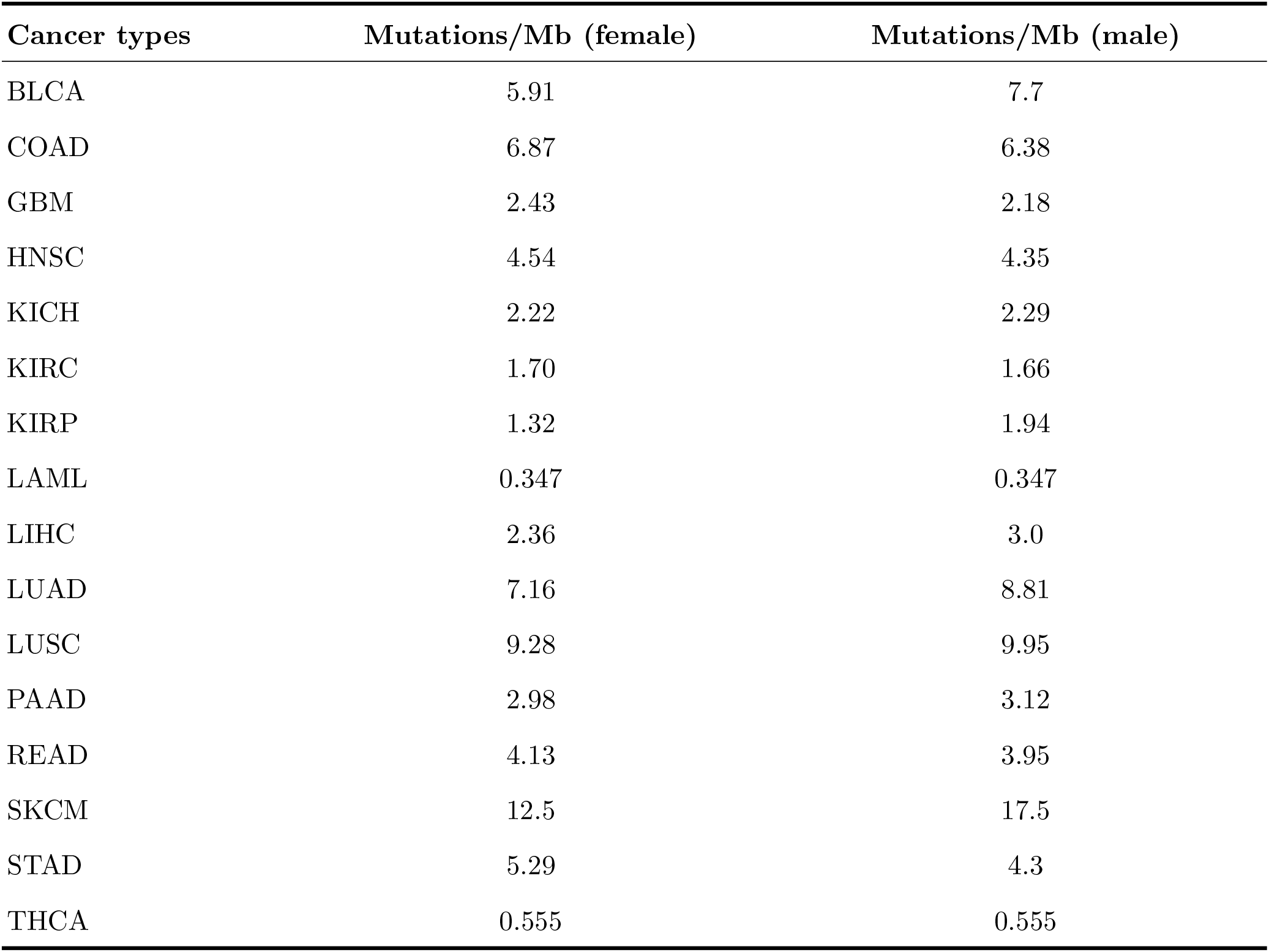
Mutation burden (median value) for both females and males across the 16 types of cancer from TCGA database^31^.

**TABLE V:**
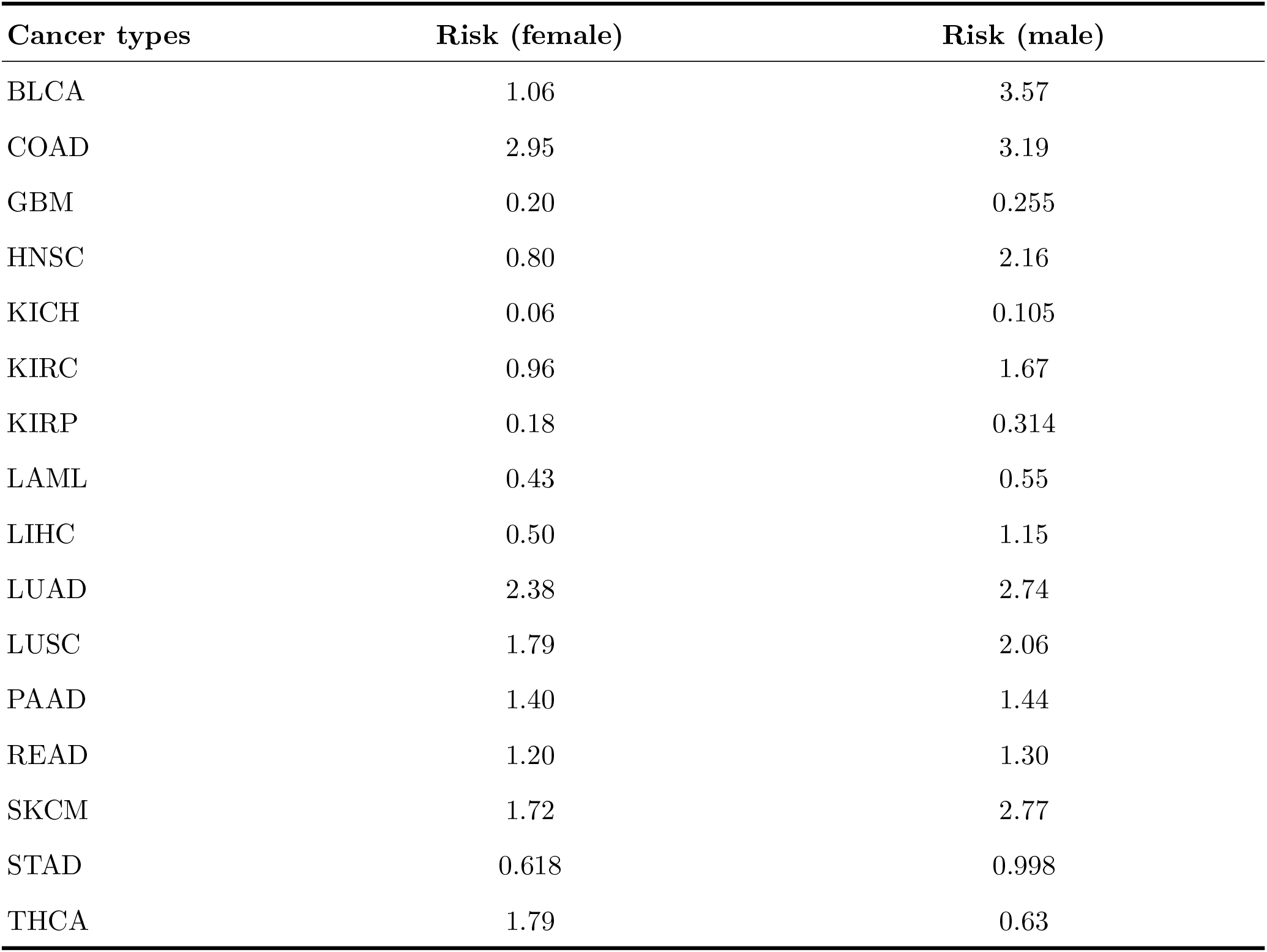
Cancer risks (%) for both female and male populations across the 16 types of cancer (see the analysis above in the Supplementary Information).

Because TMB accounts for less than 50% (≈ 0.7^2^) of differences in cancer risks among different tissues, other factors must be taken into account to explain quantitatively cancer risks among all tissues. These include the driver mutation number^47^, immunological^48^ and sex-hormone differences^49^. Nevertheless, the mutation burden is one of the predominant factors for predicting cancer risks across different tissues. If we only compare cancer risks of the same type for both the sexes, excluding gender-specific cancers (with 16 types of cancers considered here as shown in Fig. 1-3), we might be able to exclude the influences of these factors on cancer incidence, which would allow us to focus only on TMB.

### TMB explains the role of sex in cancer risk

The risk ratio for the male to female changes from 0.39 (thyroid) to 28.73 (Kaposi sarcoma)^50^. For most types of cancer, men have a higher risk compared to women. Although breast and prostate cancers are gender-specific, driven by sex steroid hormones^49^, the enhanced risk in men is still a puzzle for many other non-gender-specific cancers. There are many potential factors leading to the disparity. These include but are not limited to genetic and expression variation^51^, sex hormones^52^, environmental exposures^53^.

We investigated the relation between cancer risk and the mutation burden for both the sexes for each cancer separately (see Figs. 7A-B). Surprisingly, the cancer risk can be explained solely by the TMB score in 13 out of the 16 cancers. Nine of them, which show high cancer risk, also have high TMB score (HNSC, KIRC, KIRP, LIHC, THCA, BLCA, LUAD, LUSC, and SKCM), and four of them show almost the same risk and have similar TMB (GBM, LAML, PAAD, and READ). The negative correlation between cancer risk and TMB in KICH in Fig. 7 might be because the sample size (66 in total) is small. In this case there is a large deviation in the median age of patients between men and women populations (see Fig. S20g). The lower risk, but higher TMB in women in STAD and COAD, could be caused by other factors such as the immune response and sex-hormone. Inflammation is frequently observed in these two cancer types, which results in the activation of the immune system^54^. Because the efficiency of the immune system declines faster in men^48^, and estrogen suppresses tumor growth in female patients with COAD^55^, they might decrease the cancer risk in women with STAD and COAD.

**FIG. 7:**
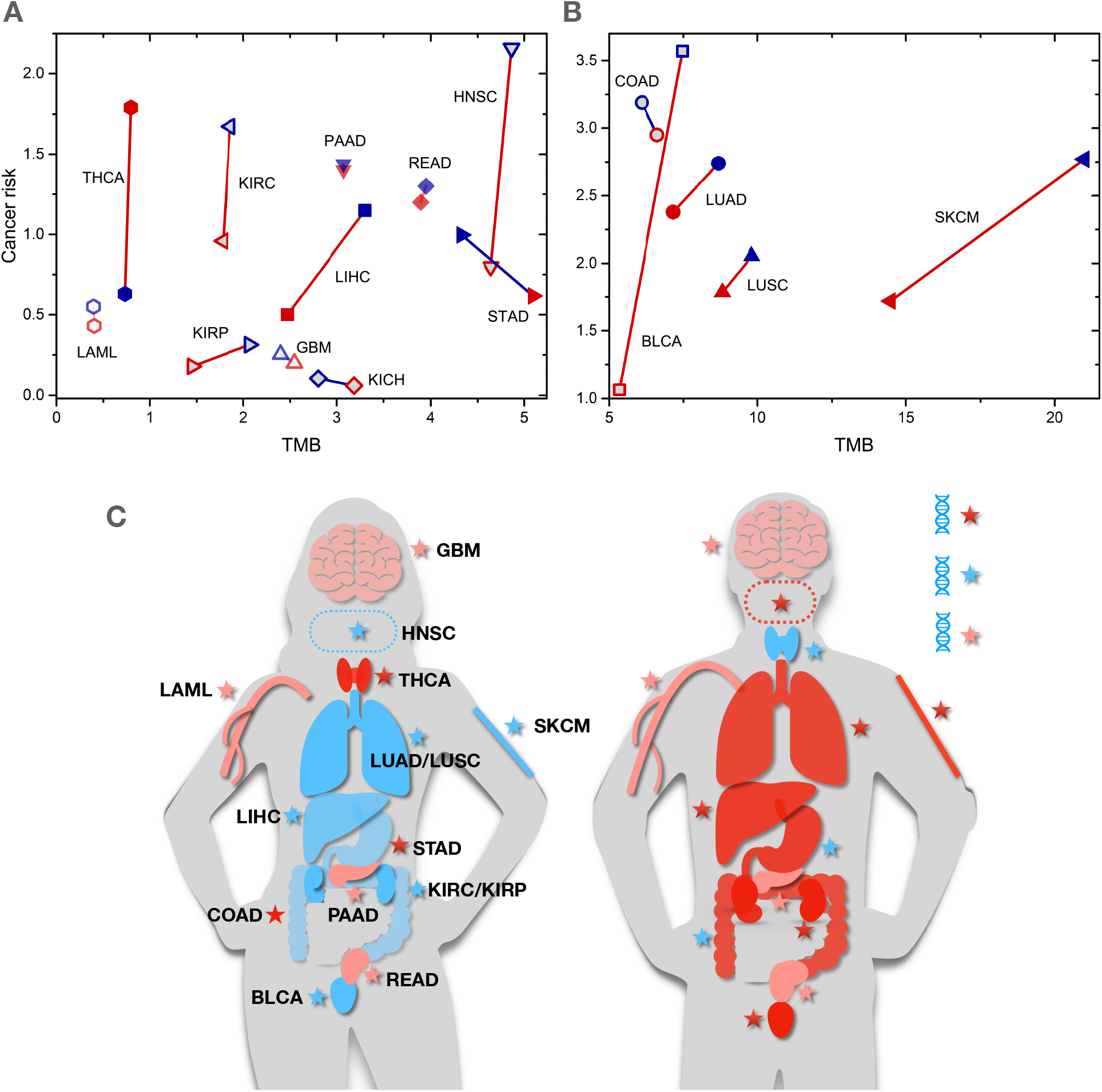
The relationship between lifetime cancer risk (%) and TMB for both the sexes. (A)-(B) The data are the same as in Fig. S24 (a) and (b), while the solid lines (red for a positive, navy for a negative correlation) connect the data for both the sexes for the same cancer type. The data are separated into two groups for better visualization. Red (navy) symbols show the data for females (males). (C) Schematic of the relative cancer risks and the TMB for females and males. The organs in red, blue, and pink colors indicate a higher, lower and similar cancer risk (relative to the other gender), respectively. Similarly, the red, blue and pink stars show a higher, lower and similar TMB in the tissue accordingly. For clarity, we just use three colors to illustrate the relative values of these two quantities between the two genders. The exact values are shown in (A)-(B).

Taken together our analyses show that the risk disparity between men and women for the same type of cancer can be explained mainly through the mutation burden in the cancer cells (see the schematic illustration in Fig. 7C). However, other factors on cancers have to be considered to explain the risk among different tissues as shown in Fig. S24 and in previous studies^15, 46^. The number (*N_d_*) of driver mutations required for cancer emergence is an important factor^47^. Because it takes a long time for a normal cell to accumulate driver mutations^56^, making the simplest assumption of independent driver mutation events, the cancer risk for tissues with a higher threshold of *N_d_* would be reduced. Consider THCA for instance, it is thought that a single driver mutation is required for the transition from a normal to a cancerous cell^47^. This might explain the relatively higher risk (see Fig. 7A) of THCA with low TMB compared to other cancers with lower risk but higher mutation burden, such as GBM which might require 6 driver mutations^47^. Although the mutation burden for SKCM is much higher than other cancers such as LUAD (HNSC), the large number of drivers (11) required for SKCM emergence could be the critical factor resulting in a similar risk as LUAD (HNSC), which might only require 4 (5) driver mutations (see Fig. 7B)^47^.

### Influence of TMB and intratumor heterogeneity (ITH) on patient survival

We have shown that high TMB is frequently related to high cancer risk for both the sexes for the same type of cancer. Although not desirable, high TMB could also elicit enhanced expression of a large number of neoantigens in cancer cells, which could be recognized by the T-cells for apoptosis and affect patient survival^57^. However, cancer cells have evolved different escape pathways, and could even disarm immune surveillance^58^. As we gain more knowledge about the mechanisms associated with these critical processes, immunotherapy may be modified to tame many metastatic cancers^16^ regardless of the TMB. At present, TMB is used as a potential biomarker for immunotherapy. Indeed, pembrolizumab has just been approved by FDA for the treatment of adult and pediatric patients with unresectable or metastatic tumor with high TMB based on cohorts with ten tumor types from the phase 2 KEYNOTE-158 study^59^. However, TMB alone is not a perfect predictor for patient response and survival^24^.

As an example, we investigated the influence of TMB, and patient age on patient response to immunotherapy (see Fig. S26 and Materials and Methods). Surprisingly, we found a much higher fraction of older patients, compared to younger ones, showed favorable response given a similar mutation burden level, which cannot be explained if only TMB is used as a predictor of the efficacy of treatment. We explained the data using a theoretical model based on the dynamics of mutation accumulation. We propose that older patients would have accumulated more clonal mutations under the same TMB level, which could be the underlying mechanism for the improved response.

To learn how the clonal characteristics of mutations influence patient survival further, we utilized the multi-region sequencing data which distinguishes the clonal from subclonal mutations better than single-region sequencing data^60^. We found that patients (lung adenocarcinoma), with a high clonal tumor mutation burden (cTMB, number of clonal mutations), show a better survival probability compared to those with a low cTMB (see Fig. 8A), thus supporting our conjecture above. However, the difference is not substantial which might due to the limited patient samples and short clinical trial duration. In addition, we also found that the patients with a low intratumor heterogeneity (ITH, defined based on the number of subclones^60^) tend to have a better survival probability compared to those with a high ITH (see Fig. 8B).

**FIG. 8:**
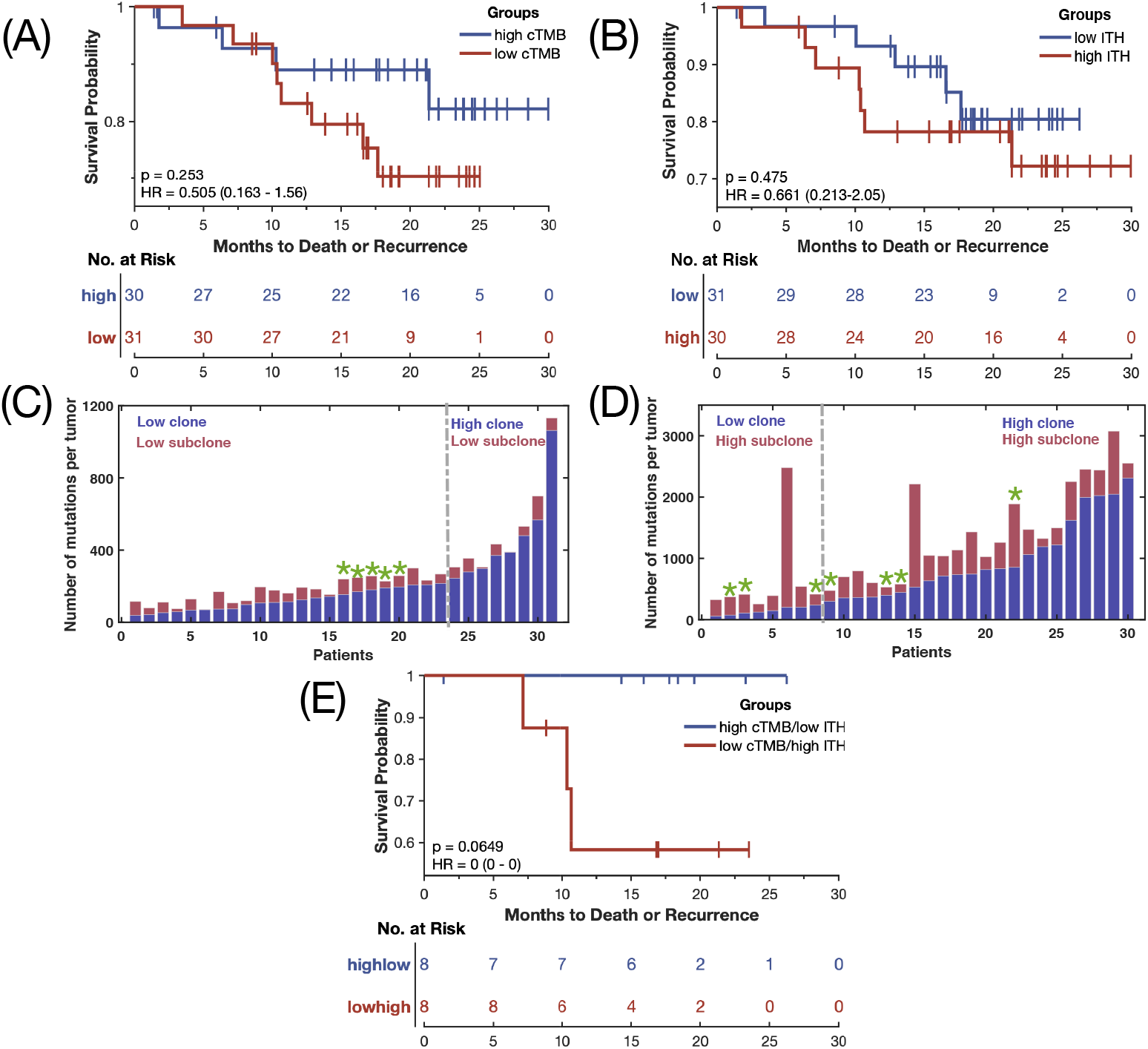
The influence of tumor mutation burden and intratumor heterogeneity on cancer (lung adenocarcinoma^60^) patient’s survival. (A) The relapse-free survival probability over a 30-month period for patients under high (above the median)/low clonal (below the median) tumor mutation burden (cTMB). (B) The relapse-free survival probability for patients under high (above the median)/low (below the median) intratumor heterogeneity (ITH, number of subclonal mutations). (C) The number of clonal (violet) and subclonal (red) mutations for patients with low ITH. (D) The number of clonal and subclonal mutations for patients with higher ITH. The dashed dotted lines in (C) and (D) separate the population into two categories based on cTMB. Green stars in (C) and (D) indicate recurrence or death. (E) The relapse-free survival probability over a 30-month period for patients under high cTMB and low ITH (low cTMB and high ITH) shown by the blue (red) line. The p-value and hazard ratio are shown in (A), (B), and (E) using the log-rank test.

Based on these observations, instead of a single variable, we combined the cTMB and ITH and separated the patients into four groups (see Figs. 8C-D). We found that only the patient group with high cTMB but low ITH does not have recurrence or death events (see Fig. 8C). A much higher survival probability (with a hazard ratio of zero) is found in this group relative to the one with low cTMB and high ITH in a marginal significance (see Figs. 8E). The reconstructed phylogenetic trees (see Figs. S27-28) show entirely different structures for the two categories of cancer patients. Therefore, instead of using one variable (TMB or ITH), a combination of these two factors could better predict the patient survival. These findings for lung adenocarcinoma show that the cancer biomarker landscape must at least be two dimensional, one representing ITH and the other TMB. Interestingly, a recent study^61^ has arrived at similar conclusions. By combining experiments on UVB-induced melanoma in mice and probing the immune response and analysis of TCGA melanoma patients, they discovered that the best survival probability is found for patients when TMB is high but ITH is low and the worst scenario is observed in patients with low TMB but high ITH (Fig. 1E in^61^). Remarkably, this is also reflected in Fig. 8E, although more experiments in a large clinical sample are needed to support the conclusion. It does appear that both TMB and ITH, in principle measurable, could potentially predict survival probability more accurately, although high ITH is almost always detrimental^11, 62, 63^.

## III. DISCUSSION

We investigated the evolutionary signatures focusing on the mutation burden in cancer patients, and their consequences on disease risks, variations between the sexes, and potential biomarkers for cancer diagnosis and prognosis. By analyzing the data for 16 cancers from the WES and 24 cancers from the WGS, we found that two distinct scenarios for the accumulation of somatic mutations during cancer progression is determined by a single parameter. If the overall mutation burden is low (*<* 3 mutations/Mb), a strong positive correlation exists between the number of mutations and the patient age at diagnosis, which is absent for high TMB cancers. If the TMB is suitably rescaled then the distribution is a standard Gaussian for all cancers with low mutation load. In contrast, a power-law distribution of TMB is found for high TMB cancers. The mutation profiles along each chromosome are different for the two groups with high TMB cancers exhibiting a large number of mutations within small segments of the chromosomes. The variations of the mutation frequency along cancer genome could be caused by chromatin structure, replication timing and other factors^64^. We also found a few exceptions with high TMB where the positive linear relation between these two quantities is recovered for cancers such as HNSC, STAD, and SKCM. Interestingly, all these types of cancer are strongly influenced by environment^41^, which we explained by including the effect of extrinsic factors on mutations within the framework of a simple model.

For cancers that show a linear relation between the TMB and PAD, half or more mutations appear before tumor initiation^10^, which might be due to the relatively short time from tumor initiation to diagnosis relative to the duration of the tissue self-renewal. In sharp contrast, the majority of the mutations appear during the tumor expansion stage for other types of cancer, which exhibit weak correlation between the TMB and PAD, such as colorectal cancer^12^. In other words, the majority of the somatic mutations appear in the trunk of the cancer phylogenetic tree for cancers with low TMB while most of them would arise in the branches for high TMB cancers, as illustrated in Fig. 3E. This is consistent with the observations in other studies on acute myeloid leukemia, chronic lymphocytic leukemia, melanoma, and colorectal cancers.^10–12, 34^. Our conclusion can be tested for other different cancer types with similar studies in the future.

The evolution of mutations in cancer cells, quantified using TMB, can serve as a valuable prognostic biomarker, allowing for a more informed treatment. Analysis of data from patients who underwent immunotherapy shows that better response in melanoma and non-small cell lung cancer is obtained in patients with high TMB^22^. In addition, recently FDA has approved pembrolizumab for several other cancer types with high TMB^59^. Nevertheless, TMB alone is not a perfect predictor for patient response^24^. We found that the fraction of older patients who show response to immunotherapy is much higher than younger ones under the same level of TMB. A further study leads us to conclude that two variables, TMB and ITH, could serve as a potential biomarker.

Apart from the genetic factors, the complex microenvironment of tumor cells is found to play a critical role during tumor initiation, progression and metastasis^65–67^. Some features of the tumor microenvironment (TME) is related to patient response to immune checkpoint blockade (ICB) therapy^68^. Therefore, a deep analysis of TME factors, such as the expression of PD-L1, the level and location of tumor-infiltrating CD8+ T cells and other features is likely to reveal additional biomarkers^68, 69^. Finally, we can search for an optimized combination of biomarkers from somatic mutations and TME factors by data mining from TCGA and other resources^26^ in order to improve the prediction of patient survival and response.

The cancer risk for different tissues has been reported to be strongly correlated with the number of the stem cell divisions^15^ while a recent research claims that the cancer risk is strongly correlated with the mutation load^46^. One of our key finding is that the mutation burden is indeed correlated with the cancer risk. Interestingly, mutation burden alone explains the disparity of cancer risk found between the sexes after we exclude sex-hormone-related cancers. However, the present and previous studies^46^ show that the mutation load alone cannot account for all the differences in cancer risk among different tissues. Therefore, factors such as the number of driver mutations, immune responses, and sex hormones must be important in understanding the variations in the cancer risks among different tissues^47–49^.

The signatures linking TMB and cancer risk is also imprinted in mutations at the base pair levels in the chromosomes. For instance, driver genes, such as DNMT3A and TET2 (see Fig. S14), are significantly mutated in LAML cancer patients. Meanwhile, recent findings show that such mutations are also found in apparently healthy persons, and are associated with enhanced cancer risk^70^. Thus, there is a link between chromosomal abnormalities and TMB. Because TMB, a coarse-grained measure alone may not be sufficient as a biomarker, it should be combined with mutations at the chromosome level to assess cancer risk.

A deeper understanding of the evolutionary dynamics is the key to devise effective therapies for cancer. Although, we have investigated this process for more than 20 cancers covering a large population of patients, it would be valuable to assess if our findings are also applicable to other cancer types. Additional data will be needed to verify the correlation between cancer risk and the mutation load for KICH and other cancers between the genders. We have assumed that the mutation load found in cancer patients reflects the mutations accumulated in ancestral cells^46^. Therefore, we need more mutation profiles from normal and precancerous lesions to further validate the conclusions reached here^12, 25, 33^. In addition, the influences of other types of mutations such as copy number variations (no clear correlation is observed between the CNV burden and PAD from our preliminary analysis), and aneuploidy should also be considered in the future. Due to the complexity of cancer diseases and presence of many different risk factors simultaneously, it is valuable to evaluate one factor at a time (even if additivity principle does not hold), if at all possible, to better understand their roles in cancer progression.

## MATERIALS AND METHODS

### Genome data and bioinformatics pipeline

The Cancer Genome Atlas (TCGA) research network has collected and analyzed the genomic, protein and epigenetic profiles of more than 11,000 specimens from more than 30 cancer types^26, 71^. We focus on the link between tumor mutation burden (TMB) and cancer risk as well as discernible differences between the sexes based on TMB. Hence, we choose 16 major cancers, and excluded data from cancers with small samples and those that are gender specific. We take the mutation data for each cancer from the Firehose pipeline developed by the Broad Institute (http://firebrowse.org). The MuTect algorithm^72–74^ is used by the platform to detect the mutation profile of cancer patients. The patient age and clinical information are obtained through the RTCGAToolbox^31^. The data with patient age and mutation information that are missing in the database are excluded from our analysis. The number of patients for each cancer type is listed in Table I in the Supplementary Information.

Some details of the bioinformatics pipeline for the whole-exome sequencing (WES) data: the standard parameters in the MuTect algorithm are used to detect the single nucleotide variants (SNVs) which can be found in the original reference^74^. Bayesian statistical analysis of bases and their qualities in the tumor and normal Binary Alignment Maps (BAMs) is used by MuTect to identify candidate SNVs at a given genomic locus. A minimum of 14 reads in the tumor and 8 in the normal tissue is required for mutation calling. The minimal allelic fraction cutoff is 0.1. A panel of normal samples as a filter to reduce false positives and miscalled germ-line events. The public dbSNP database is used to distinguish two cases: (i) sites which are known to be variant in the population and (ii) all other sites. More details about the method can be found in the original article^74^.

The whole-genome sequencing (WGS) data are taken from the ICGC/TCGA Whole Genomes Consortium which collects the genome data from 2834 patients across 38 tumor types^27^. Four different algorithms: MuTect, RADIA, Strelka, and SomaticSniper are used to detect the SNVs (at least called by 3 out of the 4 algorithms or called by SomaticSniper and one other algorithm). The minimal allelic fraction cutoff is 0.1. The SNVs are further filtered by the database dbSNP. More details about the SNV calls can be found in the SI of the reference^27^.

### Distribution of tumor mutation burden (TMB) for the sixteen types of cancer for different age groups

The distribution of TMB for distinct age groups (five years interval for each group) for the 16 cancers are shown in Figs. S1-S3 using the violin plot representation. The mean value of TMB for each group of patients is indicated by the red dot in the figure. A box plot is also shown within each violin plot. For better visualization of the TMB distribution over the whole age regime a few points with a relatively high mutation burden are not shown in some of the figures. Here, we took the latest data from the TCGA database with a much larger number of cases in contrast to previous studies^10^ which used an old version of the database with a limited number of cases. As an example, the total number of COAD and READ patients is 224 in one previous version of the database^10^, while we have 486 patients for the two cancer types here. We also separate the cancer types based on their origins, instead of combining them together as in previous studies^10, 27^. The different overall mutation rates (see Figs. 1-4) reflect the different underlying evolutionary processes of these cancer types. We used 3,000 Mb (the whole genome length) to calculate the mutation rates *µ*, the total number of mutations per year for the whole genome (not the whole-exome, with the length of about 30 Mb), in order to be comparable to other experiments directly.

Binning age groups does not change our results same as observed in previous studies^30^. In addition, we found similar results by using all patient data from WGS (see Fig. 4 and the Pearson/Spearman correlations shown Table VII in the Supplementary Information). Besides, we found that a rather accurate mutation rate could be reached by binning the WES patient data by ages as the value we obtained is almost the same (13/year for LAML as an example, see Fig. 1A) as measured in other studies (13±2 for LAML in the reference^34^). Therefore, our results are robust and do not depend on which method we used for patient data.

**TABLE VI:**
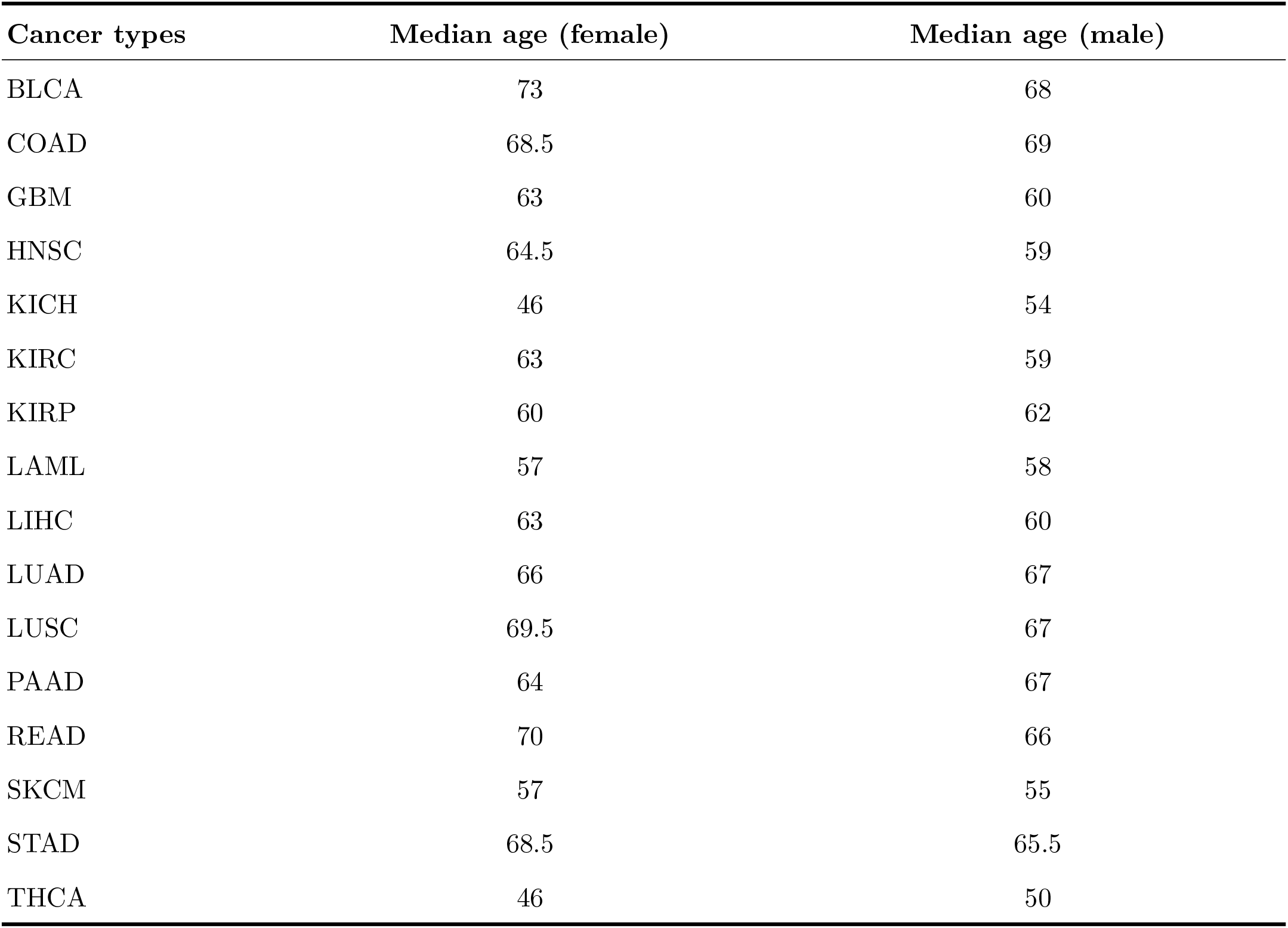
The median age (years) for female and male patients at diagnosis across the 16 types of cancer from TCGA database^31^.

**TABLE VII:**
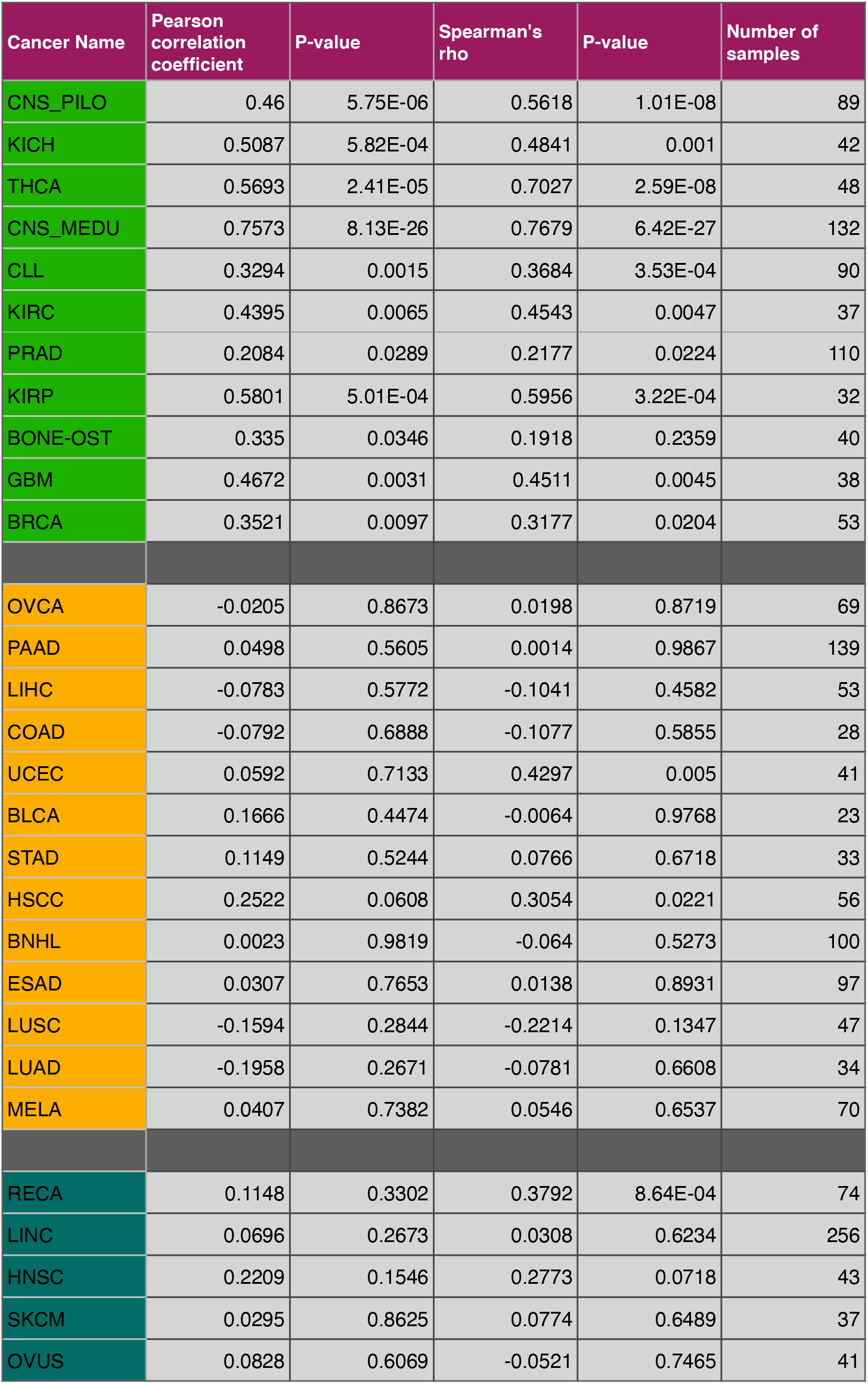
Correlation between TMB of SNVs and PAD for 24 cancer types from WGS data^27^. Two different methods (The Pearson and Spearman correlations) are used to calculate the correlations. The correlation coefficient, P-value and the number of patients for each cancer type are listed in the table. Five of the 24 cancer types from different countries are also calculated separately and listed at the end of the table. Cancer name abbreviations: CNS PILO, idney Chromophobe; THCA, thyroid Carcinoma; CNS MEDU, kidney renal clear cell carcinoma; PRAD, kidney renal papillary cell carcinoma; BONE-OST, bone osteosarcoma; GBM, glioblastoma multiforme; BRCA, breast invasive ductal carcinoma; OVCA, ovary adenocarcinoma (AU); PAAD, pancreatic adenocarcinoma; LIHC, liver hepatocellular carcinoma (US); COAD, colon adenocarcinoma; UCEC, uterus adenocarcinoma; BLCA, bladder urothelial carcinoma; STAD, stomach adenocarcinoma; HSCC, head-and-neck squamous cell carcinoma; BNHL, lymphoid mature B-cell lymphoma; ESAD, esophagus adenocarcinoma; LUSC, lung squamous cell carcinoma; LUAD, lung adenocarcinoma; MELA, skin melanoma (AU); RECA, kidney renal clear cell carcinoma (EU); LINC, liver hepatocellular carcinoma (JP); HNSC, head and neck squamous cell carcinoma (US); SKCM, cutaneous skin melanoma (US); OVAU, ovary adenocarcinoma (US).

### The five-year relative survival rate of cancers and lifetime risk of cancers

The five-year relative survival rates^43^ for various cancers are shown in Fig. S4. Survival rate is defined as the ratio of the percentage of patients who are alive for 5 years after the diagnosis of cancer to the percentage of people in the general population alive over the same time period with respect to age, sex and race^75^. Three out of the four of the most lethal cancers shown in Fig. S4 (pancreatic, liver and lung cancers, denoted by the yellow box) have relatively high TMB (see Fig. 2). We do not have sufficient mutation data for Mesothelioma, the other lethal cancer, which requires wide and deep sequencing of the mesothelioma patients^76, 77^.

### Correlation of SNVs from the whole genome and exome

Among the 2834 donors of the ICGC/TCGA Whole Genomes Consortium, we used 2583 in this study which are found to have the optimal quality^27^. From the number of SNVs from coding and noncoding DNA of each patient, we plot the number of SNVs for the whole-genome (sum of the SNVs from coding and noncoding DNA) as a function of SNVs number from the whole-exome (see Fig. S5). Each circle represents the data from one patient. A strong linear correlation is found between these two variables (with the Pearson correlation coefficient *ρ* = 0.98 and a P-value smaller than 0.001). The exome fraction is about 1% of the total genome in human^78^, and the total number of SNVs in exome is also about 1% of the value from the whole genome (see the magenta solid line in Fig. S5). Therefore, our definition of tumor mutation burden (TMB, the number of SNVs per Megabase) does not depend on which dataset (WES, or WGS) is used. This is further confirmed by analyzing the WES and WGS data which show that the conclusions are robust.

### There are two types of TMB distribution for the sixteen cancer types

Instead of considering the TMB distribution over different age groups, we also investigated the overall TMB distribution for every cancer discussed in this article. From the patient mutation data obtained from TCGA^26^, we calculated the TMB distributions for the sixteen cancers (see Fig. S6 and Fig. S7). Remarkably, for the nine cancers listed in Fig. 1 and Fig. 3, we found that their TMB distribution can be well described by a universal Gaussian function (see the green lines in Fig. S6 and the corresponding functional forms are presented in the upper right box of the figures). In contrast, a power-law (P(TMB)∼TMB*^b^*) relation is observed (except for PAAD and LUSC) for those cancers with high TMB listed in Fig. 2 (see the green lines and their functional forms in Fig. §7).

For all cancers, the distribution of patient age at diagnosis (PAD) can be approximated by a Gaussian function (see the discussion below for the PAD distribution for both sexes and Fig. S19 and Fig. S20). Given a Gaussian distribution for PAD (see Fig. S8a) and a linear relation between TMB and PAD (see Fig. S8b) as noted in Fig. 1 and Fig. 3, we would obtain a Gaussian distribution for TMB (Fig. S8c), thus rationalizing the findings in Fig. 5c and Fig. S6.

### TMB distribution for both the sexes

We investigated the TMB distribution for both the sexes after separating the patient data into two groups according to their gender. Interestingly, similar patterns (see Fig. S9 and Fig. S10) are found (see Fig. S6 and Fig. S7). After rescaling the TMB and the frequency, the TMB distributions for the both sexes across the nine cancers (listed in Fig. 1 and Fig. 3) again collapse onto a single curve represented by the standard normal distribution (see Fig. S11a). In contrast, for cancers (except PAAD and LUSC) shown in Fig. 2, the rescaled TMB distributions collapse onto a single straight line with slope value of −1 (see Fig. S11b), which is the same as noted in Fig. 5d.

The rescaling approach used in Fig. 5 and Fig. S11 is often used in other areas in order to reveal whether there is a universal underlying mechanism hidden in the data set from different measurements. Our analysis clearly shows that a clock-like mutation process indeed exists in cancers shown in Fig. 5a, although the rate can be different for each cancer type. The value of *µ*_0_ for Fig. 5b is the mean TMB value for each cancer type. Because the curves for different cancers are universal one can, in principle, use an analytic formula to relate different cancer types to each other.

### The joint probability distribution for TMB and PAD

We also calculated the joint probability distribution P(X,Y) for the rescaled TMB (TMB/T*_med_*≡X) and PAD (PAD/P*_med_* ≡Y) where T*_med_* and P*_med_* are the median value of TMB and PAD, respectively (see Fig. S12 and Fig. S13). Due to the limited data for each cancer type, we could not obtain smooth distributions for P(X,Y). Nevertheless, we could still distinguish between the cancers with relatively low and high TMB from P(X,Y). The probability distribution P(X,Y) for cancers listed in Fig. 1 and Fig. 3 is symmetric along both axes (see Fig. S12). However, the P(X,Y) for most cancers shown in Fig. 2 is irregular and biased, extending to very large values along the horizontal axis (see Fig. S13). This is consistent with the power law fat-tail distribution observed in Fig. S7.

### Fraction (*F_ini_*) of accumulated mutations before the tumor initiation

In the main text, we estimated *F_ini_* for cancers shown in Figs. 1,3 and Fig 2. For cancers displaying a strong positive correlation between TMB and patient age (shown in Figs. 1 and 3), we calculated the number (*N_af_*) of accumulated mutations after tumor initiation from the mutation rate (*µ*, product of the genome length, 3,000 Mb, and the slope of the green curve in the figures), and the latency period (*τ_L_*) of the cancer, with *N_af_* = *µ* × *τ_L_*. Then, we can obtain *F_ini_* from *N_af_* and the total number of accumulated mutations (*N_t_*) for patients at the age of diagnosis (median value), which leads to

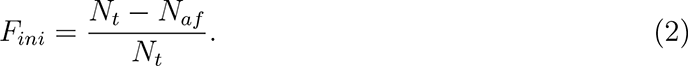

One example for THCA is discussed in the main text, with *µ* ≈ 30 mutations/year, *τ_L_* = 5 − 20 years and *N_t_* ≈ 2100 mutations, which leads to *F_ini_* ≈ 71% ∼ 93% as listed in Table II. For cancers with high TMB (shown in Fig. 2), we first calculated the upper bound of the total number (*N_I_*) of mutations a cell can accumulate before tumor initiation by taking the product of mutation rate (about 50 mutations/per year) for normal cells^12, 25, 33^, and the median age of patients at diagnosis of the cancer. The upper bound for *F_ini_* can be estimated from *N_I_* and the number of accumulated mutations (*N_t_*) for patients at the age of diagnosis (median value) as,

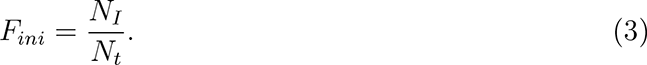

Take LIHC as an example, *N_I_* ≈ 50 × 61 ≈ 3050 (61, the median age of the LIHC patients) and *N_t_* ≈ 4 × 3000 ≈ 12000 (4, the TMB for LIHC patients at 61 years of age), which leads to *F_ini_* ≈ 25% as listed in Table III. For a few cancers (Fig. 2), the mutation rates for normal cells are not available. Hence, the value of *F_ini_* for these might show some derivations from our estimation. Using these two approaches, we estimated the values of *F_ini_*for the 16 cancers listed in Table II (for cancers shown in Figs. 1 and 3), and Table III (for cancers shown in Fig. 2). As discussed in the main text, the majority of mutations accumulate before the tumor initiation for cancers listed in Table II, while is precisely the opposite of what is found for cancers listed in Table III.

### Age distribution of cancer patients at diagnosis for both the sexes

There is an age disparity for patients at diagnosis among different types of cancer between the two sexes (see Table VI). In principle, we should not compare the cancer risks between the two populations with different ages at diagnosis directly because it is possible that one population could accumulate the same mutation burden at a younger age and at higher rate compared with the other population. The age distributions for both the sexes at diagnosis for 9 cancers (see Figs. S19) show that the median age (see the dashed line) for women is higher than for males. However, it is the opposite for 7 other cancers shown in Fig. S20. The solid lines in these two figures come from Gaussian fits. The mean value for the Gaussian distribution gives the same trend as the median age (indicated by the dashed lines) for the age disparity observed between the two sexes. The values of the median ages are used for analysis.

### The evolution of the number of mutations for both sexes

In our study, we assumed that both the sexes accumulate mutations in a similar fashion. Because the accumulation of mutations is a linear function of time in many types of cancers (see Figs. 1 and 3), we expect a similar trend should hold for the accumulation of somatic mutations in both female and male populations. We plot the TMB score as a function of the patient age for both the sexes in Figs. S21-S23. For cancers with low overall TMB (Fig. S21) or high TMB, but being strongly influenced by environment, (Fig. S23), a similar positive correlation between TMB and patient age is observed. Thus, both the sexes indeed accumulate somatic mutations in a similar way and we simply adjust their TMB with a factor of 66/P*_med_*, with 66 of the median age for all cancer patients and P*_med_* taken from Table VI. For cancers shown in Fig. S22, the TMBs for both the sexes almost do not change with the patient age. The correlation between cancer risk and TMB observed in Figs. S24a and S24b almost does not change after the age adjustment (see Figs. S24c and S24d), while the pattern shown in Fig. 7 is mainly changed only if the TMB score is low (THCA, KIRC, and GBM) compared with those shown in Fig. S25. The TMB for HNSC is relatively high but the same positive correlation is observed for this cancer due to the strong environmental influence. Therefore, the assumption that mutations accumulate in both the sexes in a roughly similar manner, as assumed in the main text, holds.

### Cancer risk and TMB without age-adjustment

We also examined the correlation between cancer risk and TMB by considering all the mutation data obtained from TCGA database (see Table IV). As cancer risk for the two sexes varies greatly, we included the data for both of them (see Table V) without considering age disparity (see Figs. S19 and S20) between male and female patients among different cancers. The Pearson correlation coefficient *ρ* = 0.65 is obtained for the cancer risk and TMB (see the best linear fit in Fig. S24c) similar to that (*p* = 0.6) found in Fig. S24a with age-adjusted TMB. A higher value (*p* = 0.7) is obtained after removing SKCM cancer as shown in Fig. S24d. This value again is almost the same (*p* = 0.69) as calculated using the data in Fig. S24b. Although cancer risk is related to the tumor mutation burden, the TMB alone cannot explain more than 50% (∼ 0.7^2^) of the differences in cancer risks among different tissue types.

### Cancer risk for both the sexes correlates with TMB after age-adjustment

The cancer risk as a function of TMB (without age-adjustment) for both sexes across different cancers are shown in Fig. S25. The data in this figure is the same as used in Fig. S24c but with lines connect the data for male and female populations for each cancer type. From Fig. S25a, it appears that there is no strong correlation between the cancer risk and TMB between males and females for cancers with low TMB. The red and blue lines show positive and negative correlations, respectively and the orange lines indicate the same TMB. On the other hand, the cancer risk is frequently associated with TMB as the latter reaches high values (see Fig. S25b). However, we neglected the known age disparity (see Figs. S19 and S20) between female and male patients at diagnosis to obtain the results. After we adjust the TMB by patient ages, as discussed in the main text, we find that the mutation burden is the critical factor in determining the different risks between female and male patients across many types of cancer.

### TMB, patient age and immunotherapy for metastatic melanoma patients

We investigated the influence of TMB, and patient age on response to immunotherapy. As an example, we first examined the age distribution for all the patients and the ones who show favorable response to immune checkpoint blockade from a clinical study for metastatic melanoma^79^ (see Fig. S26a). The age varies considerably in both the cases. The median age for patients who show response is 65 years while it is 61.5 years for the whole population. Meanwhile, the number of non-synonymous mutations for patients for the same cancer also varies greatly, as illustrated in Fig. S26b. The median value (309) of mutations for patients showing response is much higher than the value (197) of the whole population and a significant difference exists in the TMB between patients with and without clinical benefit to immunotherapy^79^. We observed that the TMB is correlated with the patient age for melanoma patients (see Fig. 3c). Thus, the patient age could also influence the responses to immunotherapy. However, we find that the patients whose age is ≥ 65 show a similar response (27%) as younger patients (22%).

In order to remove the influence of TMB and focus solely on age, we compared the patient age with different responses with similar TMB. For a similar mutation burden level, (*<*5% difference), we find a much higher fraction of old patients compared to young patients showing favorable response (see Fig. S26c, 63% versus 37%). A similar conclusion was reached by analyzing the data from other immunotherapy studies^23, 80^ with 75% and 100% fraction of older patients showing response but the sample sizes are smaller (12 and 6 cases, respectively).

We explain the data in Fig. S26c using the dynamics of accumulation of mutations discussed in the main text. A constant mutation rate is observed for melanoma patients (see Fig. 3c). A similar time period (*τ_d_*) for all the patients would be expected from the initiation to the detection of melanoma^10^. Consider two patients A and B with ages *T_A_* and *T_B_* with the same TMB, denoted by *N_T_*. The fraction 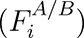 of mutations accumulated before the tumor initiation for both the patients is,

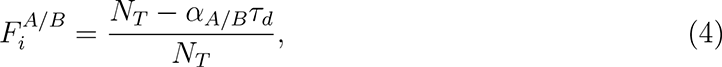

where *α_A/B_* is the rate for the mutation accumulation in patient A or B. If *T_A_ > T_B_*, then the mutation rate obeys *α_A_ < α_B_* because of *N_T_* for both the patients is the same. Therefore, the older patient would accumulate more mutations 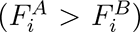 in the trunk of the tumor phylogenetic tree. A better response to immunotherapy would be expected if all tumor cells contain the neoantigens recognized by T-cells compared with the case that these neoantigens only appear in a tiny fraction of cell populations. This model explains the higher fraction of older patients showing response to immunotherapy with the same TMB, noted in the clinical studies cited above. This is consistent with the result of Figure 8(a) where a better survival probability is observed for the patient group with a higher clonal TMB.

### The phylogenetic trees for lung adenocarcinoma patients from multi-region whole exome sequencing

The SNVs obtained from the multi-region sequencing of each tumor are first clustered using the PyClone Dirichlet process clustering^81^. The mutation clusters are filtered based on the pigeonhole principle and crossing rule to ensure the constructed phylogenetic tree is accurate^60^. Based on the mutation clusters and also the values of the mean cancer cell prevalence, the phylogenetic trees (see Figs. S27-S28) are constructed using the tool CITUP^82^ for the lung adenocarcinoma patients discussed in Fig. 8E. A detailed description of the phylogenetic tree construction can be found in the SI of the reference^60^.

### Cancer risks for both sexes

#### Bladder Urothelial Carcinoma

The lifetime risk of urinary bladder cancer is 1.12% and 3.76% for women and men, respectively (https://seer.cancer.gov/csr/1975_2014/)^83^, and 95% of the cancers are bladder urothelial Carcinomas^84^. Therefore, the lifetime risk of bladder urothelial carcinoma for female and male is 1.12%×0.95=1.06% and 3.76% × 0.95=3.57%.

#### Colon Adenocarcinoma

The lifetime risk of colorectal cancer is 4.15% and 4.49% for women and men, respectively^83^. About 71% of colorectal cancers arise in the colon^85^. Thus, the lifetime risk of colon adenocarcinoma for female and male is 4.15%×0.71=2.95% and 4.49%×0.71=3.19%.

#### Glioblastoma multiforme

The lifetime risk of brain cancer is 0.54% and 0.7% for women and men, respectively^83^. About 81% of brain cancers are gliomas and glioblastomas represent 45% of gliomas^86^. Thus, the lifetime risk of glioblastoma multiforme for female and male is 0.54%×0.81×0.45= 0.20% and 0.7% × 0.81 × 0.45=0.255%.

#### Head and Neck Squamous Cell Carcinoma

The lifetime risk of oral or pharynx cancer is 0.68% for women and is 0.12% for laryngeal cancer^83^. For men, the lifetime risk of oral or pharynx cancer is 1.61% and it is 0.55% for Laryngeal cancer. Therefore, the lifetime risk of head and neck squamous cell carcinoma (including oral, pharynx and laryngeal cancers) for women and men is 0.80% and 2.16%, respectively.

#### Kidney Chromophobe

The lifetime risk of kidney and renal pelvis is 1.2% and 2.09% for women and men, respectively^83^. Among these cancers, 80% of them are from clear cell, 15% are from papillary cell and 5% are from chromophobe^87^. From these, we estimate the lifetime risk for kidney chromophobe carcinoma is 1.2% × 0.05 = 0.06% and 2.09% × 0.05 = 0.105% for women and men, respectively.

#### Kidney Renal Clear Cell

Similarly, the lifetime risk of kidney renal clear cell carcinoma is 1.2% × 0.8 = 0.96% and 2.09% × 0.8 = 1.67% for women and men, respectively.

#### Kidney Renal Papillary Cell Carcinoma

Again, the lifetime risk of kidney renal papillary cell carcinoma is 1.2% × 0.15 = 0.18% and 2.09% × 0.15 = 0.314% for women and men, respectively.

#### Acute Myeloid Leukemia

The lifetime risk of acute myeloid leukemia is 0.43% and 0.55% for women and men, respectively^83^.

#### Liver Hepatocellular Carcinoma

The lifetime risk of liver and intrahepatic bile duct cancer is 0.6% and 1.39% for women and men, respectively^83^. Hepatocellular carcinoma represents 90% of these cancers^88^. Hepatitis C infection is among 10% of all hepatocellular carcinoma^89^ and 1% of the US population is infected with Hepatitis^90^. According to the calculation in previous research^15^, the lifetime risk of liver hepatocellular carcinoma is 0.50% and 1.15% for women and men not infected by Hepatitis C, respectively.

#### Lung Adenocarcinoma

The lifetime risk of lung and bronchus cancer is 5.95% and 6.85% for women and men, respectively^83^. About 40% of these cancers are adenocarcinoma and 30% of them are squamous cell carcinoma^91^. Therefore, the lifetime risk is 5.95% × 0.4 = 2.38% and 6.85% × 0.4 = 2.74% for females and males, respectively.

#### Lung Squamous Cell Carcinoma

Similarly to the analysis for lung adenocarcinoma, the lifetime risk is 5.95%×0.3 = 1.79% and 6.85% × 0.3 = 2.06% for females and males, accrodingly.

#### Pancreatic Adenocarcinoma

The lifetime risk of pancreatic cancer is 1.54% and 1.58% for females and males, respectively^83^. About 96% of these cancers are exocrine cancers with adenocarcinoma being the dominant one of 95%^92^. The lifetime risk is 1.54% × 0.96 × 0.95 = 1.40% and 1.58% × 0.96 × 0.95 = 1.44% for females and males, respectively.

#### Rectum Adenocarcinoma

The lifetime risk of colorectal cancer is 4.15% and 4.49% for females and males as mentioned above^83^. About 29% of these cancers arise in the rectum^85^. Therefore, the lifetime risk of rectum adenocarcinoma is 4.15%×0.29=1.20% and 4.49%×0.29=1.30% for female and male, respectively.

#### Cutaneous Skin Melanoma

The lifetime risk of cutaneous skin melanoma for females and males is 1.72% and 2.77%, respectively^83^.

#### Stomach Adenocarcinoma

The lifetime risk of stomach cancer is 0.65% and 1.05% for females and males, respectively^83^. Adenocarcinoma represents about 95% of these cancers^92^. Thus, the lifetime risk is 0.65% × 0.95 = 0.618% and 1.05% × 0.95 = 0.998% for both sexes, accrodingly.

#### Thyroid Carcinoma

The lifetime risk of thyroid carcinoma for females and males is 1.79% and 0.63%, respectively^83^.

## Acknowledgments

We are grateful to Dr. S. Gail Eckhardt for her interest and useful comments, and Philipp M. Altrock, Shaon Chakrabarti, and Anatoly B. Kolomeisky for pertinent discussions. This work is supported by the National Science Foundation (PHY 17-08128), and the Collie-Welch Chair through the Welch Foundation (F-0019).

## Author contributions

X.L. and D.T. conceived and designed the project, X.L. performed the research, X.L. S.S and D.T. co-wrote the paper.

## TABLES

Table I shows the number of patients for each type of cancer used in our analyses for WES data. Table II and III give the fraction (*F_ini_*) of accumulated mutations before the initiation of tumors for two categories of cancer shown in Figs. 1, 3 and Fig. 2, respectively. Table IV gives the median value of (both synonymous and non-synonymous) mutations per megabase for both sexes across 16 types of cancer. Table V shows the cancer risk for both female and male populations in 16 different types of cancer. Table VI lists the median age for male and female patients at diagnosis. Table VII shows the correlation between TMB of SNVs and PAD for 24 cancer types from WGS data.

## SUPPLEMENTARY FIGURES

**Figure S1.**
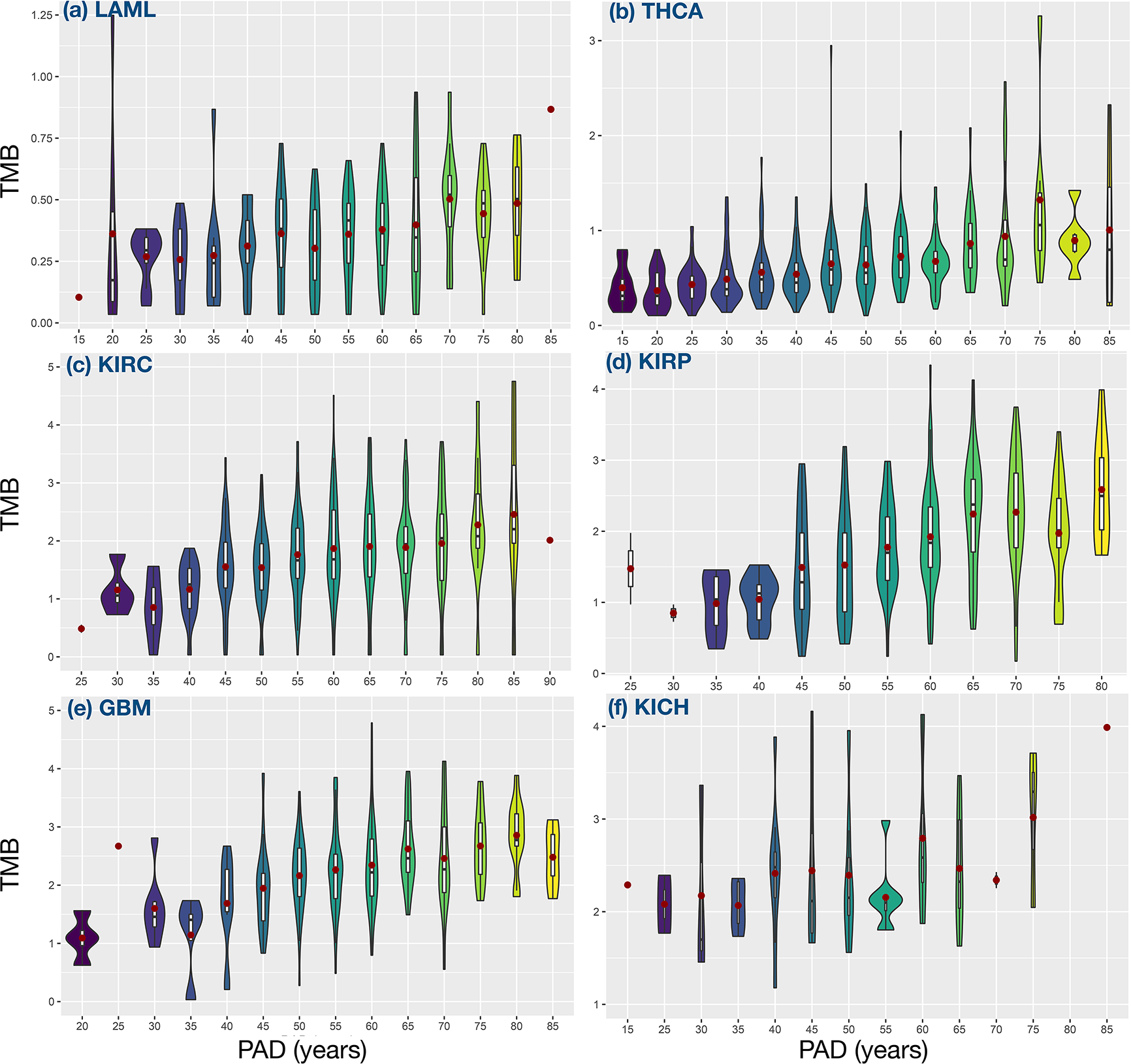
Violin plots for the distribution of tumor mutation burden (TMB) as a function of patient age for low TMB cancers shown in Fig. 1. The red dot is the mean value of the TMB. The inset with each violin plot shows a box plot. The unit for TMB is the number of mutations per Mb. The type of cancer is labeled in dark blue color. In Figs. (S2-S3), we show the same plot but for different cancer types. A few patient data with very high TMB are removed in Figs. (S2-S3) for better illustration.

**Figure S2.**
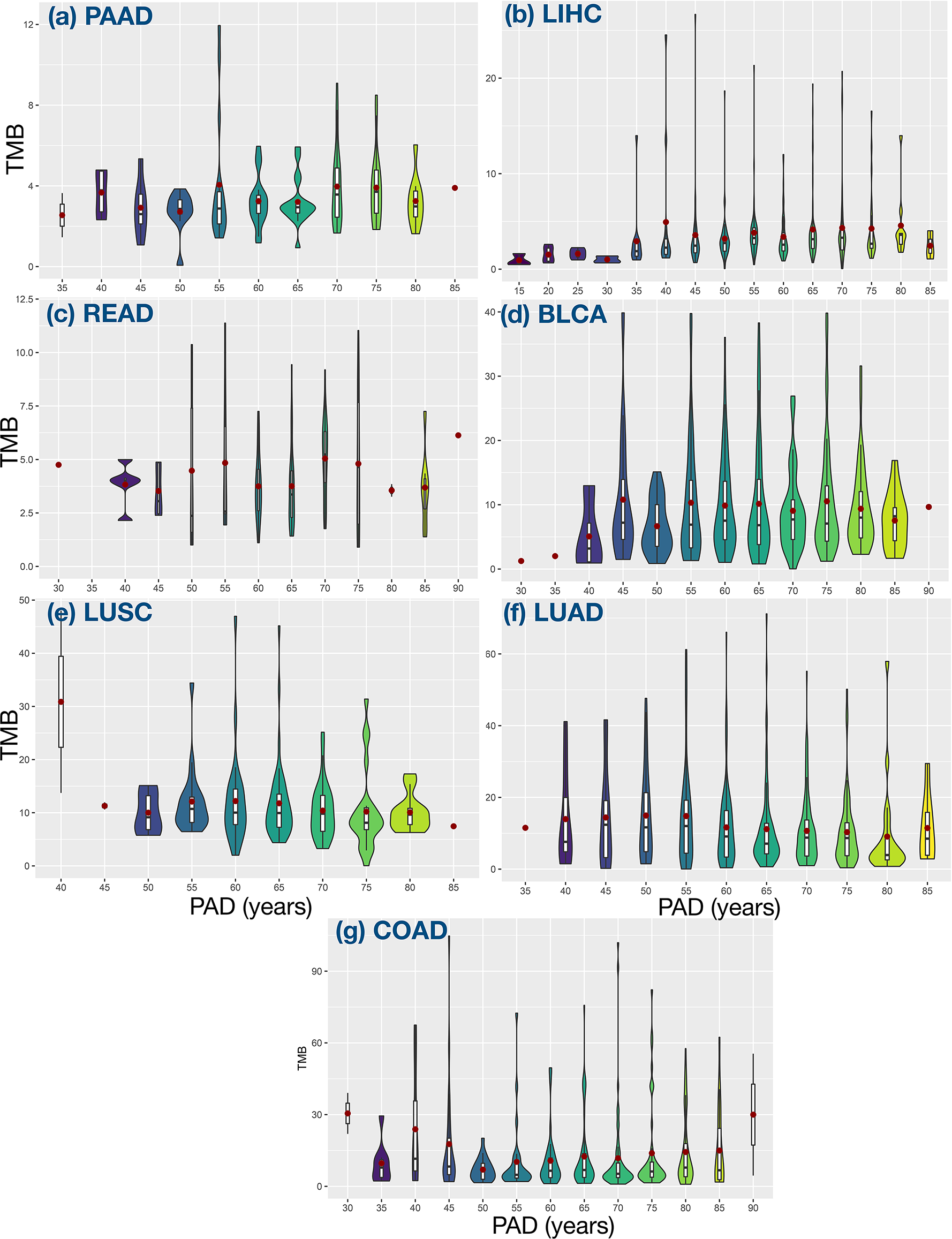
Violin plots for high TMB cancers shown in Fig. 2.

**Figure S3.**
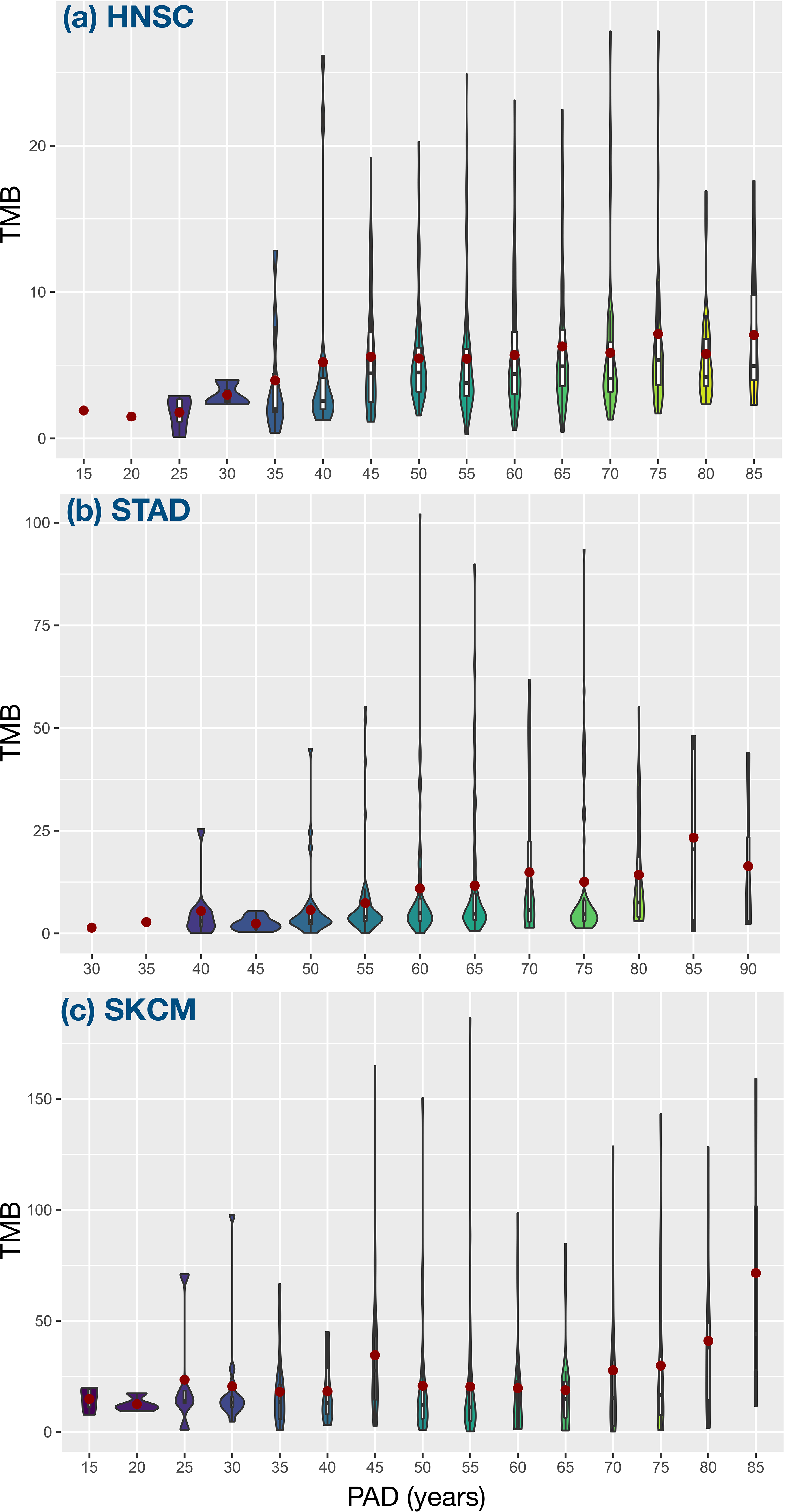
Violin plots for HNSC, STAD, and SKCM shown in Fig. 3.

**Figure S4.**
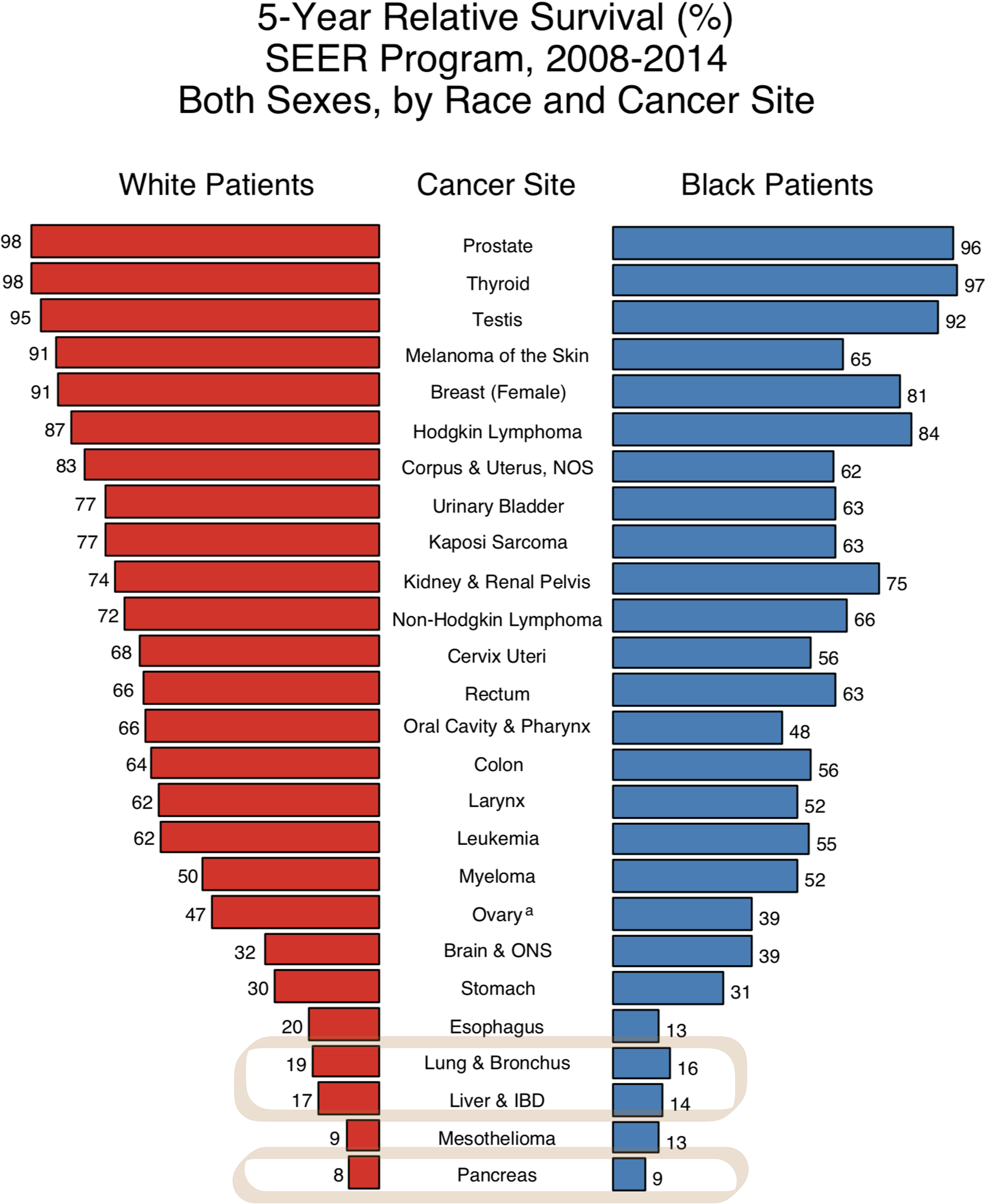
Five-year relative survival rate for different cancers^43^. Three of the most lethal cancers (lung, liver and pancreatic) in the box belong to the category shown in Fig. 2 in the main text.

**Figure S5.**
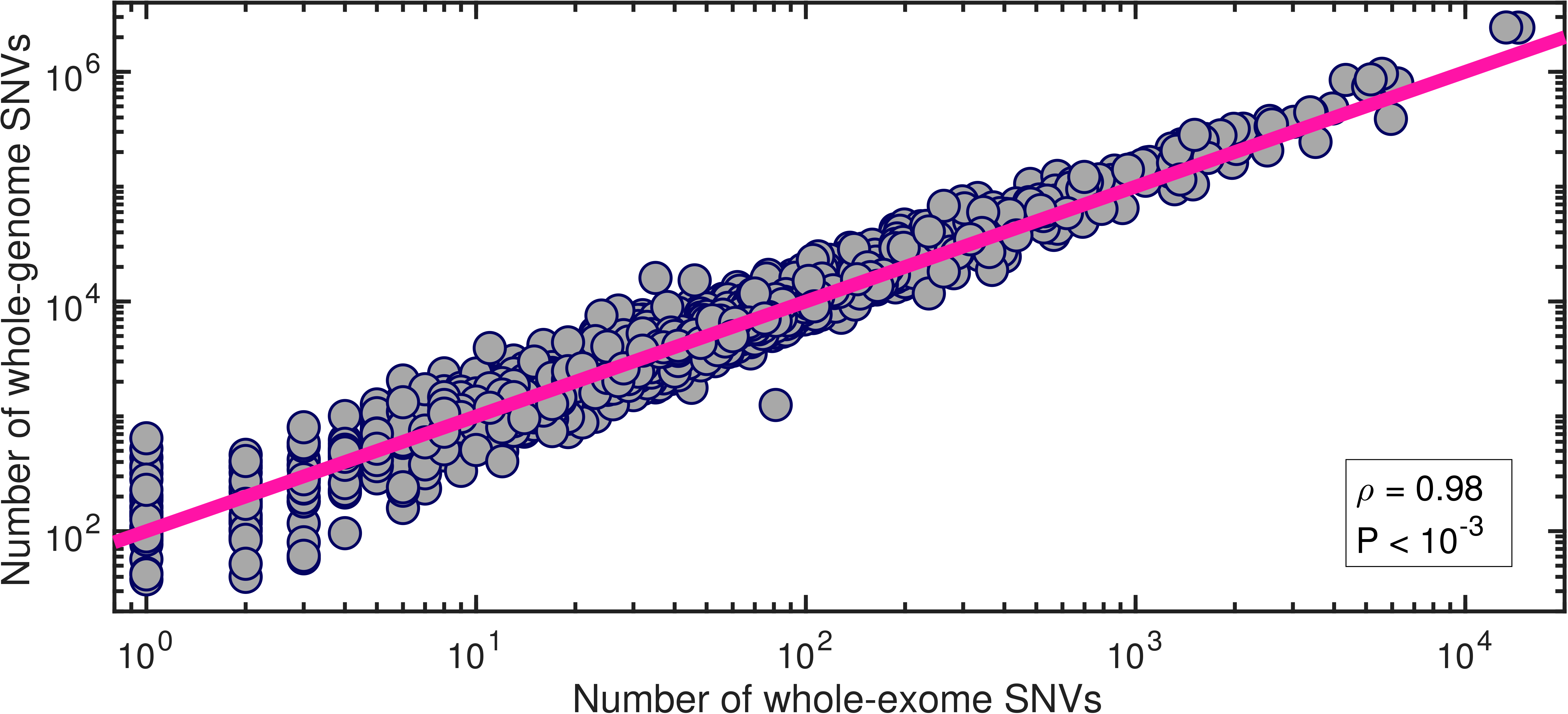
The correlation between the number of SNVs for the whole-genome and that for the whole-exome. The data (filled circles) are taken from the whole-genome sequencing of 2583 cancer patients across 38 tumor types^27^. A clear linear relation (with the Pearson correlation coefficient *ρ* = 0.98 and a P-value smaller than 0.001) is observed between these two variables. The line is described by the function: Y = 100X.

**Figure S6.**
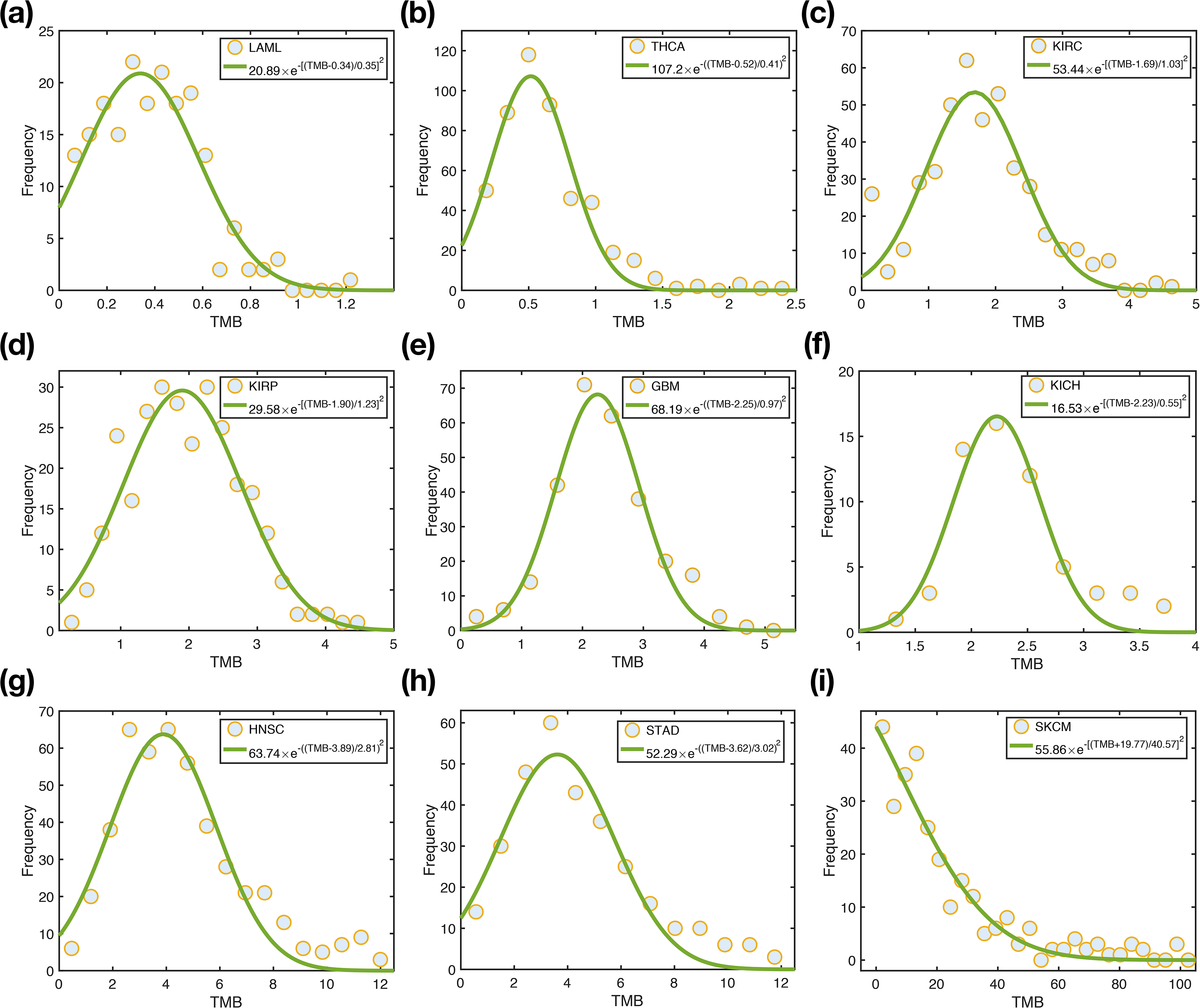
The TMB distribution over the nine types of cancer showed in Fig. 1 and Fig. 3 in the main text. The data (filled circles) are taken from the TCGA database as in the main text. The solid green curves (described by the functions in the upper right box of the figure) show the best fit to the data.

**Figure S7.**
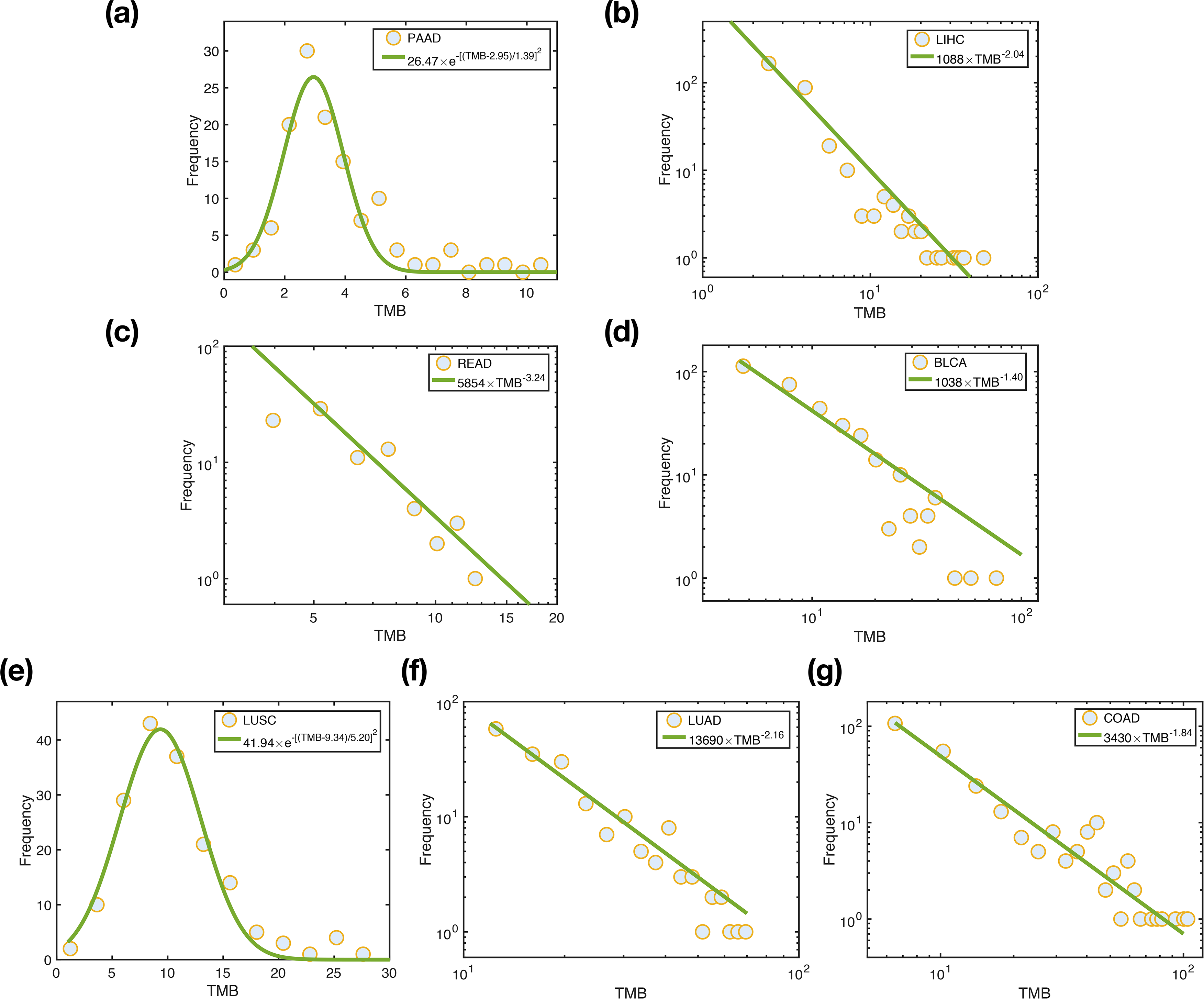
The TMB distribution for the cancers analyzed in Fig. 2. The meaning of the symbols and curves are the same as in Fig. S6.

**Figure S8.**
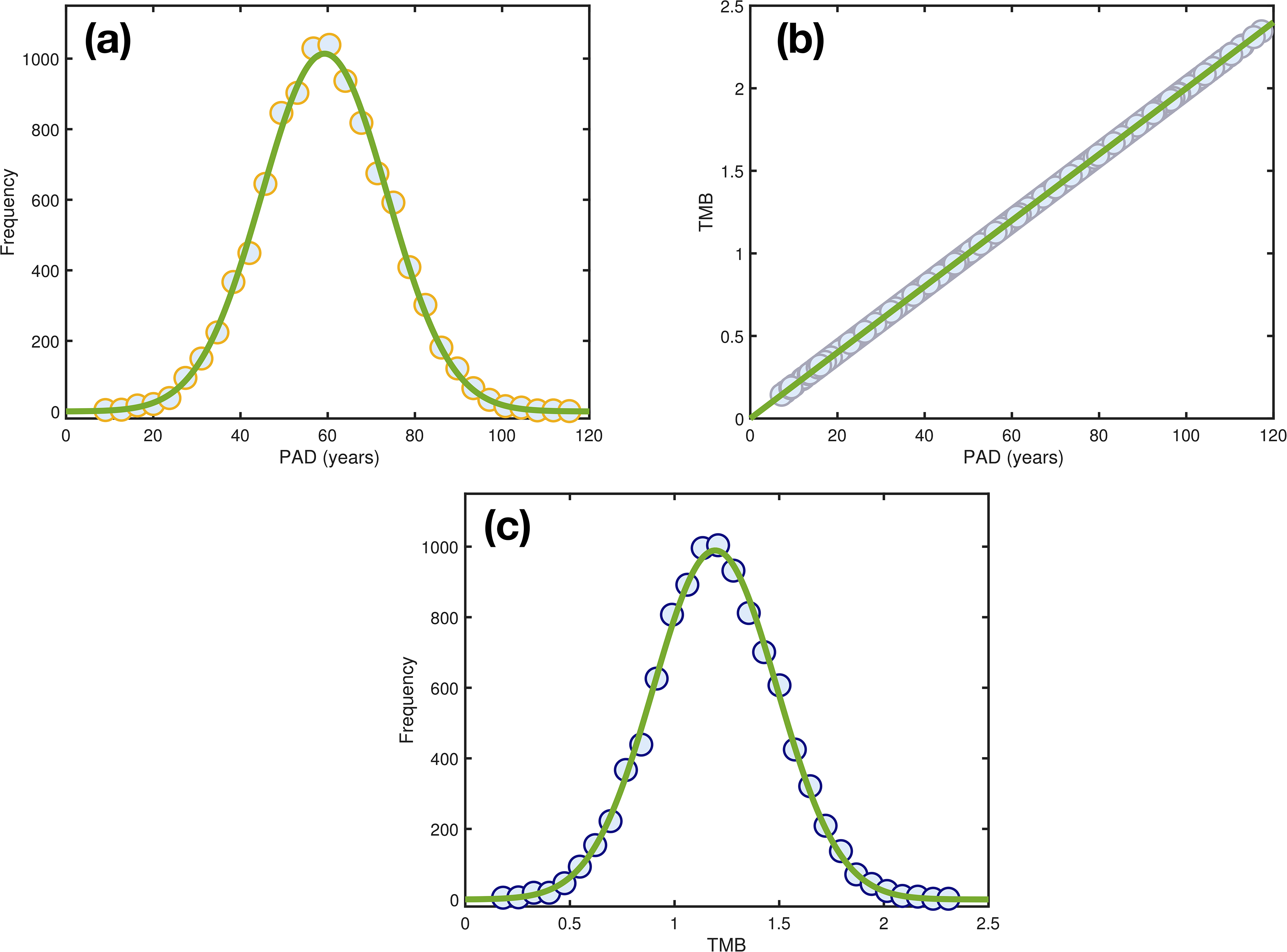
(a) The distribution of PAD obtained from a 10,000 sample using the Gaussian distribution. The green curve is described by a Gaussian function, 014×*e*^−[(PAD−59.28)*/*20.36]2^. (b) The TMB for each patient is proportional to their age (TMB = 0.02× PAD, see the green line). (c) The TMB distribution obtained from the 10,000 patient samples above. The green curve is again described by a Gaussian function, 989.3 × *e*^−[(TMB−1.2)*/*0.42]2^. The circles are simulation data and the green lines show the best fit to the data.

**Figure S9.**
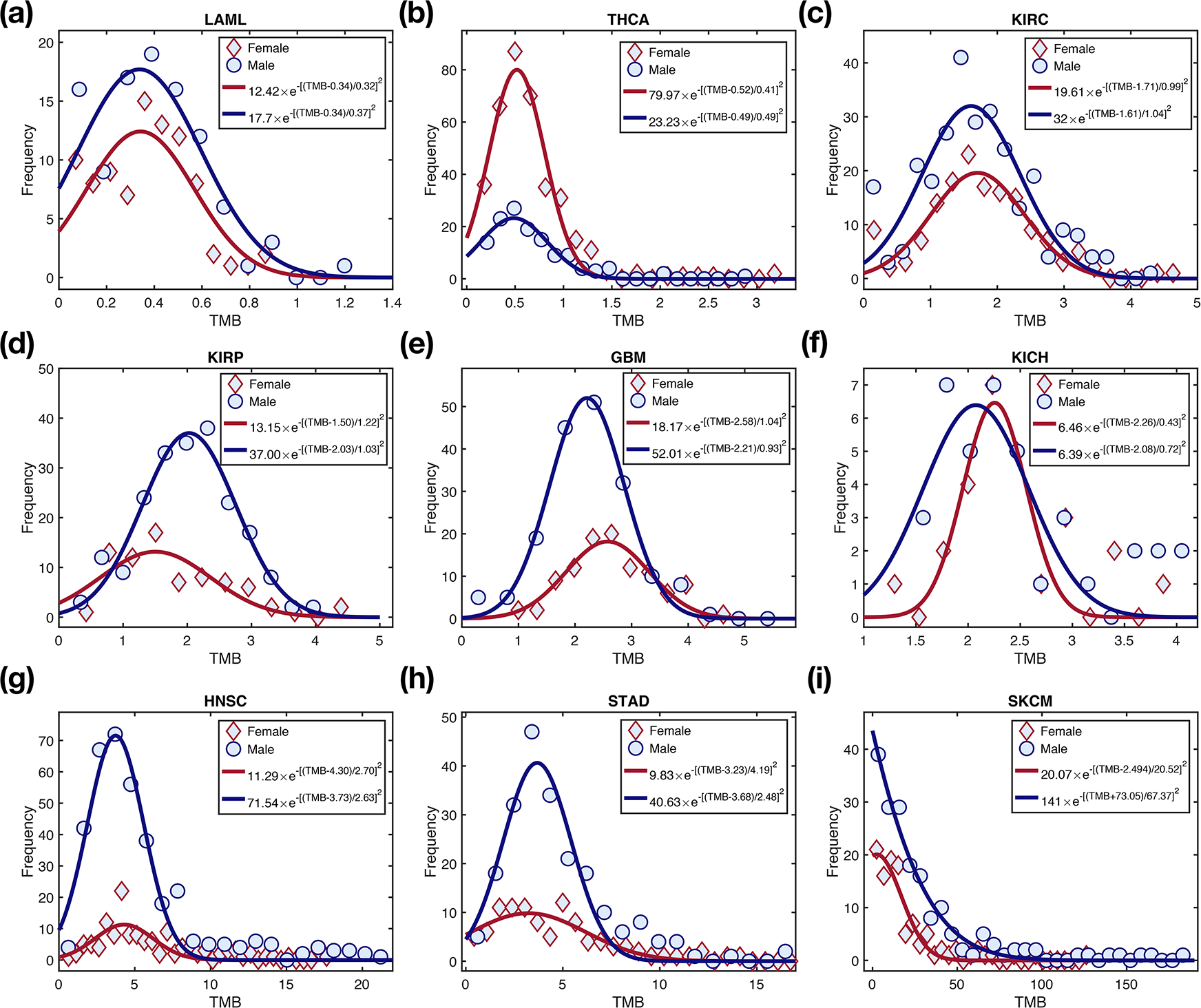
The TMB distribution for both the sexes for the nine cancers analyzed in Fig. 1 and Fig. 3. The data for females and males are represented by diamonds and circles, respectively. And the corresponding fitting functions (listed in the upper right box of the figure) for the data are shown by the red and blue curves.

**Figure S10.**
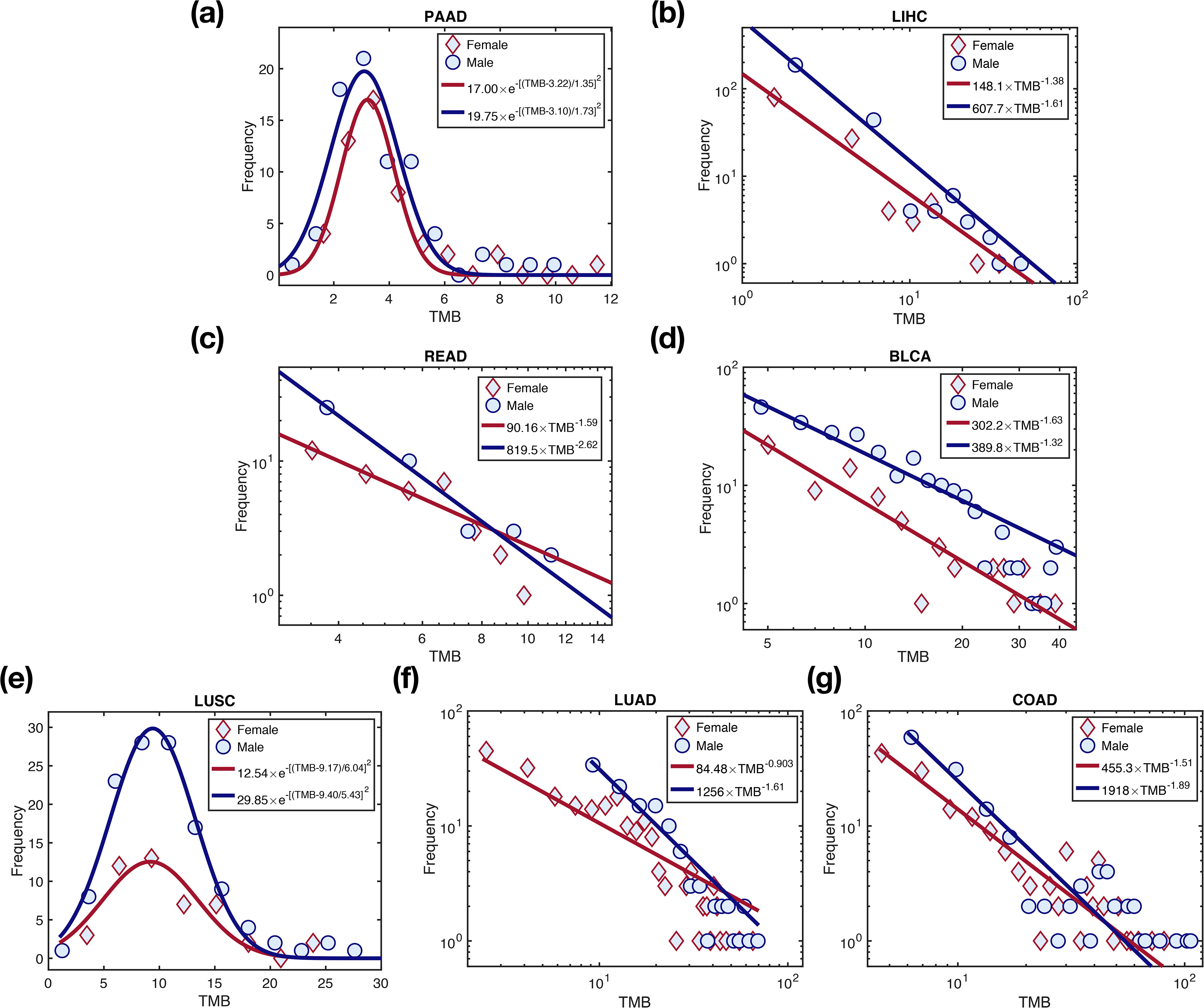
The TMB distribution for both the sexes over the seven cancers analyzed in Fig. 2. The meaning of the symbols and curves are the same as in Fig. S9.

**Figure S11.**
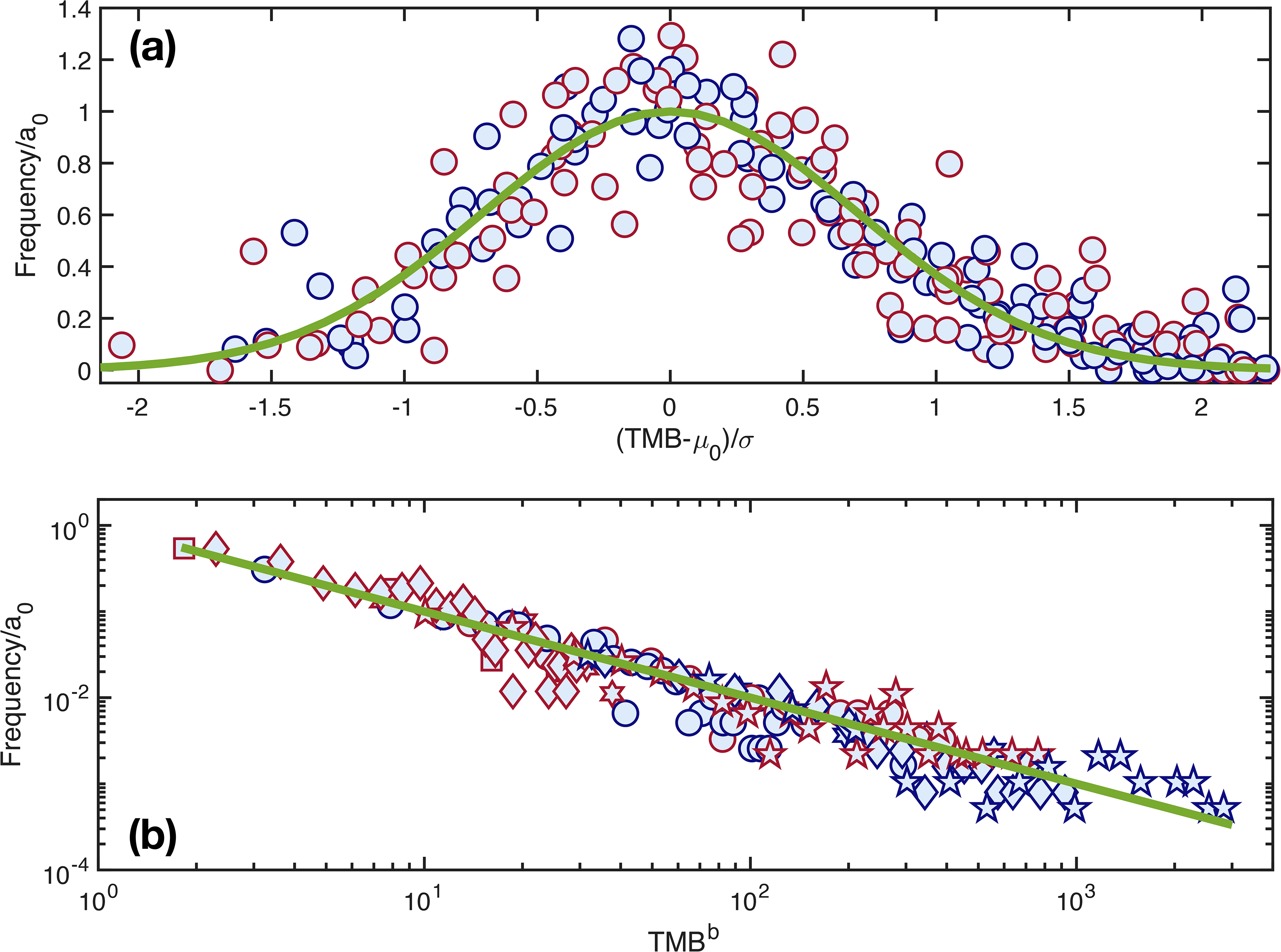
The rescaled TMB distribution for the 16 cancers. (a) A Gaussian distribution (with the mean value *µ*_0_, variance *σ*, and coefficient *a*_0_) describes (see Fig. S9) both the sexes for the low TMB cancers. The rescaled TMB distributions for these cancers collapse into the standard normal distribution (see the green line). (b) A long tail distribution, described by a power-law (see Fig. S10 showing *P* (TMB) = *a*_0_ × TMB*^b^*) is found for the cancers (except for PAAD and LUSC) listed in Fig. 2 in the main text. The green straight line has a slope value of −1. The red (blue) symbols represent data for females (males).

**Figure S12.**
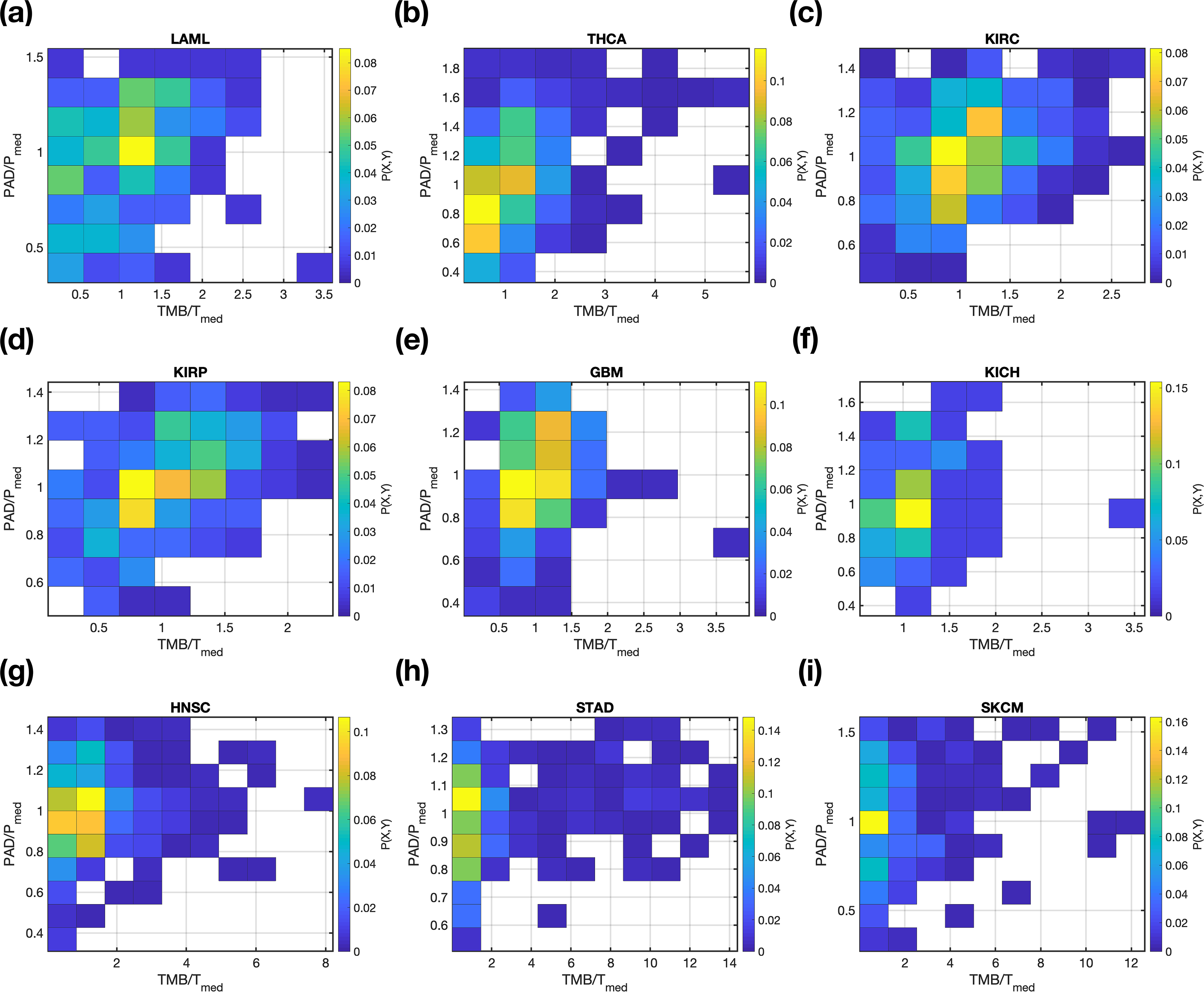
The joint probability distribution, *P* (*X, Y*), for TMB and PAD among the nine cancers. *X* = TMB/T*_med_* with T*_med_* the median value of TMB. *Y* = PAD/P*_med_* with P*_med_* the median value of PAD. The probability *P* (*X, Y*) is normalized by the total number of patients in each cancer and is color coded according to the value indicated by the scale on the right side of each figure.

**Figure S13.**
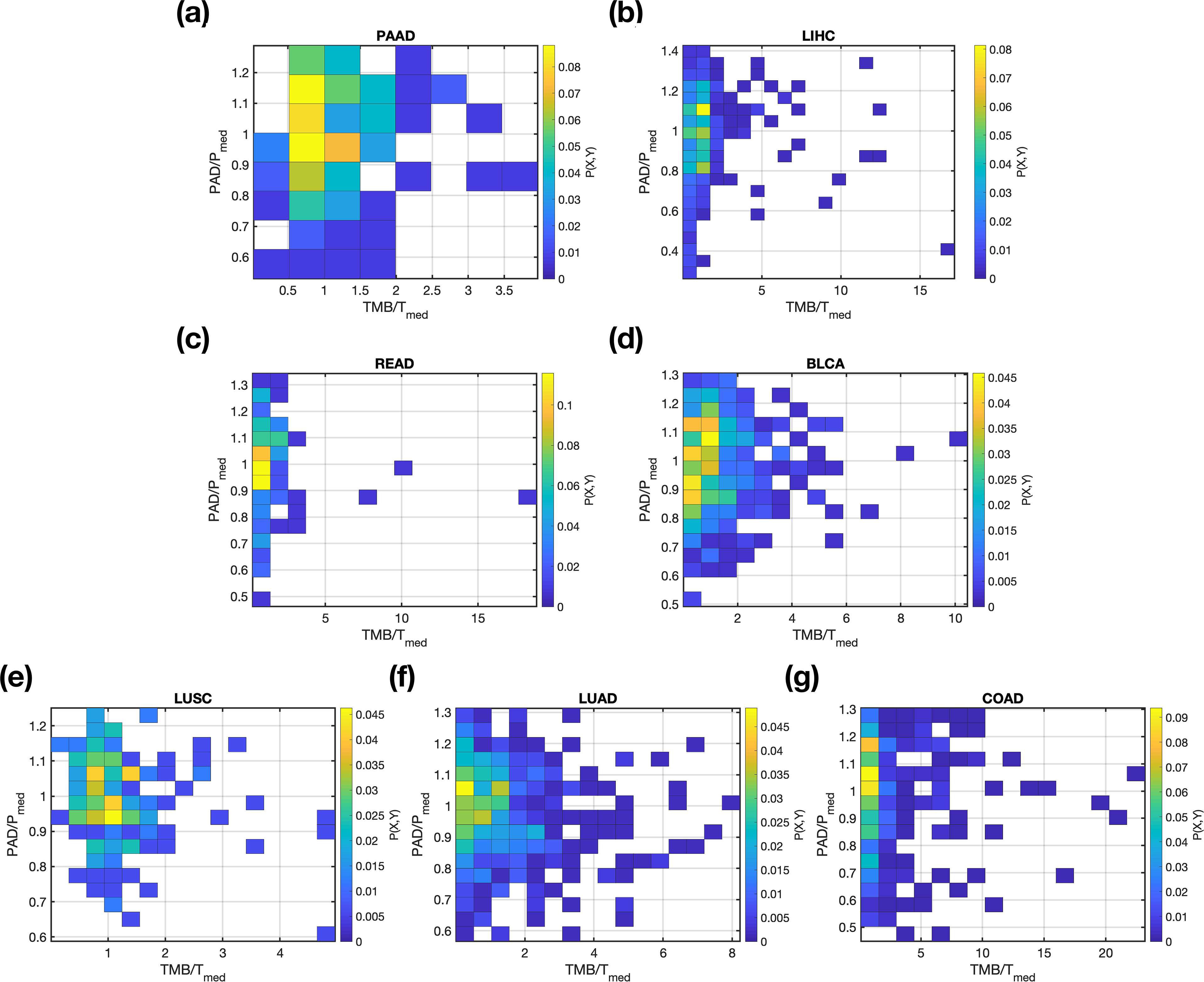
Same as Fig. S12 except these are for cancers analyzed in Fig. 2.

**Figure S14.**
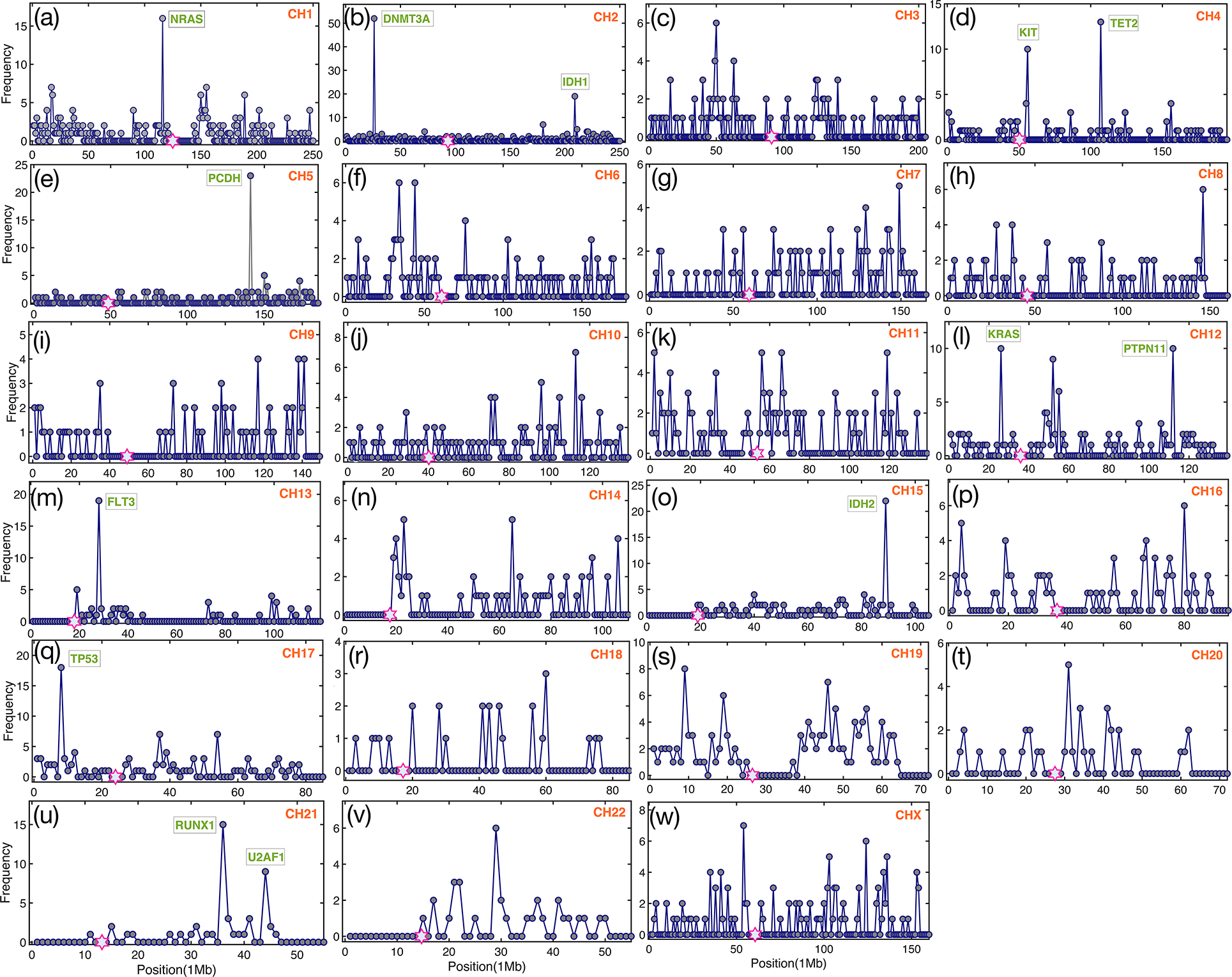
Tumor mutation frequency (number of mutations in a chromosome region of 1 Mb length) along each chromosome for the LAML. The genes (labelled in green), mutated at a high frequency, are listed in the figure. The hexagram in magenta shows the centromere location on each chromosome. The total number of patients is 197, taken from the Firehose pipeline (http://firebrowse.org).

**Figure S15.**
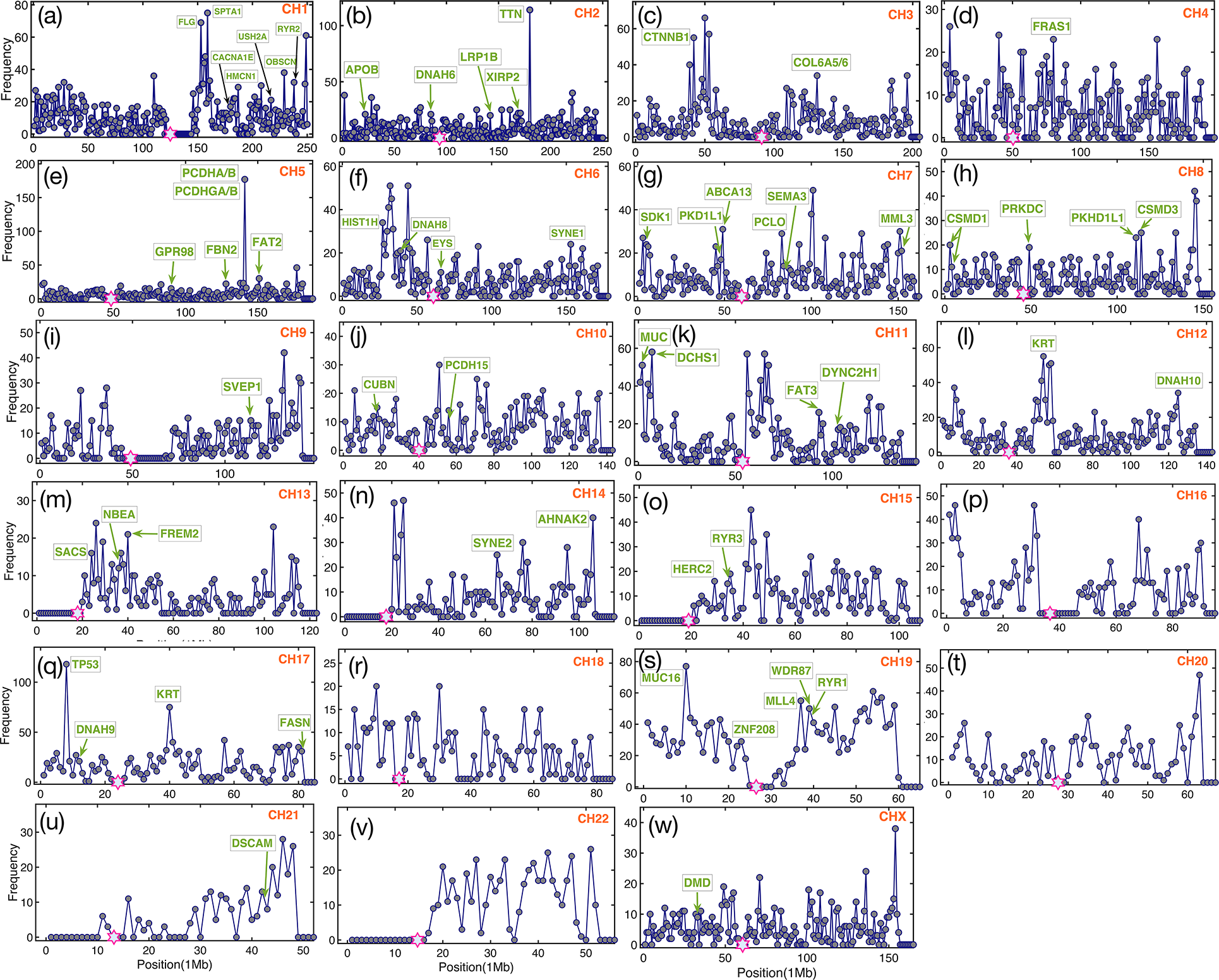
Same as Fig. S14 except these are for the LIHC cancer. The total number of patients used here is 198, taken from the Firehose pipeline (http://firebrowse.org).

**Figure S16.**
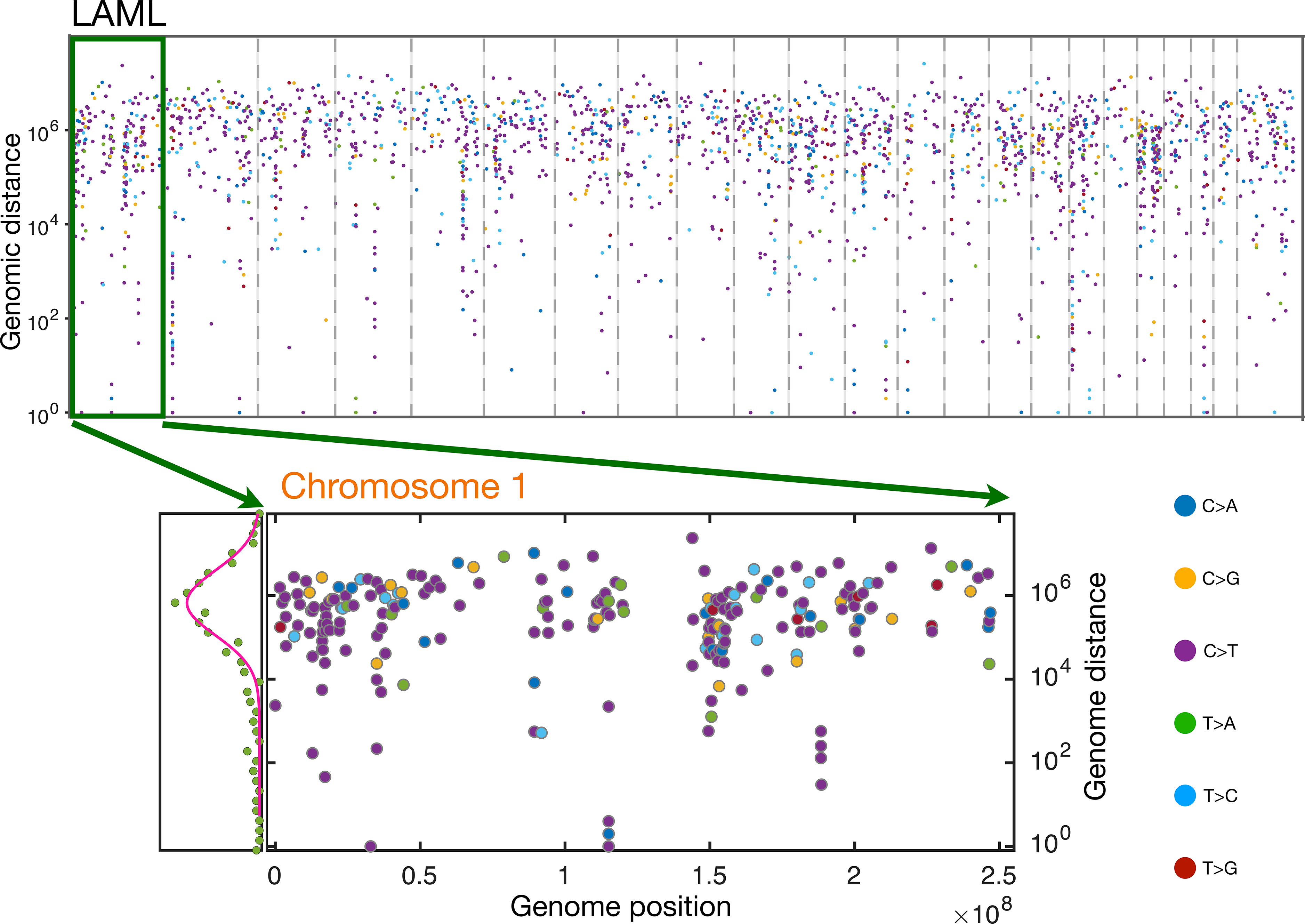
The upper panel shows the rainfall plot for the low TMB LAML (see also Fig. 5e in the main text). A zoom-in of the green rectangle region (chromosome 1) is shown in the lower panel. The distribution of genomic distance between two successive mutations in chromosome 1 is also shown on the left of the lower panel. The solid line, a Gaussian-function, provides an excellent fit to the data. The unit for genome position/distance is in basepair. The colors of the dots represent six different classes of base substitution, as demonstrated at the right of the figure. No hypermutation region is observed in this figure. The complete absence of SNVs in this chromosome close to the centromere region (around 125Mb) is striking.

**Figure S17.**
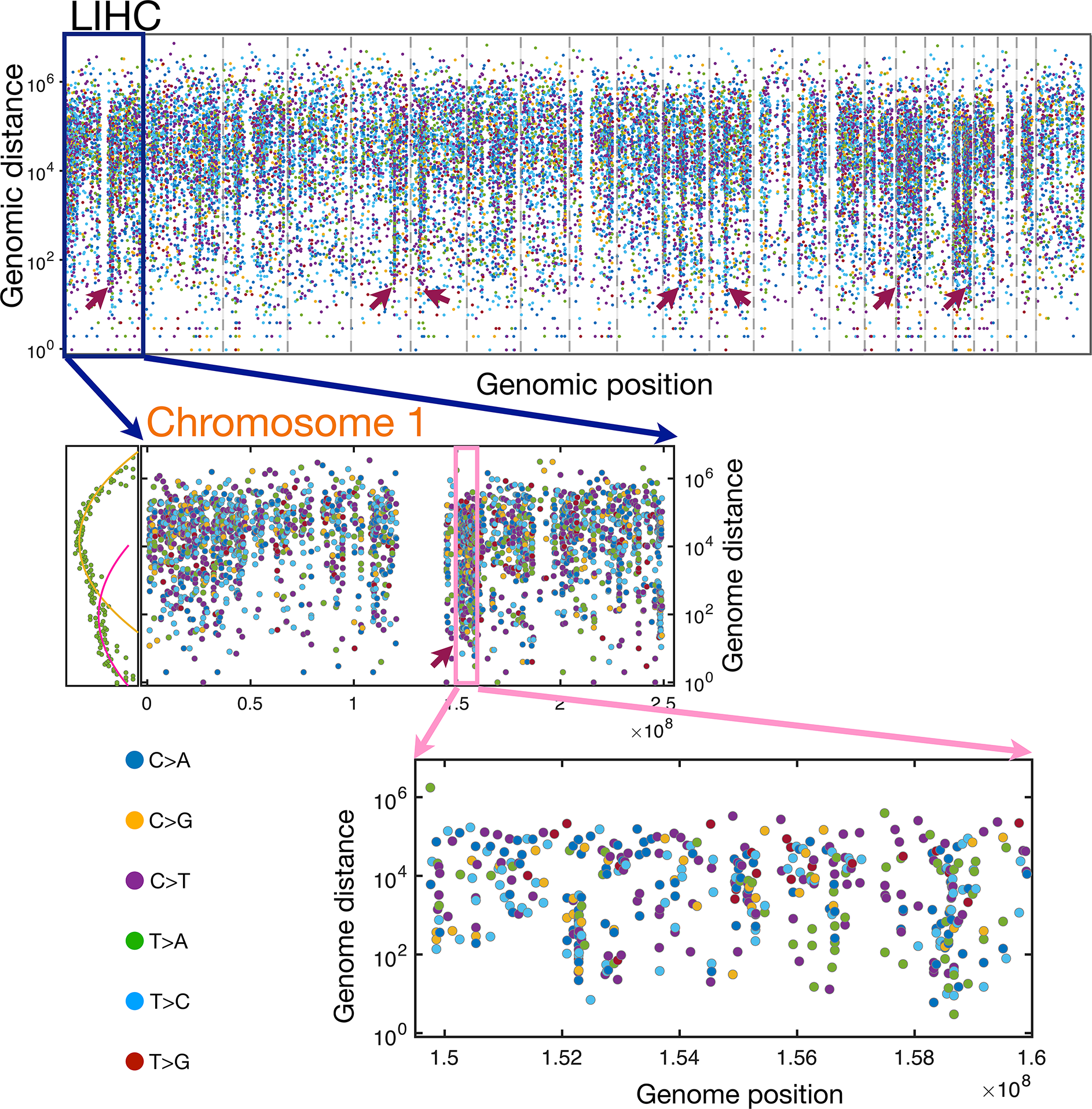
The upper panel shows the rainfall plot for LIHC (see also Fig. 6F). Just as in Fig. S16, we illustrate the rainfall plot for chromosome 1 only in the middle panel. The distribution of genomic distance is also shown on the left which can only be described by two Gaussian-functions (see the solid lines) instead of a single one used in Fig. S16. A zoom-in of the hypermutation region (see the pink rectangle) in chromosome 1 is demonstrated in the bottom panel. The red arrowheads indicate hypermutation regions. Comparison of the data in this figure and Fig. S16 illustrates stark differences in the density of mutations.

**Figure S18.**
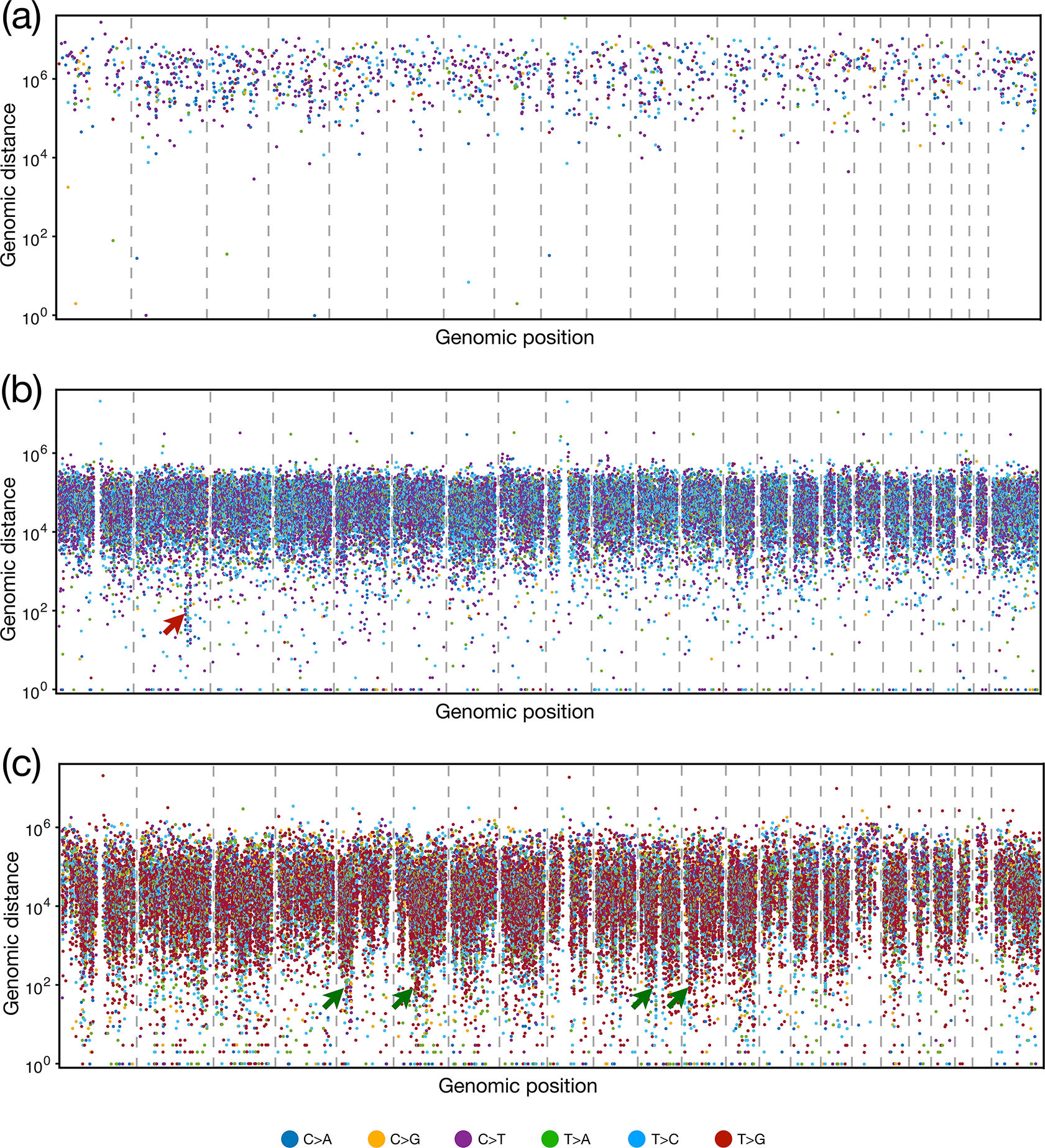
Kataegis patterns for the whole genome of (a) LAML, (b) LIHC, (c) ESAD cancer patients. The somatic mutations represented by small dots are ordered on the x axis in line with their positions in the human genome. The vertical value for each mutation is given by the genomic distance from the previous mutation. The colors of the dots represent different classes of base substitution, as demonstrated at the bottom of the figure. Several hypermutation regions are indicated by the red/green arrowheads. The data are taken from the reference^27^.

**Figure S19.**
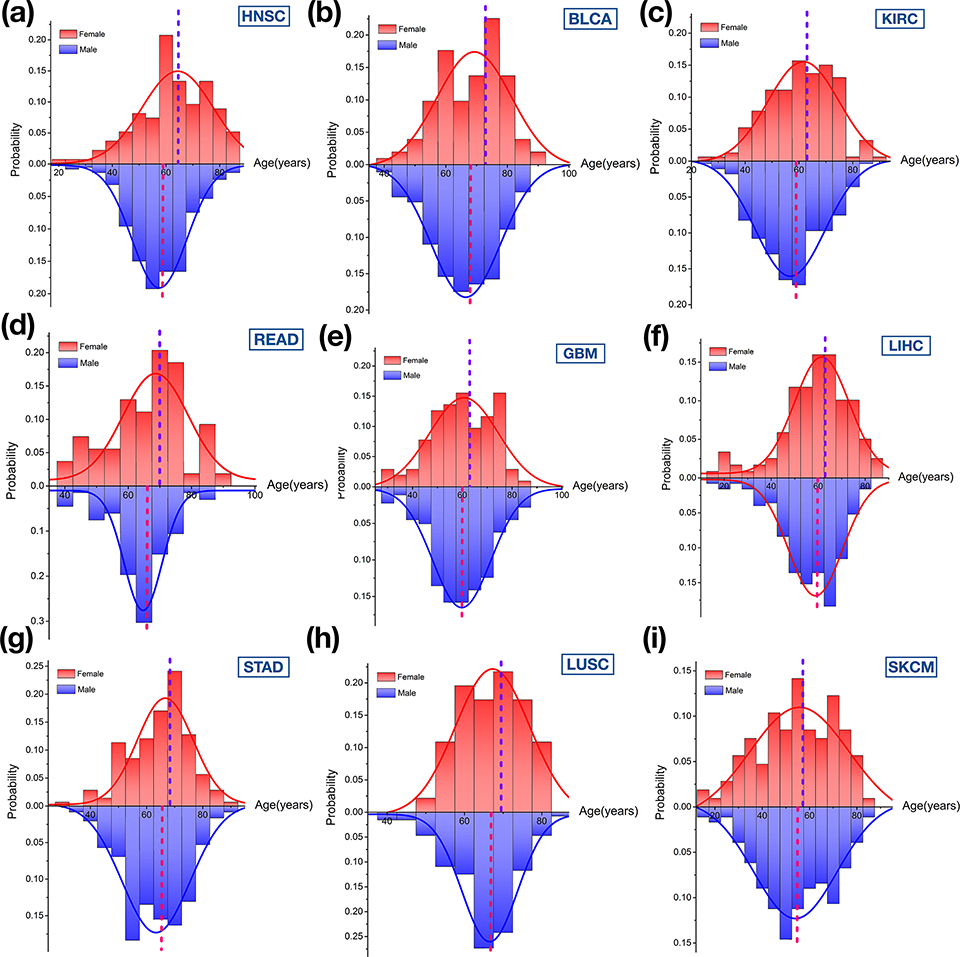
The patient age distribution (PAD) for both the sexes (red for females and blue for males) across the nine cancer types. The solid lines show the Gaussian-function fit of the data. The dashed lines indicate the median age of cancer patients. The median age for female is larger than that for male in this group of cancers. The median ages for female and male patients at diagnosis are listed in Table VI.

**Figure S20.**
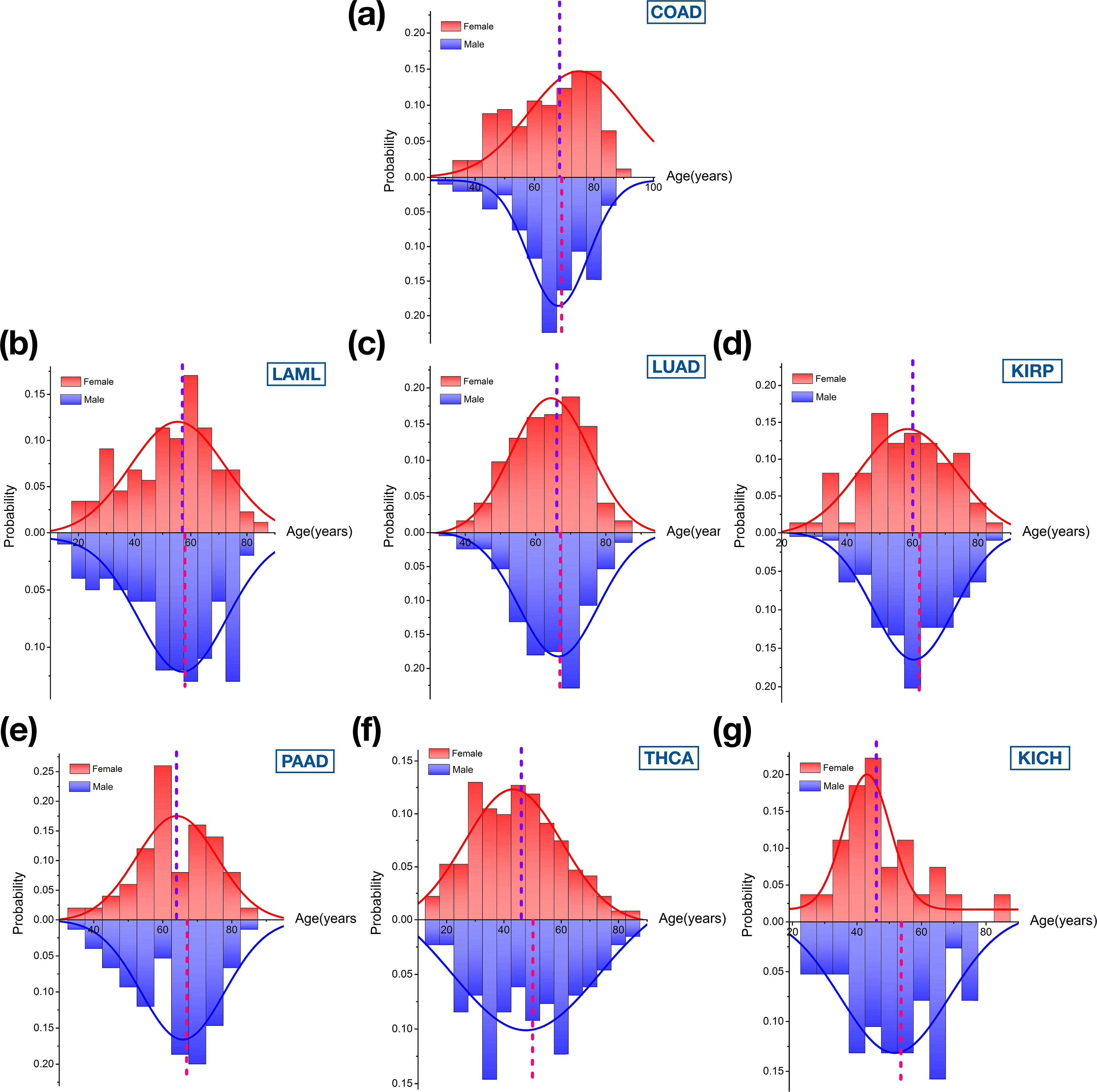
Same as Fig. S19, except we plot the data for different cancer types. Different from Fig. S19, the median male age is typically larger than for females.

**Figure S21.**
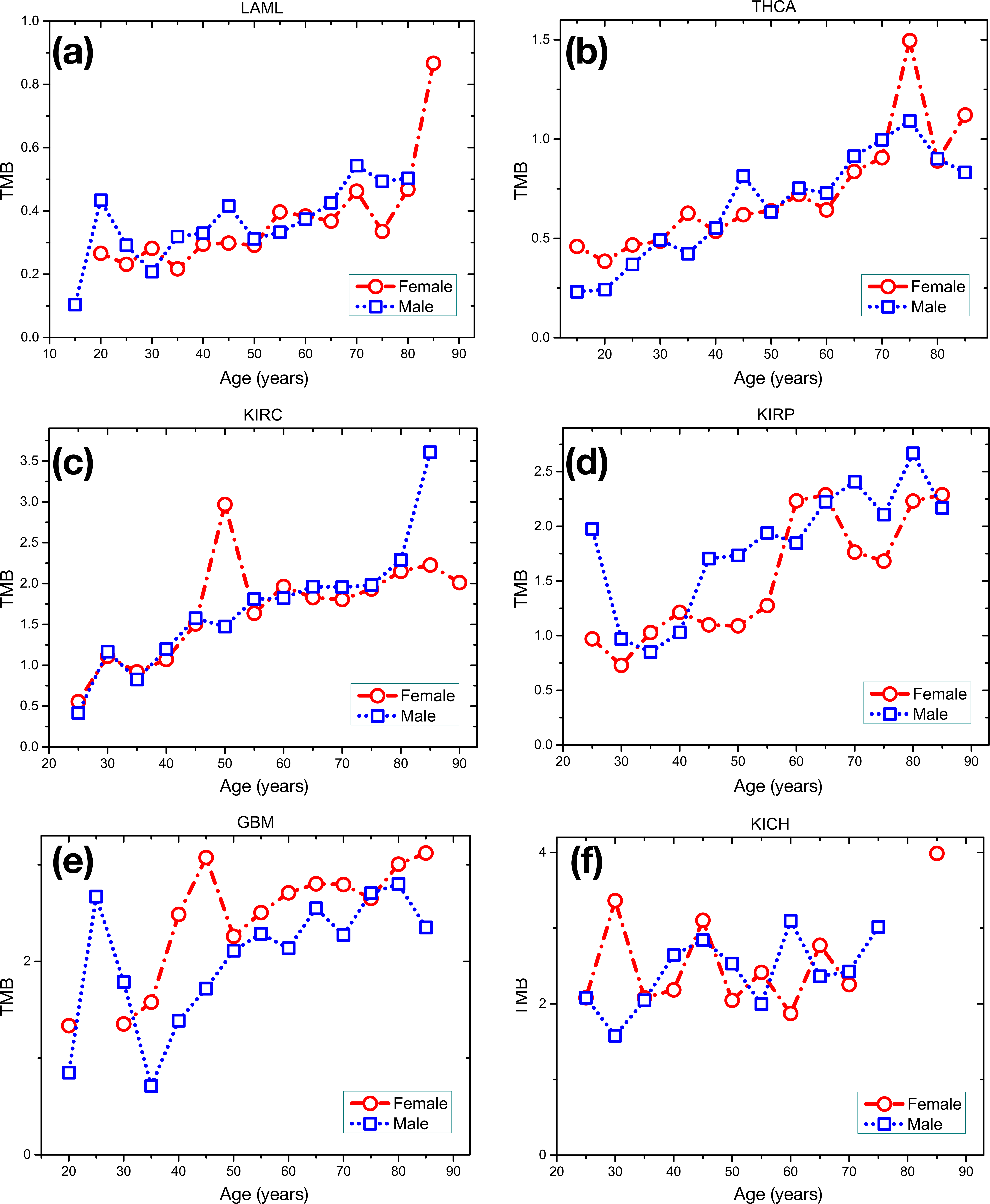
The tumor mutation burden as a function of age for both the sexes with low overall TMB. The mean value for the mutation burden is used for each 5-year period.

**Figure S22.**
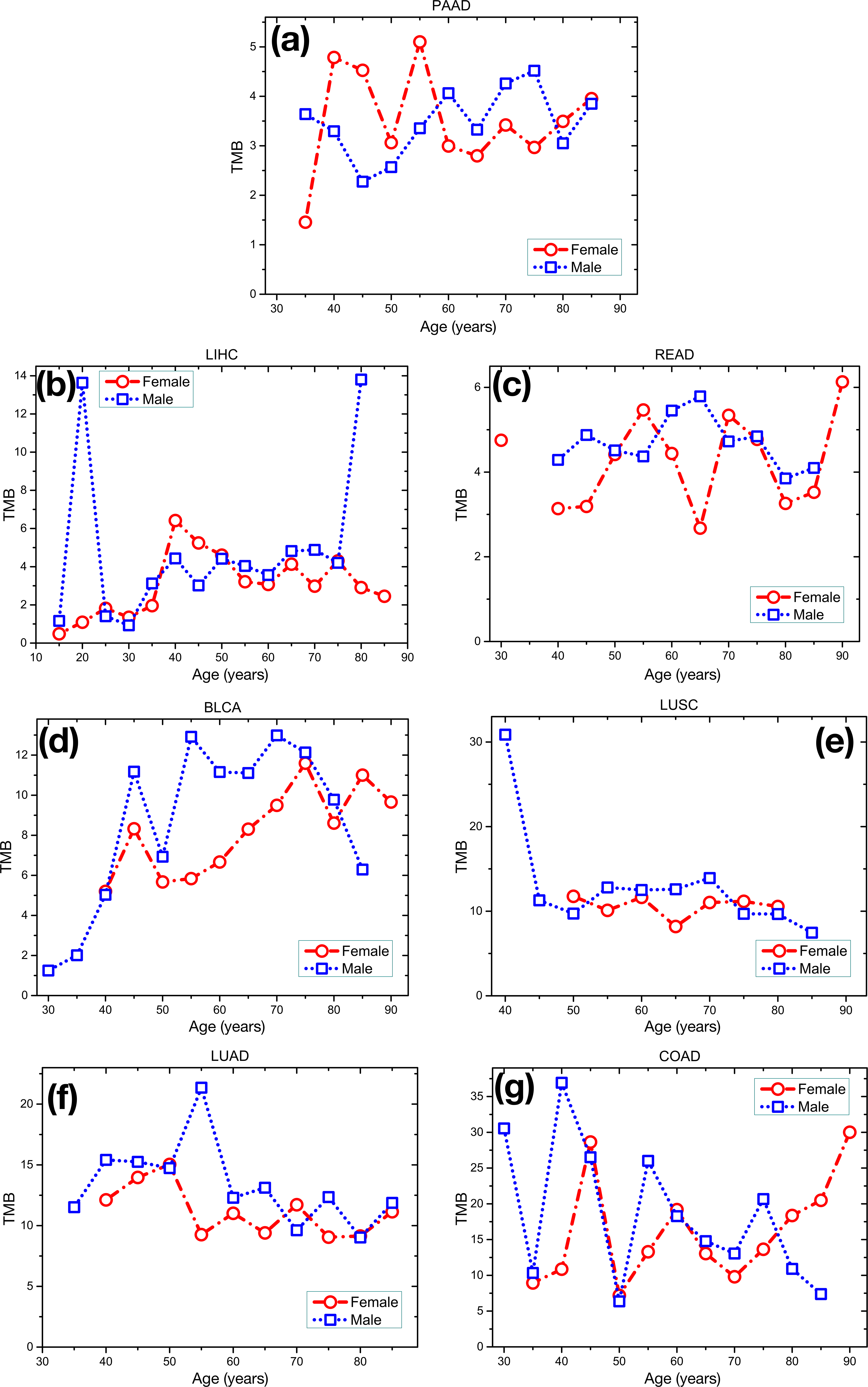
Same as Fig. S21, except we plot the data for cancer types with high overall TMB.

**Figure S23.**
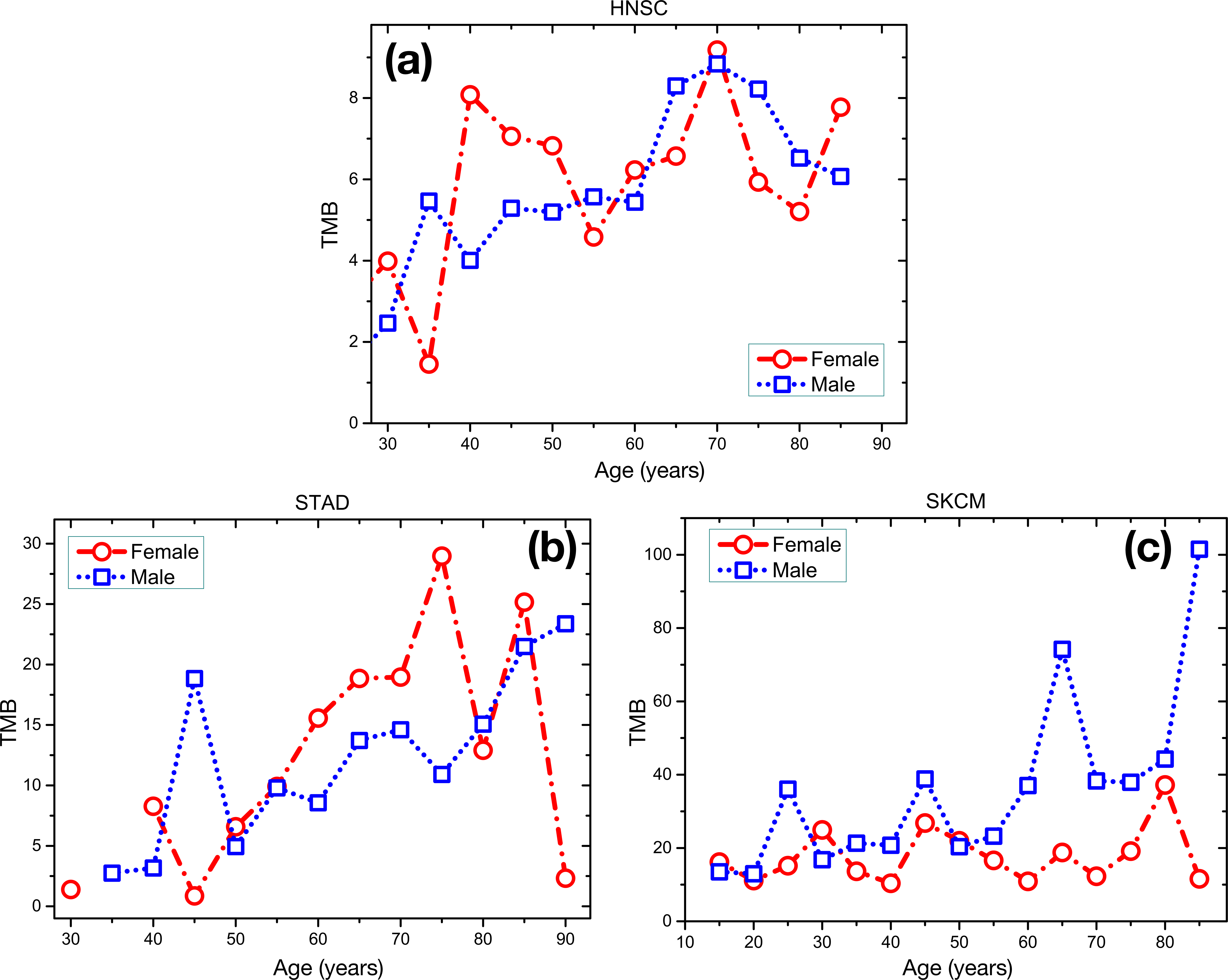
The tumor mutation burden as a function of age for both sexes at high overall mutation burden with strong environmental influence.

**Figure S24.**
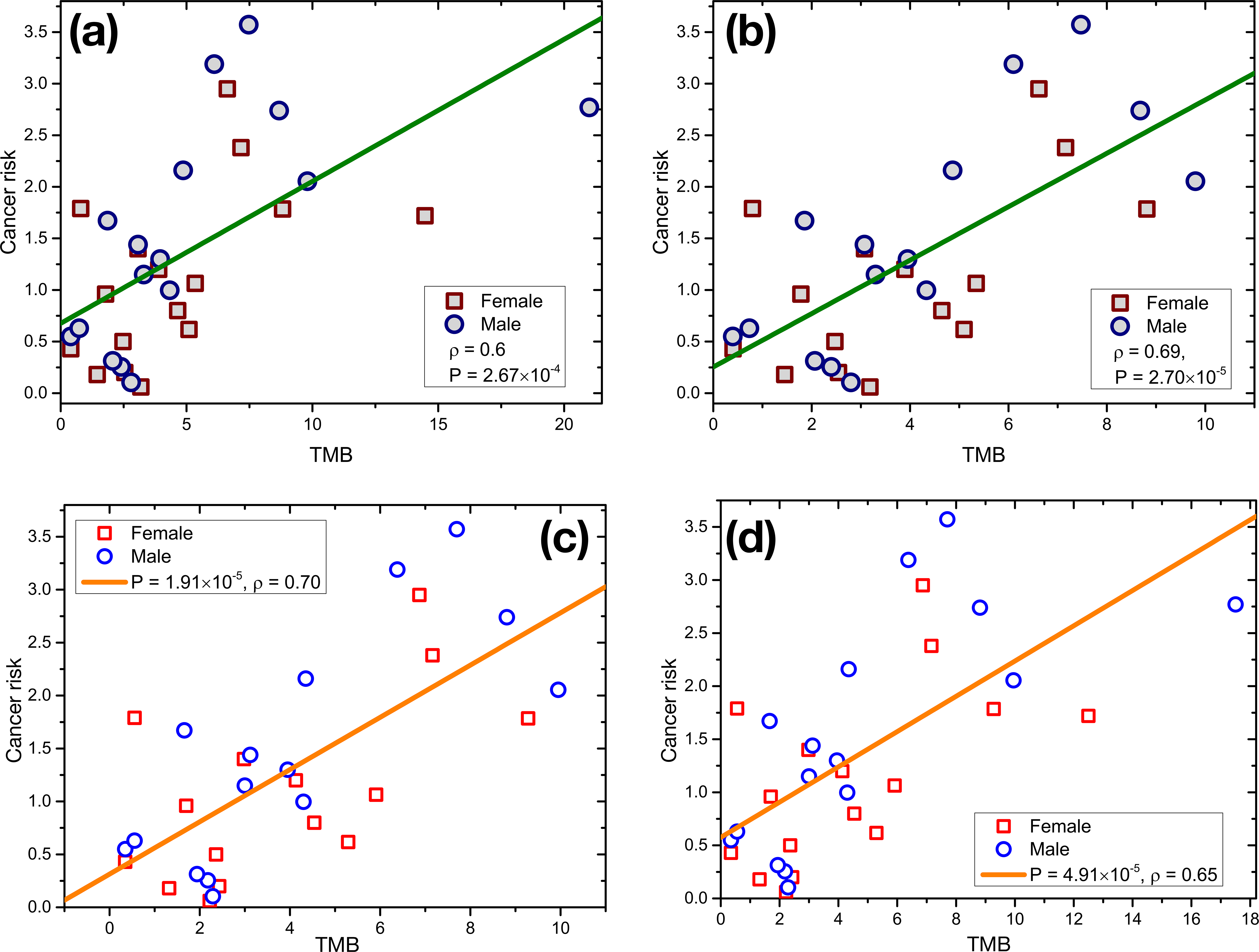
Cancer risk as a function of the TMB across (a), (c) 16 types or (b), (d) 15 types (excluding SKCM) of cancer for both the sexes. The median value for mutations (sum of non-synonymous and synonymous mutations) per megabase (Mb) is used. The solid line gives the regression linear. The P value from an F test, and the Pearson correlation coefficient *ρ* for each cancer type are also shown in the figure. The age-adjusted TMB is used in (a), and (b) but not in (c) and (d).

**Figure S25.**
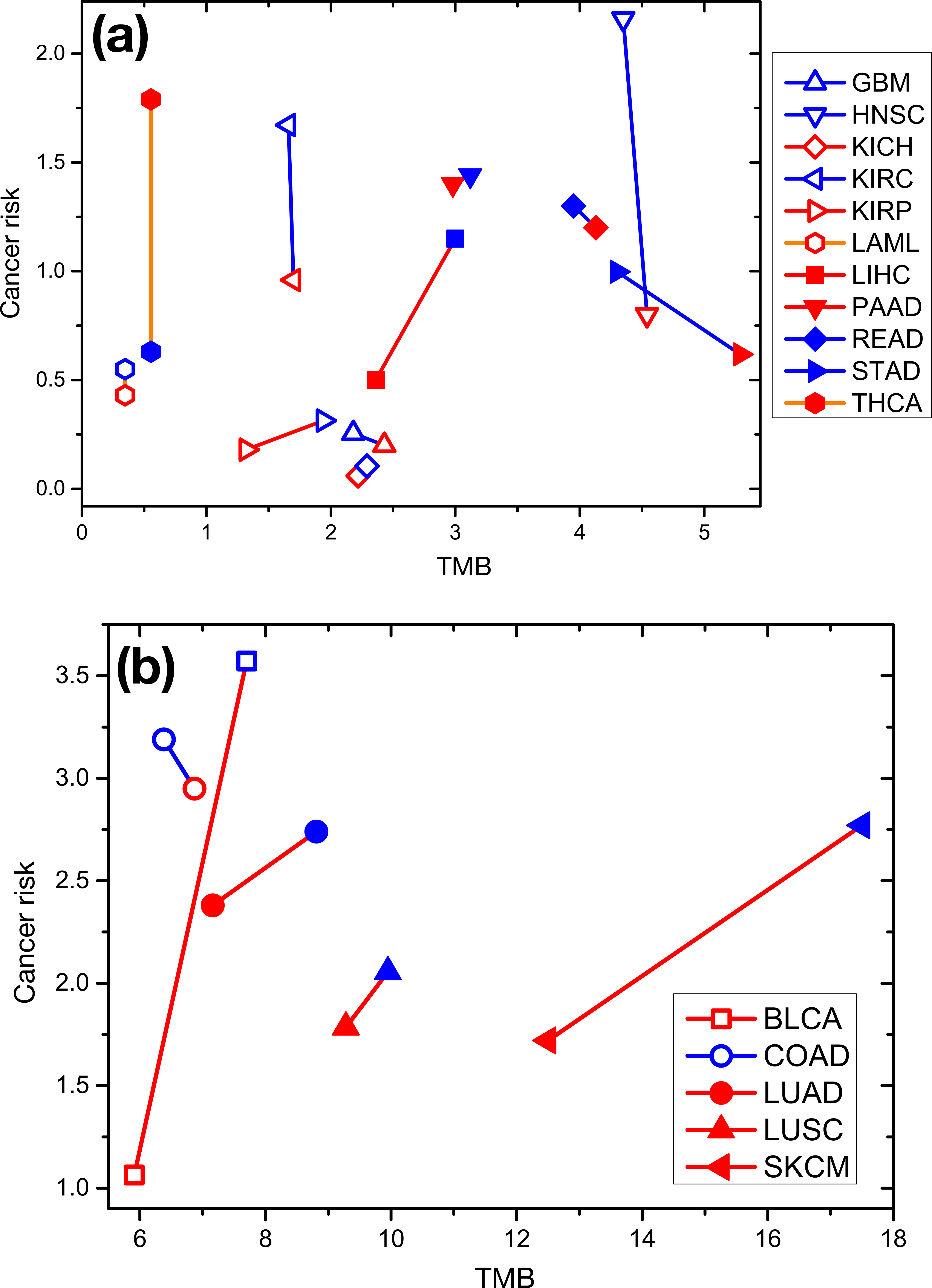
The relationship between cancer risk and mutation burden for both the sexes across 16 types of cancer. The data are the same as in Fig. S24 while the solid lines connect data for both sexes under the same type of cancer. The data are separated into two groups for visualization purpose. Red symbols show the data for females and the data for males are illustrated in blue.

**Figure S26.**
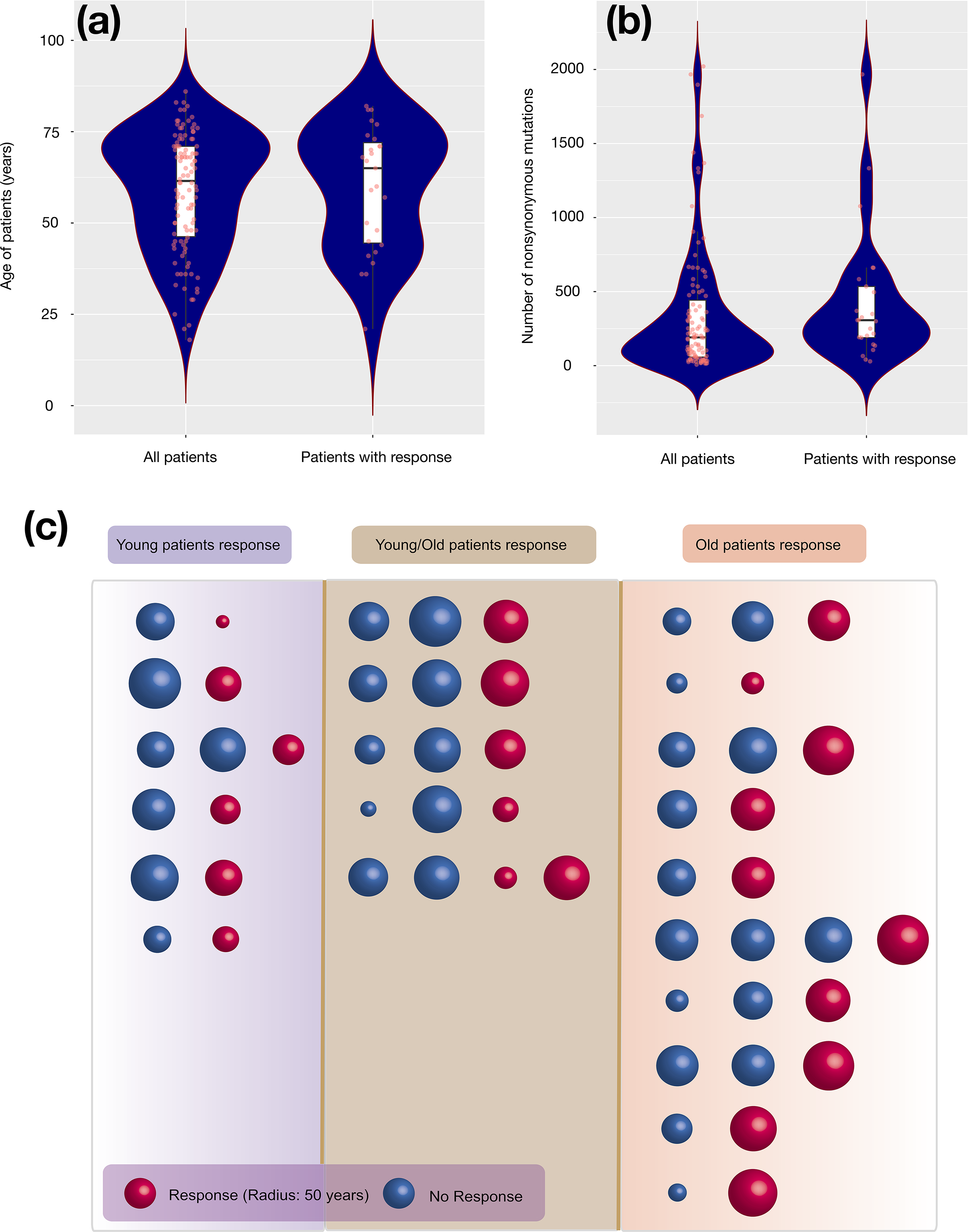
(a) Violin plots for the age distribution of all patients and the patients showing response to the cytotoxic T-lymphocyte-associated antigen 4 (CTLA-4) blockade^79^. (b) Violin plots for the number of non-synonymous mutations for all patients and the patients responding to CTLA-4 blockade. Five data points are not shown in (b) because they are outliers. A box plot is overlaid on each violin plot, and all the data are shown in red dots in the same figure. (c) Schematic of the dependence of response of melanoma patients of differing ages to immunotherapy. The differences in values of the TMB across the rows is within 5%. Red sphere indicates patients who show response to treatment and patients who do not show response are illustrated by blue spheres. The size of the sphere is scaled with patients age as indicated in the lower left corner. The left (right) column gives all pairs in which young (old) patients respond to the treatment. The middle column shows all mixed cases where either young or old patients respond to the treatment.

**Figure S27.**
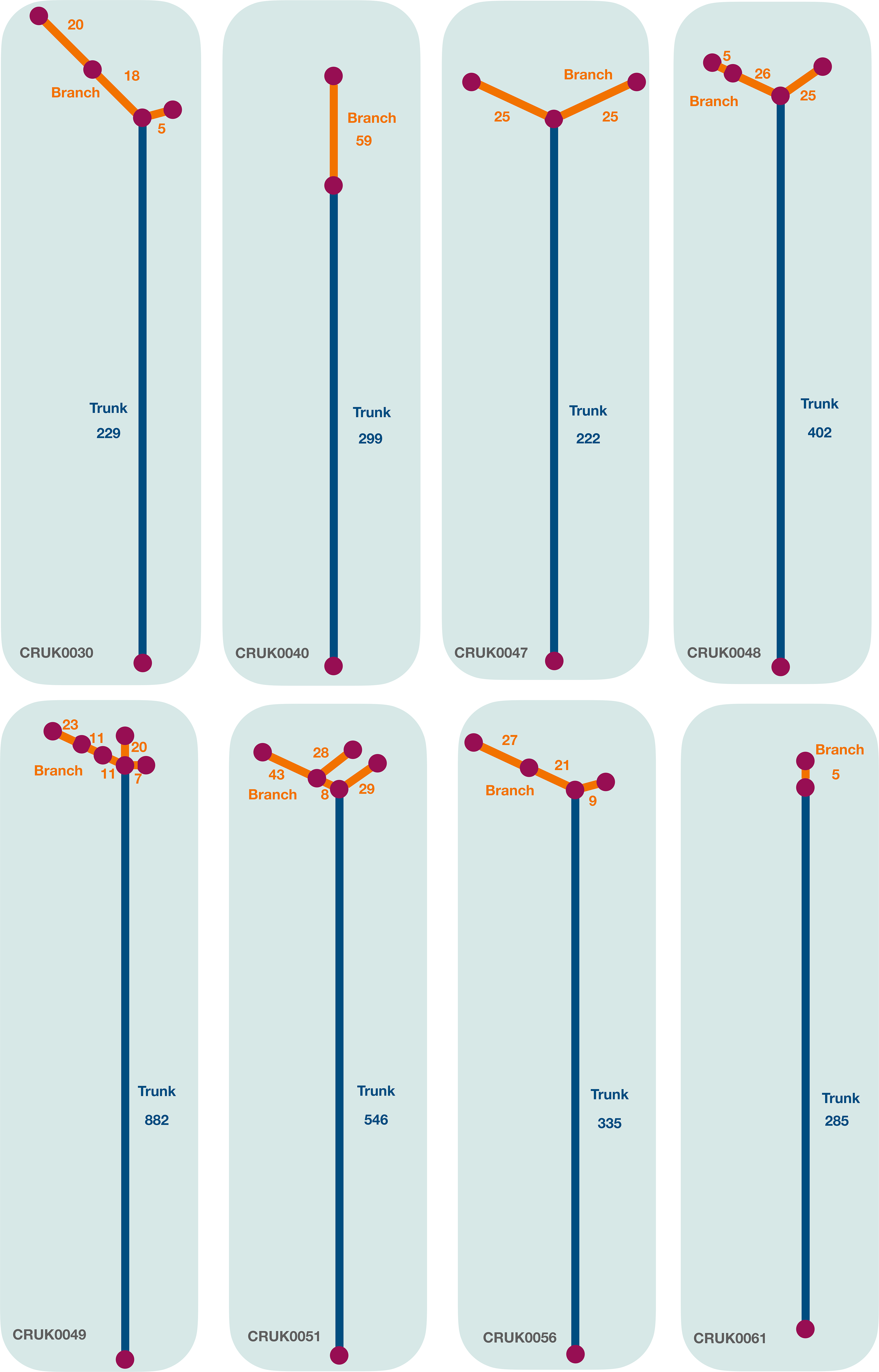
The phylogenetic trees (revised from Ref.^60^) for cancer patients with high cTMB and low ITH discussed in Figure 8(E). The length of the trunk (branches) is proportional to the number of clonal (subclonal) mutations found in the patient which is also listed in the figure accordingly. The trunk of the phylogenetic trees is in navy and the branches are in orange color.

**Figure S28.**
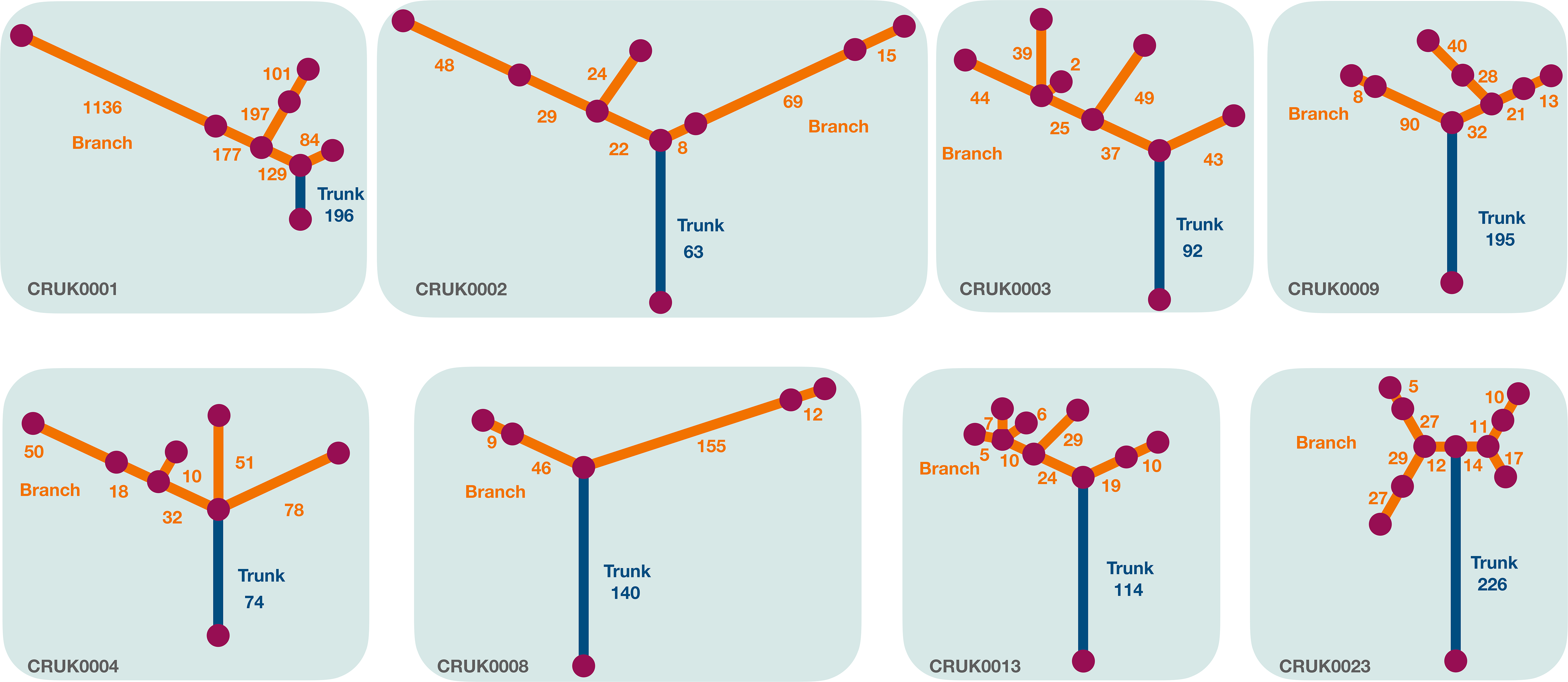
Same as Fig. S27 but for the cancer patients with low cTMB and high ITH discussed in Figure 8(E).

## SOURCE DATA

**Figure 1 source data 1-6** The tumor mutation burden data from whole-exome sequencing and patient clinical information (gender, age) for cancer types in Fig. 1 are included in the data file.

**Figure 2 source data 1-7** The tumor mutation burden data from whole-exome sequencing and patient clinical information (gender, age) for cancer types in Fig. 2 are included in the data file.

**Figure 3 source data 1-3** The tumor mutation burden data from whole-exome sequencing and patient clinical information (gender, age) for cancer types in Fig. 3 are included in the data file.

**Figure 4 source data 1** The tumor mutation burden data from whole-genome sequencing and patient clinical information (gender, age) for all 24 cancer types in Fig. 4 are included in the data file.

**Figure 6 source data 1-2** All single nucleotide variants from whole-exome sequencing for LAML (197 patients) and LIHC (198 patients) are included in the data file.

**Figure 8 source data 1** The (sub)clonal mutations from multi-region sequencing for 61 lung adenocarcinoma patients are included in the data file.

**Figure S18 source data 1** All single nucleotide variants from whole-genome sequencing for LAML, LIHC, and ESAD cancer patients are included in the data file.

